# Transposable element–host genome evolutionary arms race revealed by multi-modal epigenomic profiling in a telomere-to-telomere human genome reference

**DOI:** 10.64898/2026.03.19.712972

**Authors:** Daniil Nikitin

## Abstract

For a quarter of a century transposable elements have been recognized as a major component of the human genome, comprising 46.1% according to recent estimates, and as key drivers of regulatory innovation as well as participants in an ongoing evolutionary arms race with host defense systems. Using the newly released T2T ENCODE dataset, we quantified the epigenetic impact of 3.7 million transposable elements across evolutionary time by analyzing seven epigenomic modalities in twelve human cell lines, spanning six transposon classes, 44 families, and 1,122 subfamilies. We show that SVA elements exhibit the strongest signatures of the arms race, characterized by progressive escape from H3K9me3-mediated heterochromatinization accompanied by increased acquisition of CTCF binding and enhancer-associated chromatin marks. Among Alu elements, the AluYb8 and AluYb9 subfamilies display age-dependent accumulation of CTCF binding, while seven LTR subfamilies (HERV16-int, MER11C, LTR43-int, HERVE-int, LTR22C, LTR5_Hs, HERVIP10FH-int) demonstrate dynamic evolutionary behavior within active chromatin, H3K9me3 chromatin and CTCF contexts. We further evaluated the relative contribution of distinct epigenomic modalities to the host–transposable element conflict and found that transposon-driven evolution is dominated by evasion of host-imposed heterochromatinization primarily at H3K9me3 and secondarily at H3K27me3, together with progressive invasion into CTCF-rich regions. In contrast, enhancer, promoter, and H3K36me3 marks appear to play more limited roles. Collectively, these findings deepen our insight into the coevolutionary epigenomic dynamics between human genome and transposable elements and the associated processes driving regulatory innovation.

## 1. INTRODUCTION

According to the complete telomere-to-telomere (T2T) human genome assembly (Nurk *et al*. 2022) transposable elements (TEs) occupy approximately 46.1% of the human genome. Of this total, 3.6% corresponds to DNA transposons, while the remaining 42.5% consists of retroelements (REs), which propagate through RNA intermediates via reverse transcription. The interaction between TEs and the host genome represents a long-standing evolutionary arms race, in which newly mobilized elements are counteracted by a variety of genomic defense mechanisms, including microRNAs (Tristán-Ramos *et al*. 2020), piwi-interacting RNAs (Rosenkranz, Zischler, and Gebert 2022), heterochromatinization (Stamidis and Żylicz 2023), human silencing hub (Lehner 2025), Toll-like receptors (Gázquez-Gutiérrez *et al*. 2021) and adenosine deamination (Modenini, Abondio, and Boattini 2022a).

A particularly striking example is the KRAB-ZNF transcription factor family, the largest in tetrapods (Yang P, Wang, and Macfarlan 2017a), which has evolved under strong positive selection to repress emerging TE families. This family underwent rapid lineage-specific expansion in primates, producing hundreds of primate-specific KRAB-ZNF genes (Jacobs *et al*. 2014). Perturbation of this TE suppression network has been implicated in several human diseases, including neurodegenerative disorders (Chen, Maupas, and Nowick 2025).

TEs have also played pivotal roles in major evolutionary transitions within the human evolutionary lineage. They have been proposed as drivers to eukaryogenesis (Vosseberg and Snel 2017), the origin of adaptive immunity (Yakovenko *et al*. 2021) and the evolution of the invasive placenta (Carter 2021). Other milestones—such as the emergence of multicellularity (Johnson 2008) and the expansion of the human neocortex (Garza *et al*. 2023)—were also significantly influenced by TEs, albeit less directly.

In the biomedical context, TEs are increasingly recognized as contributors to human disease, including neurodegeneration (Ravel-Godreuil *et al*. 2021), autism spectrum disorders (Techaniyom *et al*. 2025) and cancer (Chaaban *et al*. 2025). Their involvement in oncogenesis is particularly important, since TE-derived nucleic acids and tumor-associated neoantigens can modulate tumor immunity. Furthermore, both reverse transcriptase inhibitors and epigenetic modulators—targeting the retrotransposition process—are being explored as novel therapeutic strategies in oncology (Wang *et al*. 2024).

The functional characterization of human TEs has been an area of intense research and has been reviewed elsewhere, encompassing studies of TE-derived transcripts (Lanciano and Cristofari 2020, Lim, An, and Park 2025), their epigenetic landscapes (Stefano Di 2022), and their cis-regulatory effects on neighboring genes (Ali, Han, and Liang 2021). The evolutionary dynamics of TEs have also been extensively explored, including analyses of primate-specific families, population variation, and the characteristic “boom-and-bust” cycles of TE activity (Bourgeois and Boissinot 2019). However, studies explicitly linking TE evolution with functional regulatory consequences—particularly in the context of transcriptional and epigenetic modulation—remain comparatively limited.

Recent large-scale datasets have begun to bridge this gap. For example, analyses of the ENCODE4 cell lines dataset revealed that approximately 25% of human candidate cis-regulatory elements are TE-derived (Du *et al*. 2024). Similarly, an earlier study of normal tissues from the Roadmap Epigenomics Project estimated that about one quarter of the human regulatory epigenome is TE-associated, 47% of human TEs can attain the active regulatory state (Pehrsson *et al*. 2019).

Our research consortium has systematically investigated these relationships over the past decade. We first characterized transcription factor binding site (TFBS) profiles across human LTR families and subfamilies, examining their variation with divergence from consensus sequences as a measure of evolutionary age (Garazha *et al*. 2015). Building on these findings, we developed RetroSpect, a computational framework to quantitatively assess the regulatory impact of REs on host gene expression and pathway activation (Nikitin *et al*. 2018). Applying this approach to ENCODE data encompassing 563 transcription factors and six chromatin modifications across multiple cell lines (Nikitin *et al*. 2018, 2019, 2019, Nikitin, Garazha *et al*. 2019b, 2019a), we identified key biological processes enriched in RE-associated regulatory inputs. These included miRNA-mediated gene regulation (Zottel *et al*. 2020), olfactory and color vision pathways, fertilization mechanisms, immune responses, and metabolic processes related to amino acid, fatty acid, and xenobiotic metabolism—indicating accelerated rates of regulatory evolution in these systems (Nikitin, Garazha *et al*. 2019b).

Subsequent analyses integrated active (H3K4me1, H3K4me3, H3K9ac, H3K27ac) and repressive (H3K27me3, H3K9me3) histone marks, confirming that RE-enriched and RE-depleted processes exhibit coordinated patterns of regulatory evolution depending on chromatin state and corroborating earlier findings of RE-activated processes from the TFBS analysis (Nikitin *et al*. 2019).

More recently, an integrative analysis across five cell lines (Nikitin 2025) extended RetroSpect to identify pathways co-marked by RE-linked promoter (H3K4me3, H3K9ac) and enhancer (H3K4me1, H3K27ac) activity, while depleted of heterochromatin marks. This consensus-based approach revealed that RE-linked promoter marks are preferentially associated with oncogenic and immune pathways, including those implicated in chronic myeloid leukemia, small-cell lung cancer, and innate immune responses to HTLV-1 infection.

However, all previous analyses were constrained by reliance on older human genome assemblies (hg19, hg38). The advent of the T2T complete genome assembly (Nurk *et al*. 2022) has significantly improved TE resolution, identifying approximately 1% additional Alu and L1 elements and up to 1.2% more members of major HERV families. Notably, 118,787 out of 4,531,994 TEs (2.6%) had their genomic coordinates refined, and the assembly revealed the tendency of fully active REs to reside within centromeric high-order repeat structures, retaining transcriptional competence.

These advances underscore the need for a comprehensive, T2T-based functional and evolutionary analysis of human TEs, encompassing both their evolutionary trajectories and epigenomic profiles. In the present study, leveraging the recently released T2T ENCODE dataset, we examined the epigenetic dynamics of 3.7 million TEs (including 0.46 million DNA transposons) across 12 human cell lines over evolutionary time. Using seven epigenomic marks (CTCF, and the histone modifications H3K4me1, H3K4me3, H3K9ac, H3K27ac, H3K27me3, H3K9me3), we characterized six major TE classes—LINE, SINE, LTR, SVA, Helitron, and DNA elements—along with 48 TE families and 1139 classified subfamilies. This study builds upon our earlier RetroSpect-based analyses of TE regulatory evolution (Garazha *et al*. 2015, Nikitin *et al*. 2019), extending them to a T2T-resolved, epigenome-wide evolutionary framework.

Our analysis reveals that SVA elements represent the most epigenetically dynamic and evolutionarily recent TE class, characterized by progressive evasion of heterochromatinization, accumulation of enhancer-associated H3K4me1 mark and CTCF binding. Moreover, their peak (mode) divergence is the lowest one (6%) among the other classes. Two Alu subfamilies (AluYb8 and AluYb9) and seven LTR ones (HERV16-int, MER11C, LTR43-int, HERVE-int, LTR22C, LTR5_Hs, HERVIP10FH-int) were the most dynamic.

Moreover, we present the new evidence of the relative contributions of seven epigenomic modalities to the evolutionary arms race between TEs and human genomes. Our results indicate that TEs preferentially evade defensive heterochromatinization primarily associated with H3K9me3, and to a lesser extent H3K27me3, while preferentially integrating within chromatin architectural regions marked by CTCF as a possible evolutionary strategy. In contrast, enhancer-associated (H3K27ac, H3K4me1), promoter-associated (H3K4me3), and transcribed gene body associated (H3K36me3) dimensions of TE–host co-evolution appear to be less influential overall or largely cell-type specific. Collectively, these findings advance our understanding of TE-driven regulatory innovation in human evolution.

## 2. RESULTS

### 2.1. Transposon classes and their divergence

As the first stage of our analysis, we characterized the age distribution of all TEs in the human genome (Figure 1B) as well as of individual TE classes (Figure 1C). The length-weighted average Kimura 2 parameters (K2P) CpG adjusted divergence from the consensus sequence was used as a proxy for evolutionary age (Giordano *et al*. 2007).

**Figure 1.**
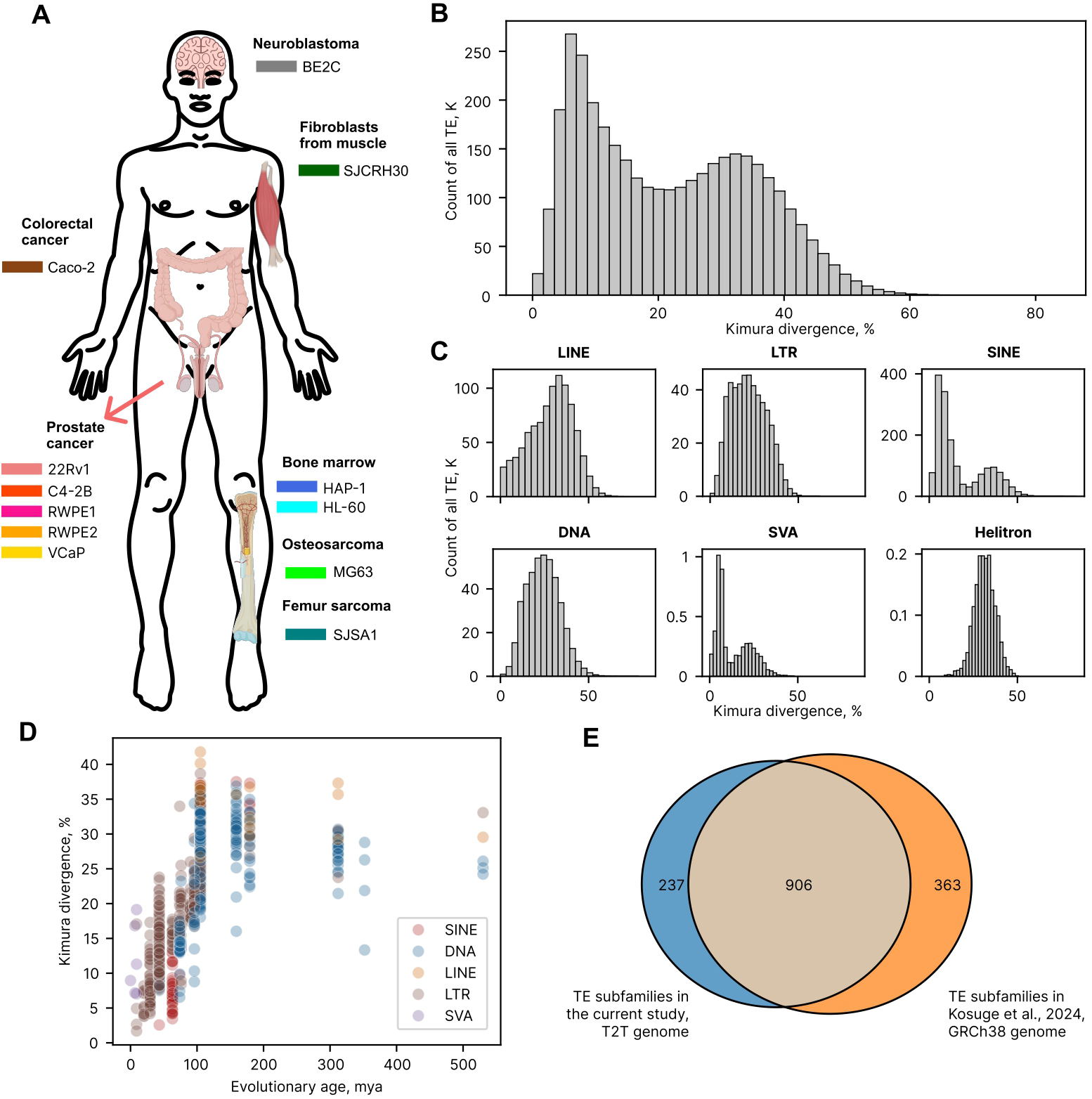
Overview of the analyzed cell lines and TEs. (A) Schematic representation of the twelve human cell lines and their corresponding organs or tissues of origin, compiled from the ATCC database (ATCC: The Global Bioresource Center | ATCC, n.d.) and illustrated using BioRender. The color scheme assigned to each cell line is used consistently throughout subsequent figures in this study. (B) Distribution of Kimura 2 parameter (K2P) divergence scores across all TEs examined, measured in percents. (C) Divergence score distributions by TE class, shown for 1,660,854 SINEs, 949,629 LINEs, 499,814 LTRs, 434,591 DNA transposons, 4,972 SVAs, and 1,710 Helitrons. (D) Correlation of average divergence by TE subfamily and evolutionary age estimations from (Kosuge, Ito, and Hamada 2024). (E) Venn diagram showing intersection between TE subfamilies in (Kosuge, Ito, and Hamada 2024) and in the current study.

The overall distribution of all annotated elements displayed a bimodal pattern, with peaks at approximately 6–7 and 31–32% divergence. These two divergence modes likely represent two major waves of TE activity, roughly aligning with (i) the divergence of Old and New World monkeys (∼35–40 Mya) and (ii) the early radiation of mammalian lineages (∼150 Mya), consistent with previous defragmentation-based age estimates (Giordano *et al*. 2007, Craig, Hedges, and Kumar 2024).

When analyzed by TE class, LINE, LTR, DNA, and Helitron elements each exhibited unimodal divergence distributions, suggesting a single dominant burst of amplification, with approximate peak K2P divergence values of 33%, 24%, 25%, and 31%, respectively (Figure 1B). In contrast, SINE and SVA elements showed bimodal distributions, reflecting two distinct proliferation episodes centered at approximately 6–7% and 36–37% (SINEs) and 6% and 24% (SVAs).

To further validate T2T–based K2P divergence as a proxy for evolutionary age, we leveraged TE age estimates from a recent KRAB-ZNF and TEs evolutionary arms race study (Kosuge, Ito, and Hamada 2024) based on the GRCh38 assembly and compared these estimates with divergence scores at the level of individual TE subfamilies (Figure 1D). Up to approximately 110 million years ago (mya) and divergence values of 25–30%, evolutionary age and divergence exhibited a strong linear relationship (Pearson *r* = 0.831, *p* = 4.8 * 10^-193^), whereas older subfamilies converged at divergence values of ∼30% largely independent of their inferred age. Furthermore, of the 1,122 TE subfamilies analyzed in the present study, 906 (79.3%) overlapped with the subfamilies reported by (Kosuge, Ito, and Hamada 2024) (Figure 1E). Collectively, these results support the use of average divergence as a reliable proxy for the evolutionary age of TE families and subfamilies up to 110 mya with the current data.

Additionally, we investigated a connection between the CpG adjusted K2P divergence and other metrics of evolutionary age that are available in UCSC Genome browser or in the complete RepeatMasker output for the T2T genome: average divergence score per 1000 base pairs according to UCSC genome browser track, percent divergence by RepeatMasker, ratio of transitions to transversions and CpG unadjusted Kimura divergence (Supplementary Figure 1A). For the repeats consisting of individual parts called by RepeatMasker (such as 3’ end, 5’ end and central ORF for LINE elements), length-weigted average was taken for all te metrics. As expected, average divergence by UCSC Genome browser linearly correlated with percent divergence by RepeatMasker (with a slope 1), whereas Kimura divergence estimates (both CpG-adjusted and unadjusted) formed a non-linear curve: at the Kimura divergence exceeding 25%, average divergence by UCSC Genome browser reached approximately 200-210 substitutions per 1000 bp, indicating an impact of multiple substitutions per single bp. Moreover, CpG adjusted K2P divergence showed two separate curves for SINEs and LINEs, with LINEs showing less average divergence under the same values of CpG adjusted K2P divergence, which was not seen for the unadjusted K2P divergence (Supplementary Figure 1A).

### 2.2. Dependency between epigenetic activity and evolutionary age of TEs at the level of TE classes

To assess how epigenetic activity and regulatory potential vary across TE classes as a function of evolutionary age, we analyzed correlations between ChIP–seq enrichment signals, taken individually across all available cell lines and histone modifications or CTCF, and TE divergence scores separately for each of the six major TE classes (Figure 2). Overall, Spearman correlation coefficients spanned a range from −0.502 to 0.459 and were centered near zero, indicating that, at the class level, epigenetic activity shows predominantly weak to moderate dependence on evolutionary age (Figure 2A, Supplementary File 1).

**Figure 2.**
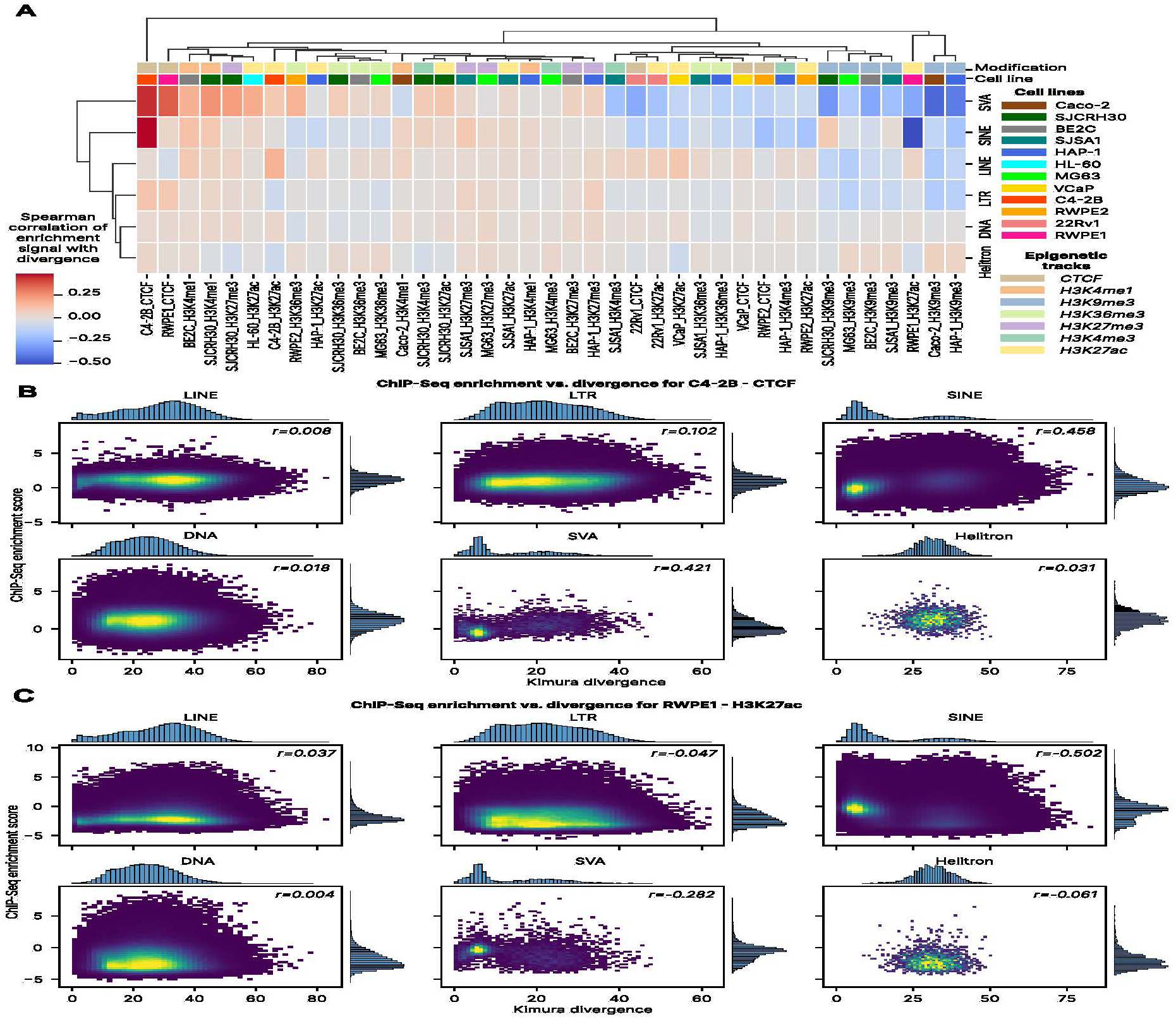
Correlation between epigenetic enrichment and evolutionary divergence of human TEs at the level of classes. (A) Clustermap of Spearman correlation coefficients between ChIP-seq enrichment signals and divergence scores across TE classes, cell lines, and epigenomic modalities. (B) Correlation between CTCF ChIP-seq enrichment in the prostate-derived cell line C4-2B and TE divergence scores, shown separately for six TE classes: LINEs, LTRs, SINEs, DNA transposons, SVAs, and Helitrons. (C) Corresponding correlation between H3K27ac ChIP-seq enrichment in another prostate cell line, RWPE1, and divergence scores for the same six TE classes.

The heterochromatin-associated histone mark H3K9me3 exhibited a consistent clustering pattern across 4 cell lines (SJSA1, MG63, BE2C, SJCRH30: two sarcoma cell lines, fibroblasts and neuroblastoma), showing notably negative correlations with divergence for SVA elements (−0.161 to −0.326). This pattern likely reflects a shared epigenetic mechanism of constitutive heterochromatinization that weakens with the evolutionary age of SVAs. Interestingly, CTCF in RWPE1, H3K4me1 in BE2C and SJCRH30, H3K27me3 in SJCRH30 and H3K27ac in HL-60 formed a distinct cluster, showing weakly positive correlations within the SVA class (Spearman *r* = 0.155–0.343). These findings suggest that subsets of SVA elements can adopt active enhancer or chromatin architectural roles, or undergo conditional repression via Polycomb-mediated mechanisms increasingly over evolutionary time.

Apart from CTCF in C4-2B and H3K27ac in RWPE1 (Figure 2B, C), SINE elements displayed negligible correlations with divergence across all cell lines and histone marks (Spearman *r* = −0.208 to 0.107). A similar absence of correlation was observed for LINEs, LTRs, Helitrons, and DNA transposons, with Spearman *r* ranging from −0.168 to 0.152, −0.159 to 0.102, −0.078 to 0.047, and −0.053 to 0.031, respectively. Notably, Helitrons, having the lowest number of members (1,710), were co-clustered with DNA elements (434,591 copies) which is an indirect argument against the outlying clustering of SVA elements being explained by their low number (4,972), allowing high random fluctuations of correlations.

In the overall clustermap (Figure 2A), CTCF in C4-2B and H3K27ac in RWPE1 appeared as distinct outgroups. CTCF in C4-2B displayed positive correlations with divergence for both SINEs and SVAs (Figure 2B), suggesting that the architectural potential of these elements may increase with evolutionary age. While the role of Alu elements in CTCF-mediated chromatin organization is well established (Dixon *et al*. 2012), a similar trend has not been convincingly demonstrated for SVAs except for the BORIS paralog (Pugacheva *et al*. 2016). However, as this pattern was not reproduced in other prostate or non-prostate cell lines, its broader biological significance remains uncertain. Conversely, H3K27ac in RWPE1 (Figure 2C) displayed a negative correlation with divergence in SINEs and SVAs, indicating that enhancer-associated activity diminishes in evolutionarily older repeats (K2P divergence scores 20–50; enrichment scores −2 to −4). Again, this trend was not replicated across other cell lines.

Overall, the distribution of Fisher *z*-transformed Pearson correlation coefficients was highly non-normal (Shapiro–Wilk *p =* 2.0*10^-15^), symmetric, and strongly leptokurtic (kurtosis = 7.99, p-value = 8.8*10^-12^), indicating the divergent evolutionary patterns. The largest absolute correlations and the highest variance were observed for SVA elements (Levene test *p* = 5.9*10^-14^ compared to all other classes; multi-class Levene’s *p* = 3.7*10^-16^). This finding may indicate that SVAs are engaged in the most rapid evolutionary arms race with host defense systems or that their regulatory influence is changing more dynamically than in other TE classes. Notably, LINEs, SINEs, and LTRs are 100–200 times more abundant than SVAs in the human genome, suggesting that individual families within these numerically dominant classes may nevertheless harbor similarly pronounced regulatory dynamics that are masked at the aggregate class level.

We performed additional background checks to investigate whether the observed patterns are non-random or not a by-product of other dependencies. Keeping in mind the fact that ChIP-seq reads mappability could depend on TE length (Sexton and Han 2019), we correlated TE length (according to UCSC Genome browser track) with their Kimura divergence by classes (Supplementary Figure 1B). SINEs showed the strongest negative correlation of length with divergence (Spearman r = -0.520), with LINEs being the second one (Spearman r = -0.480). Both groups had distinct bimodal shapes with an older peak having lower length. Nevertheless, SVA elements, which demonstrated the strongest and the most variable correlations of different epigenomic modalities in different cell lines, had relatively low absolute value of length correlation with divergence (Spearman r = -0.295). Therefore, differential mappability of ChIP-seq reads on variable TE lengthes could not explain a half of the observed patterns.

Moreover, since evolutionary age correlates with Kimura divergence only up to aproximately 110 mya and 20-25% divergence (Figure 1D), we studied the robustness of the observed correlations and their clustering when they were calculated under the divergence cut-off at the level of 20%, 22.5% and 25% (Supplementary Figure 2A, 2B, 2C respectively). In all the clustering attempts, all the cell lines where epigenomic profiles of the constitutive heterochromatin marker H3K9me3 were available, showed co-clustering with weakly negative correlations in LINE, LTR and SVA elements, which indicates a robust evolutionary dependency. CTCF in C4-2B was always the most basal branch, whereas H3K27ac in RWPE1 was second or third most basal branch, exchanging with H3K27ac in C4-2B. Clustering pattern of TE classes was highly unstable.

### 2.3. Dependency between epigenetic activity and evolutionary age of TEs at the family level

To investigate potential correlations between epigenetic activity and evolutionary age at the level of individual TE families, we analyzed all 44 families annotated in the RepeatMasker database. For each family, Spearman correlation coefficients were calculated between ChIP–seq enrichment signals and K2P divergence scores across the full set of 39 cell line–epigenetic mark pairs (Supplementary File 1). The resulting clustermap, together with family-specific TE counts and mean divergence scores, is presented in Supplementary Figure 3 and provides a framework for identifying TE families actively engaged in evolutionary arms races with host defense mechanisms. Examination of the clustermap revealed that the majority of TE families exhibit low absolute correlation values (<0.2, 95.9%), suggesting that these families are largely evolutionarily quiescent and do not display evidence of mutational evasion from host-imposed epigenetic repression or of progressive acquisition of activating chromatin modifications over time.

As in the case of TE classes, we did the analogous clustering of the correlations that were calculated based on the cut-off at the level of 20%, 22.5% and 25% K2P divergence (Supplementary Figures 4A, 4B, 5A, respectively). The overall structure of the cut-off clustermaps was similar, with the majority of families having near-zero correlations and the maximum/minimum ones reaching 0.8 correlation by their absolute value. We compared the original clustermap dendrogram (Supplementary Figure 3) with the derived ones using Spearman cophenetic correlation, and plotted distribution of such cophenetic correlations for 10000 random permutations of these dendrograms (Supplementary Figure 5B, 5C and 5D for cut-off levels of 20%, 22.5% and 25%, respectively). All the comparisons showed that the observed differences are within a range of neutral expectation (empirical percentile is between 5% and 95%). Therefore, the dendrograms of correlation coefficients for the original and the cut-off correlations could be interpeted as statistically indistinguishable.

Interestingly, TE families with low copy numbers tended to occupy peripheral branches of the clustermap and displayed relatively higher correlation coefficients (Supplementary Figure 3). To quantify this relationship, we compared total TE counts per family with the corresponding average absolute Spearman correlation, using the latter as a proxy for the strength of the evolutionary arms race. This analysis revealed a nonlinear, decreasing relationship, well-approximated by a sigmoidal curve (fitted via nonlinear least squares, R^2^=0.575, p=2.5*10^-9^, Figure 3A). Notably, average correlations approached negligible values (<0.05) for families exceeding 500 copies. To assess whether correlations in low-copy families might be inflated by random variation, we compared the observed sigmoidal fit against a background dataset of 100 random shufflings of divergence and enrichment values for each family (Supplementary Figure 6A) using an F-test (p=1.9*10^-34^; Supplementary File 1). These observations suggest that small TE families may undergo active, successive evasion of host defenses, whereas larger families are subject to stronger selective constraints and more extensive epigenetic repression. SVA elements represent a notable exception, exhibiting both a relatively high copy number (4,972) and a moderate average correlation (0.151). Furthermore, no consistent relationship between correlation strength and K2P divergence was observed (Figure 3B), indicating that both evolutionarily young and old elements are predominantly quiescent. We also tested the observed dependencies against an average length of TE elements by families, observing no dependency between family-level averages of length and K2P divergence (Supplementary Figure 6B) and between average absolute correlation and length (Supplementary Figure 6C). As a final validation against the random background, FDR-corrected Mann-Whitney comparison between the observed and the random correlations revealed significant differences for all TE classes, the most significant ones for LTR elements (Supplementary Figure 6D), albeit the absolute magnitudes of change between the observed and the random distribution were quite low except SVA elements (0.015 average absolute Spearman correlation for DNA elements, 0.038 for LINEs, 0.034 for LTRs, 0.006 for Helitrons, 0.015 for SINEs and 0.140 for SVAs).

**Figure 3.**
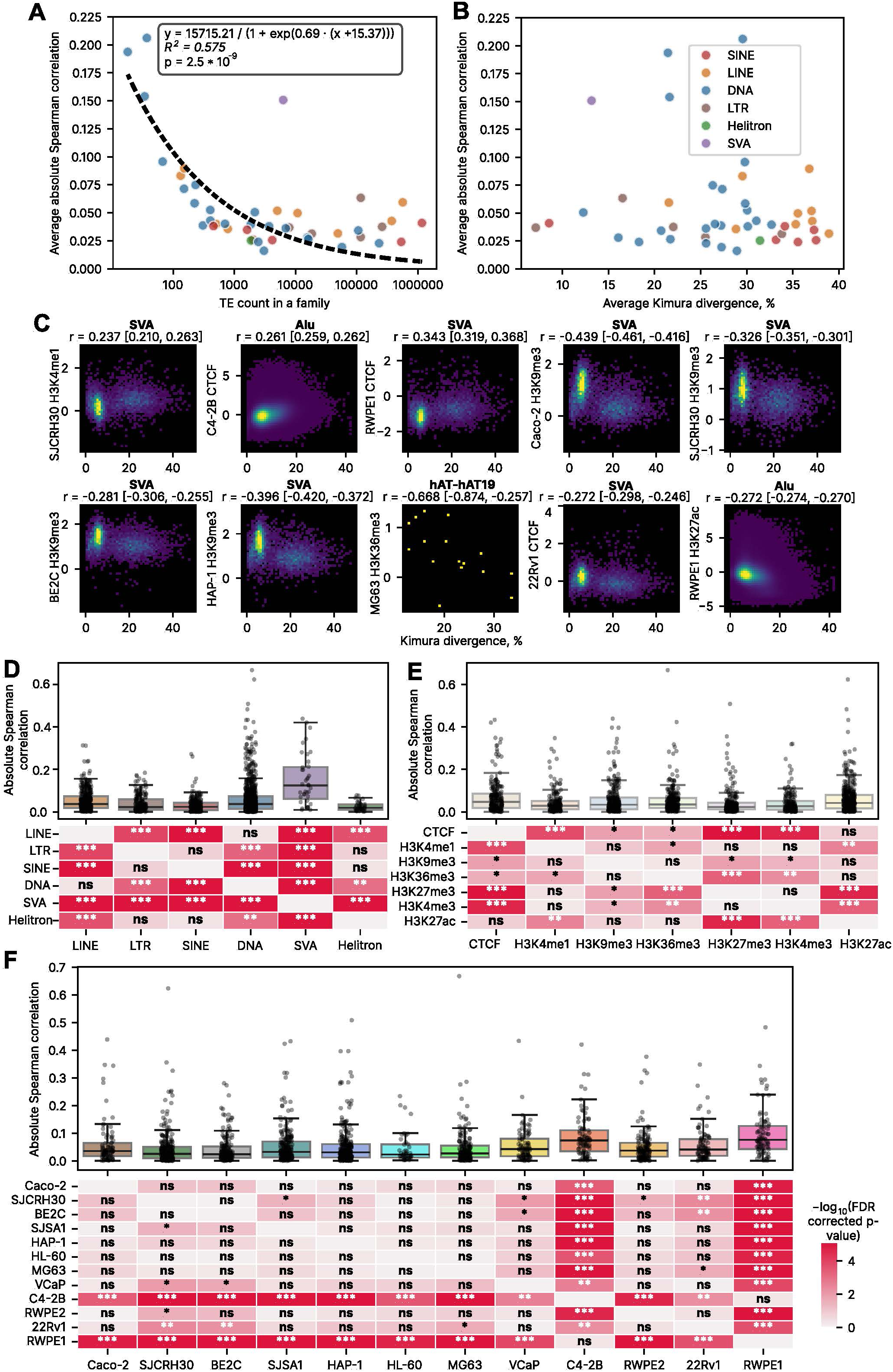
Evolutionary relationships between regulatory impact and divergence at the level of TE families. (A) Scatter plot showing the relationship between the average absolute Spearman correlation (ChIP–seq enrichment versus Kimura divergence, averaged across cell lines and epigenetic tracks) and the number of elements per TE family. The dashed curve represents the sigmoidal approximation, with the fitted equation shown in the inset. R^2^ and p values are reported for the predicted and observed data. (B) Scatter plot of the same average absolute Spearman correlation versus the average Kimura divergence of TE families. (C) Heatmaps illustrating the dependence of ChIP–seq signal on divergence for 10 TE family–cell line–epigenetic modality combinations, for which the lower bound of the 95% confidence interval exceeded 0.2 or the upper bound was below −0.2. Confidence intervals are indicated in square brackets; color intensity reflects the number of observations per bin. (D) Comparison of absolute Spearman correlations across TE classes. (E) Comparison of absolute Spearman correlations across epigenetic modalities. (F) Comparison of absolute Spearman correlations across cell lines. For all group comparisons, significance is assessed by the Mann–Whitney test and FDR-corrected: ns, p > 0.05; *, 0.01 < p < 0.05; **, 0.001 < p < 0.01; ***, 0.0001 < p < 0.001; ****, p < 0.0001.

In general, the distribution of Fisher *z*-transformed Pearson correlation coefficients was again highly non-normal (Shapiro–Wilk *p =* 6.7*10^-38^), symmetric, and strongly leptokurtic (kurtosis = 11.89, p-value = 3.6*10^-76^), reflecting heterogeneous evolutionary modes among TE families and highlighting the subset of families with active epigenetic evolution. Given the substantial number of low-copy families (e.g., hAT-Tip100?, TcMar-Pogo, hAT?, hAT-hAT19 with fewer than 100 members, and 12 additional families with 100–1,000 members), we calculated 95% confidence intervals for the Spearman correlations, requiring a lower bound of at least 0.2 to identify low but statistically significant correlations. Twelve TE family–mark combinations met these criteria, with four showing significantly positive and eight significantly negative correlations (Table 1). Among these, seven were associated with SVA elements, two with Alu elements, and one with hAT-hAT19 (18 members). Notably, H3K9me3 and CTCF exhibited four significant correlations each, indicating that heterochromatin evasion and chromatin architectural integration constitute the predominant modes of the evolutionary arms race between TEs and host defense systems. Representative significant correlations for SVA elements in C4-2B (CTCF) and RWPE1 (H3K27ac) are shown in Figures 2B, 2C, with the remaining significant associations illustrated in Figure 3C. Two of them, namely Alu elements-linked CTCF in C4-2B cell line and **H3K27ac** in **RWPE1**, were found among the significant correlations both for all Kimura divergence TE dataset and those trimmed by 20, 22.5 and 25% threshold divergence values (Supplementary Figure 6E)

**Table 1.** TE families displaying significant positive or negative correlations with ChIP–seq enrichment signals in specific cell lines and chromatin marks. Bold rows denote cases of TE family, cell line and epigenomic modality that were robust compared to Kimura divergence cut-offs of 20%, 22.5% and 25%.

| Cell line | Epigenomic modality | Family name | Spearman r | p-value | 95% CI low | 95% CI high | Adjusted p-value |
| --- | --- | --- | --- | --- | --- | --- | --- |
| SJCRH30 | H3K4me1 | SVA | 0.237 | $4.8 \times 10^{-63}$ | 0.209 | 0.263 | $5.2 \times 10^{-62}$ |
| **C4-2B** | **CTCF** | **Alu** | **0.261** | **$<10^{-300}$** | **0.259** | **0.263** | **$<10^{-300}$** |
| C4-2B | CTCF | SVA | 0.421 | $1.3 \times 10^{-208}$ | 0.396 | 0.444 | $2.8 \times 10^{-207}$ |
| RWPE1 | CTCF | SVA | 0.343 | $1.1 \times 10^{-136}$ | 0.318 | 0.369 | $1.8 \times 10^{-135}$ |
| Caco-2 | H3K9me3 | SVA | -0.439 | $2.5 \times 10^{-229}$ | -0.462 | -0.415 | $6.0 \times 10^{-228}$ |
| SJCRH30 | H3K9me3 | SVA | -0.326 | $2.5 \times 10^{-121}$ | -0.351 | -0.300 | $3.7 \times 10^{-120}$ |
| BE2C | H3K9me3 | SVA | -0.281 | $3.5 \times 10^{-89}$ | -0.307 | -0.254 | $4.6 \times 10^{-88}$ |
| HAP-1 | H3K9me3 | SVA | -0.396 | $6.5 \times 10^{-183}$ | -0.421 | -0.371 | $1.3 \times 10^{-181}$ |
| MG63 | H3K36me3 | hAT-hAT19 | -0.668 | $4.7 \times 10^{-3}$ | -0.887 | -0.203 | $1.4 \times 10^{-2}$ |
| 22Rv1 | CTCF | SVA | -0.272 | $1.9 \times 10^{-84}$ | -0.298 | -0.245 | $2.4 \times 10^{-83}$ |
| **RWPE1** | **H3K27ac** | **Alu** | **-0.272** | **$<10^{-300}$** | **-0.274** | **-0.270** | **$<10^{-300}$** |
| RWPE1 | H3K27ac | SVA | -0.282 | $6.8 \times 10^{-91}$ | -0.308 | -0.256 | $9.0 \times 10^{-90}$ |

As illustrated in Figure 3C, H3K9me3 enrichment exhibits a consistent decrease over evolutionary time across the two successive waves of SVA proliferation, with both peaks remaining above zero in four cell lines. CTCF-associated trends were concordant between the two cell lines in which statistically significant correlations were detected. Alu elements showed a gradual reduction of the active enhancer-associated mark H3K27ac and a concomitant increase in CTCF binding with evolutionary age; however, these trends were not consistently reproduced across additional cell lines. Finally, DNA TEs of the hAT-hAT19 family displayed a decrease in H3K36me3 occupancy over time, again restricted to a single cell line.

To systematically evaluate the relative contributions of cell type and epigenetic modality to the TE–host evolutionary arms race, we compared the absolute values of family-level Spearman correlations between ChIP–seq enrichment and divergence across TE classes (Figure 3D), epigenetic modalities (Figure 3E), and cell lines (Figure 3F). All TE classes exhibited statistically significant differences, with SVA elements once again demonstrating the highest overall intensity of evolutionary dynamics. In contrast, effect sizes were substantially smaller across epigenetic modalities and among cell lines. For example, the constitutive heterochromatin mark H3K9me3 was statistically indistinguishable from the enhancer-associated mark H3K27ac, as was the functionally distinct pair H3K27me3 and H3K4me3. Cell lines displayed comparable degrees of biologically meaningful grouping: the prostate cancer–derived cell lines C4-2B and RWPE2 differed significantly, whereas no significant difference was observed between the osteosarcoma-derived MG63 and the prostate cancer–derived VCaP.

An orthogonal variance-based analysis using a multigroup Levene test applied to *z*-transformed correlation coefficients revealed that variance differences were far more strongly associated with TE class (p = 1.7*10^-27^) than with cell lines (p = 8.9*10^-11^) or epigenetic modalities (p = 1.5*10^-5^). Collectively, these findings indicate that the evolutionary arms race between TEs and host genome defense systems operates largely in a non–tissue-specific manner. It should be noted, however, that germline cell lineages—where TE repression is expected to be particularly stringent—were not represented in the present analysis.

### 2.4. Dependency between epigenetic activity and evolutionary age of TEs at the level of subfamilies

To examine TE–host co-evolution at higher resolution, we calculated Spearman correlation coefficients between ChIP–seq enrichment signals and Kimura divergence scores at the level of individual TE subfamilies (n = 1,122). Similar to the family-level analysis, the average absolute correlation coefficient was strongly associated with subfamily copy number (Figure 4A; sigmoidal fit R^2^= 0.509), whereas no apparent dependence on the average subfamily divergence was observed (Figure 4B). We investigated the putative dependency between these correlations and TE length in order to control for potential mappability differences by TE length (Supplementary Figure 7A) and observed higher intensity of potential evolutionary arms race in the full length LTR subfamilies, especially of the ERV1 family (Spearman r = 0.435), and a weaker analogous dependency for L1 subfamilies (Spearman r = 0.325).

**Figure 4.**
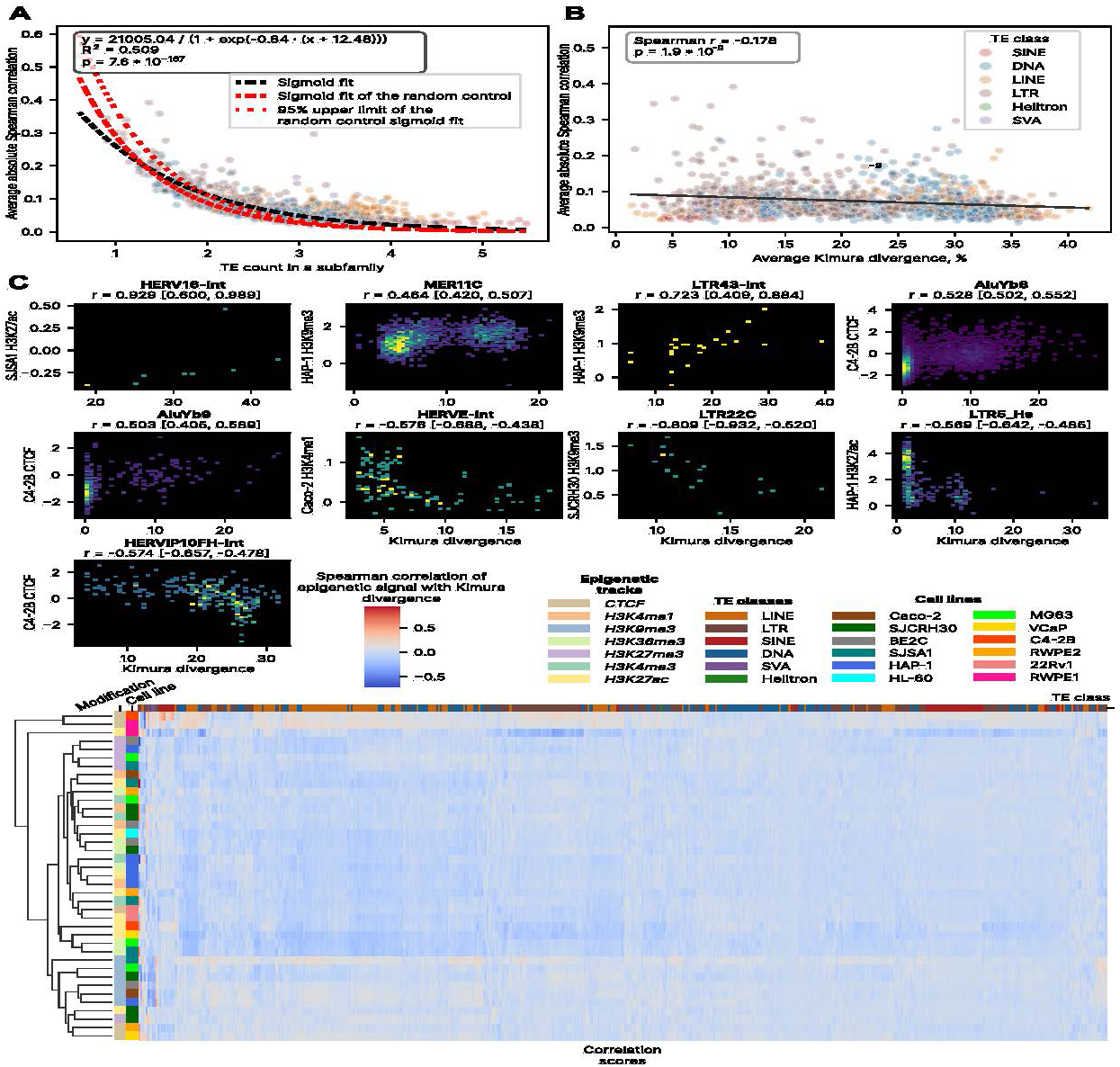
Evolutionary interplay between epigenetic activity and divergence at the level of TE subfamilies. (A) Scatter plot illustrating the relationship between the average absolute Spearman correlation (epigenetic signal versus divergence) and subfamily copy number. The black dashed curve represents the sigmoidal fit with the corresponding equation shown in the inset. R^2^ and p values are computed between observed and predicted points. The red dashed curve indicates the random-control sigmoid fit, and the red dotted line denotes the 95th percentile of random sigmoid fit residuals. (B) Scatter plot of the same average absolute Spearman correlation versus average divergence of TE subfamilies. (C) Heatmaps showing the dependence of epigenomic signal on divergence for 10 TE subfamily–cell line–epigenetic modality combinations meeting both the |r| ≥ 0.4 confidence interval criterion and the random-residual threshold. Confidence intervals are shown in brackets; color indicates bin occupancy. (D) Clustermap of correlation coefficients between epigenetic signal and divergence for 39 cell line–epigenetic modality combinations, calculated using the 500 most abundant TE subfamilies.

To control for spurious correlations arising from stochastic effects in low-copy subfamilies, we performed 100 random permutations of epigenetic signal values within each subfamily and modeled the resulting correlations as a function of subfamily size. The 95th percentile of the relative residuals between observed and sigmoid-predicted random correlations was used as a significance threshold (Figure 4A, Supplementary Figure Figure 7B), such that only correlations exceeding this boundary were considered non-random. An additional criterion of the closest to zero 95% confidence interval limit to be at least 0.4 by the absolute value allowed to retrieve 9 cases of significant moderate correlation between epigenetic signal intensity and evolutionary age (Table 2, Figure 4B). These cases likely represent ongoing evolutionary conflicts between specific TE subfamilies and host defense mechanisms.

**Table 2.**
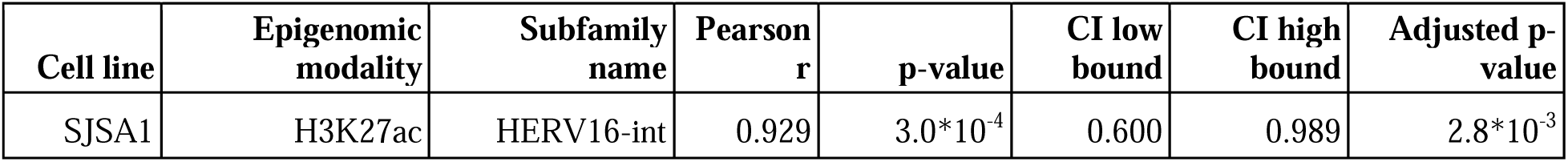
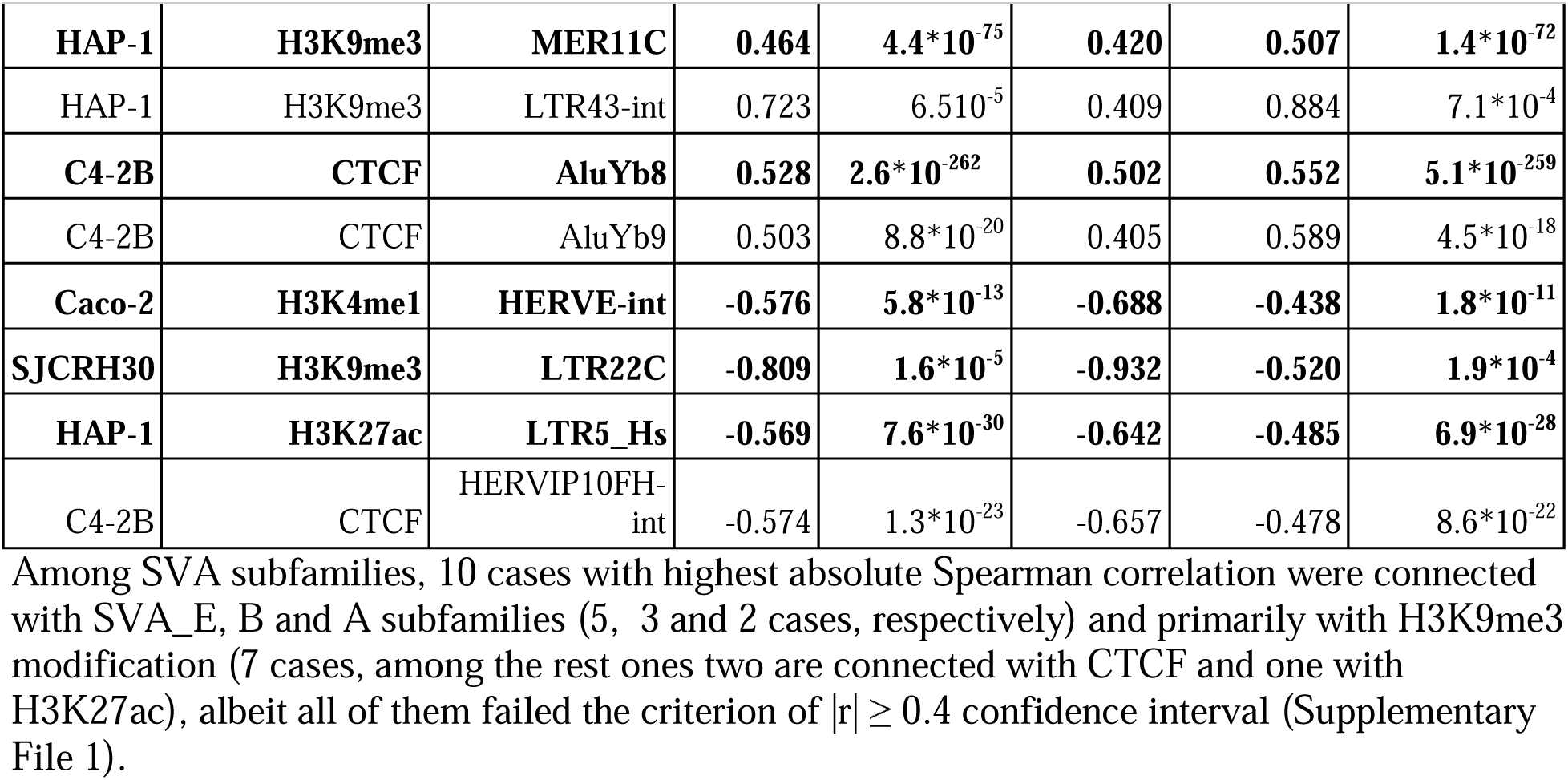
TE subfamilies, epigenetic modalities, and cell lines exhibiting significant correlations between ChIP–seq enrichment and evolutionary age. Subfamilies were additionally filtered using the 95th percentile threshold of the random correlation residuals derived from the sigmoid fit. The subfamilies marked in bold were robust in the intersectional analysis (Supplementary Figure 7C).

Significant subfamilies belonged exclusively to Alu (AluYb8, AluYb9) and LTR (HERV16-int, MER11C, LTR43-int, HERVE-int, LTR22C, LTR5_Hs, HERVIP10FH-int) elements, with no DNA transposons, no LINEs and, surprizingly, no SVA elements detected among them. At the subfamily level, these two TE classes were represented in a highly non-uniform manner (7 LTR and 2 SINE subfamilies), in contrast to the family-level analysis in which 11 out of 14 significant cases involved SVA elements. As observed at higher hierarchical levels, the dominant signatures of active evolutionary interaction were associated with chromatin architectural regulation (CTCF, 3 subfamilies) and evasion of constitutive heterochromatin (H3K9me3, 3 subfamilies). Active LTR elements were to the significant degree intact subfamilies: 4 out of 7 subfamilies, compared to 102 out of 559 total LTR subfamilies. Notably, the rest 24 AluY subfamilies showed absolute Spearman correlation at a mean level of 0.054 with standard deviation of 0.062.

Among SVA subfamilies, 10 cases with highest absolute Spearman correlation were connected with SVA_E, B and A subfamilies (5, 3 and 2 cases, respectively) and primarily with H3K9me3 modification (7 cases, among the rest ones two are connected with CTCF and one with H3K27ac), albeit all of them failed the criterion of |r| ≥ 0.4 confidence interval (Supplementary File 1).

To evaluate the robustness of the observed top TE subfamilies by evolutionary age, we also performed intersectional stability analysis of subfamilies with the closest to zero 95% confidence interval limit to be at least 0.4 by the absolute value under Kimura divergence scores clipped by 20%, 22.5% and 25% and without clipping (Supplementary Figure 7C). There were 3 consensus top subfamilies: AluYb8-linked CTCF in C4-2B, MER11C-linked H3K9me3 in HAP-1 and HERVK11D-int linked H3K36me3 in RWPE2 – the latter did not pass the random 95% percentile filtering. Also, we have found 4 consensus bottom subfamilies that showed their upper CI bound lower than 0.4: HERVE-int linked H3K4me1 in Caco-2, LTR5_Hs linked H3K2 7ac in HAP-1, LTR22C linked H3K9me3 in SJCRH30 and HERV-Fc1_LTR3 linked H3K36me3 in SJCRH30, the latter again did not pass the random filtering.

Overall, only 374 out of 43,758 tested subfamily–cell line–epigenetic modality combinations (0.85%, 1122 TE subfamilies * 39 cell lines and epigenetic modalities) exhibited moderate correlations (|r| ≥ 0.4), while 2,286 cases (5.2%) showed weak correlations within the range of 0.2 ≤ |r| < 0.4 (Figure 4D), indicating that the vast majority of extant TE subfamilies are currently evolutionarily quiescent with respect to host epigenetic regulation. Despite general weakness of the correlations, they were significantly higher than the random ones by all TE classes except Helitrons according to Mann-Whitney FDR-corrected test (Supplementary Figure 8A), albeit the absolute differences between the TE class-level medians were low: 0.022 for DNA elements, 0.05 for LINEs, 0.02 for LTRs, 0.052 for SVA and 0.019 for SINEs.

The distribution of correlation coefficients at the subfamily level was even more strongly non-normal and leptokurtic than at the family or class levels (Shapiro–Wilk p = 4.4 *10^-98^), symmetric, and extremely heavy-tailed (kurtosis = 96.8, p < 10^-200^), highlighting the highly selective nature of active TE–host evolutionary interactions. This pattern was further supported by pronounced variance differences in z-transformed correlations between TE classes (Levene test p = 4.3*10^-170^), epigenetic modalities (p = 1.6*10^-34^), and cell lines (p = 2.5*10^-125^).

Finally, clustering of epigenomic modalities based on their correlations with divergence revealed a non-random organization. Four out of five H3K27me3 cell lines (all ones except the SJCRH30 one) were co-clustered, and all cell lines harboring H3K9me3 modification were co-clustered with CTCF in RWPE2 and VcaP and with H3K27me3 and H3K27ac in SJCRH30, demonstrating large-scale grouping of heterochromatin dependencies in different TE classes and subfamilies.

The remaining epigenetic signals did not show consistent grouping, nor did cell lines, although both modalities and cell lines displayed significant non-random structure by PERMANOVA test (p < 10^-6^ for cells lines and epigenomic modalities; 10^6^ permutations). As in the case of TE families analysis, we also tested the robustness of the observed clustering patterns under Kimura divergence scores being clipped by 20%, 22.5% and 25% - and we found no significant differences of the cophenetic distance in pairwise comparisons of the clipped and the full dataset dendrogram when compared with 500 random permutations of the dendrograms, with the single exception of 22.5% dendrogram compared with the dendrogram obtained on the unclipped dataset (Supplementary Figure 8B).

Finally, the co-clustering of repressive chromatin marks was conserved when clustering was restricted to subfamilies belonging to major TE classes (LINEs, SINEs, LTRs), whereas DNA elements did not exhibit comparable stability (Supplementary Figure 9). LINE clustering revealed one basal cluster of H3K9me3 with CTCF (Supplementary Figure 9A), for SINEs the CTCF and H3K9me3 cluster was less pronounced, located within a larger group of H3K36me3 and H3K4me1 (Supplementary Figure 9B), for LTRs all cell lines with H3K9me3 formed a single group without other modalities (Supplementary Figure 9C). DNA elements revealed no co-clustering of epigenomic modalities, whereas a single cell line HAP-1 was co-clustered in a single exclusive group.

## 3. DISCUSSION

### 3.1. Kimura 2 parameters divergence from the consensus sequence as a proxy for evolutionary time

According to the neutral theory of molecular evolution (Kimura 1991), the sequence divergence of an individual TE from its subfamily consensus, being corrected for multiple substitutions and GC-content, can serve as an approximate measure of the evolutionary time elapsed since its insertion (Kawase and Ichiyanagi 2023). However, this metric becomes unreliable for highly diverged or fragmented elements. To address this limitation, a more sophisticated *defragmentation* framework was proposed, based on the principle that recent TE insertions can disrupt or fragment older ones, but not vice versa. This rescaling method has been successfully applied to over 40% of all human TEs (Giordano *et al*. 2007).

Previous studies have also employed Kimura divergence—either for entire elements or their long terminal repeats (LTRs)—as a proxy for evolutionary age. This approach has been used in the investigation of salamander genomes linking extreme genome size to reduced recombinational deletion of LTRs, leaving only the so-called “solo-LTRs” (Frahry *et al*. 2015), in studies of *Alu* elements showing loss of activity beyond 10% divergence (Bennett *et al*. 2008), as well as in genomic analyses of TEs in coelacanth (Chalopin *et al*. 2014), giraffe (Petersen *et al*. 2021) and wheat (Wicker *et al*. 2022).

In line with this extensive precedent and considering both the simplicity and availability of divergence data in the T2T genome assembly, we employed Kimura 2-parameters divergence as a practical and widely accepted proxy for TE evolutionary age in this study. Independent validation using recent evolutionary age estimates (Kosuge, Ito, and Hamada 2024) further confirmed the robustness of this approximation (Figure 1D). Finally, the ability to observe biological differences between TE classes (Supplementary Figure 1A) additionally validated K2P CpG adjusted divergence as a main metric of evolutionary age for TEs in the current study.

### 3.2. Evolutionary dependence of epigenetic signals on TE divergence

The prevailing model of TE evolutionary dynamics states that newly integrated TE loci are primarily detected by host genome defense machineries as exogenous genetic sequences, leading to their transcriptional silencing via targeted DNA methylation and mutagenic inactivation (Schlesinger *et al*. 2013, Turelli *et al*. 2014). Over time, however, as mutations accumulate and render their ORF non-functional, the host genome may co-opt the remaining TE sequences - particularly their transcription factor binding sites (TFBS) - for regulatory means (Buttler and Chuong 2022, Oomen and Torres-Padilla 2024). This functional exaptation involves the progressive selection of mutations that repurpose former viral regulatory elements into cis-regulatory modules, thereby contributing to the diversification of transcription factor binding landscapes and modulating host gene expression in an evolutionary context (Buzdin, Prassolov, and Garazha 2017). Alternatively, TEs may evade host repression and regain proliferative potential, giving rise to the classical KRAB–ZNF–mediated evolutionary arms race (Yang P, Wang, and Macfarlan 2017b).

Under such a framework, evolutionarily active TE families are expected to display systematic changes in occupancy by key epigenetic marks over evolutionary time, including histone modifications and DNA-binding proteins such as CTCF. A similar approach was recently implemented using ChIP–seq–based modeling of the KRAB–ZNF–TE arms race (Kosuge, Ito, and Hamada 2024), demonstrating that younger TE subfamilies—primarily LTRs, LINEs, and SVA elements—escape repression by pre-existing KRAB–ZNFs, followed by the emergence of new repressors through gene duplication. Interestingly, older KRAB–ZNFs were shown to repress younger TE subfamilies, whereas the reciprocal pattern was not observed, implying progressive weakening of heterochromatinization with TE age.

Consistent with this model, we observed a progressive decrease of H3K9me3 enrichment in SVA elements with evolutionary time (Figures 2A, 3C). Moreover, H3K9me3—rather than H3K27me3—robustly formed a distinct correlation cluster at the level of TE classes under Kimura divergence upper clip by 20%, 22.5% and 25% (Supplementary Figure 2), highlighting its central role in the repression of newly integrated TEs, as previously suggested (Lupo *et al*. 2013). Notably, H3K9me3 and H3K27me3 profiles formed distinct clusters at the level of TE subfamilies specifically for retrotransposons (Figure 4D, Supplementary Figure 9) but not at the family level (Supplementary Figures 3-5), demonstrating scale-specific organizational principles.

### 3.3. TE classes, families, and subfamilies exhibiting the most rapid evolutionary arms race

Among the six major TE classes analyzed (LINEs, SINEs, LTRs, DNA elements, Helitrons, and SVAs), only SVA elements displayed consistent signatures of an active evolutionary arms race (Figure 2). SINE elements had only two outlying cases of accelerated evolution (CTCF in C4-2B and H3K27ac in RWPE1), all the rest classes showed a relatively quiscent dynamics. This could be in part explaned by the bimodal distribution of evolutionary age in SVA and SINE elements, albeit more mechanistic explanations are possible. A key example of the SVA-related conflict is the stepwise evolution of the host gene ZNF91, which adapted specifically to repress successively younger SVA subfamilies in primates (Jacobs *et al*. 2014). Strong evidence also exists for a parallel arms race involving LINE-1 elements, as ZNF93 evolved specifically to repress L1PA subfamilies that had mutated to evade ancestral repressors (Castro-Diaz *et al*. 2014, Jacobs *et al*. 2014). At a broader scale, vertebrate-wide analyses have shown massive expansions of the KRAB–ZNF family driven by the necessity to repress successive waves of LTR retrotransposons (Thomas and Schneider 2011).

Nevertheless, a recent ChIP–seq–based analysis (Kosuge, Ito, and Hamada 2024) identified 24 KRAB–ZNF binding sites in DNA elements, 87 in SINEs, 101 in SVAs, 313 in LINEs, and 979 in LTRs. When normalized by TE abundance according to the same research, these correspond to 6.2*10^-5^, 4.3*10^-5^, 7.8*10^-4^, 2.7*10^-4^ and 1.4*10^-3^, respectively, indicating the highest relative conflict intensity in LTRs, followed by SVAs and LINEs. Methodological differences between these studies and the present work likely explain the observed discrepancies.

Our results indicate that SVA elements progressively accumulate CTCF binding, evade H3K9me3-mediated repression, and acquire enhancer-associated H3K4me1 in a subset of cell lines (Figure 2A). At the family level, both SVA and Alu elements showed strong signatures of an evolutionary arms race, with SVAs deviating by ∼0.13 in average correlation above the sigmoid background (Figure 3A). Notably, the two-peak evolutionary pattern observed for SVAs (Figure 3B) was not attributable to distinct subfamilies: all three SVA subfamilies found in the top 10 SVA cases by highest absolute Spearman correlation (SVA_E, SVA_B, SVA_A)—including both retrotransposition-competent (SVA_E) and currently immobile lineages (SVA_A, SVA_B) (Burns 2019)—displayed this pattern, suggesting two putative waves of proliferation within an ongoing arms race, the pattern that was not shown previously.

The Alu subfamilies with the most prominent arms race (AluYb8, AluYb9) belong to the youngest and active AluY family (Burns 2019). Their dynamics were manifested as progressive accumulation of CTCF binding over time, consistent with the established role of Alu elements in chromatin boundary formation through CTCF recruitment (Gu *et al*. 2016). Younger copies of these subfamilies may remain transiently heterochromatinized, impairing CTCF binding (Wen *et al*. 2012). Importantly, none of the remaining 22 AluY subfamilies or other SINEs present in the T2T-lifted RepeatMasker track displayed significant correlations according to the stringent statistical thresholds for evolutionary conflict.

Finally, seven LTR subfamilies (HERV16-int, MER11C, LTR43-int, HERVE-int, LTR22C, LTR5_Hs, HERVIP10FH-int), having 1446, 1431, 27, 142, 25, 345, 952 members, respectively, also exhibited strong evolutionary dynamics involving H3K4me1, H3K27ac, H3K27ac, H3K9me3 and CTCF profiles. HERV16, an evolutionary old endogenous retrovirus, was previously reported as co-evolving with HLA class I duplications (Kulski *et al*. 1999). MER11C, an evolutionary young human LTR, is silenced by KRAB-ZNF induced DNA methylation and acts as long non-coding RNA, whose abnormal expression is linked to neurological disorders and cancer (Dini, Taheri, and Shirvani-Farsani 2024, Derakhshan *et al*. 2025) – and we showed that it has H3K9me3 enrichment decreasing with evolutionary age. LTR43 was found among HLA class I cluster retroelements whose expression is regulated by SVA elements with potential implication in Parkinson disease (Kulski, Pfaff, and Koks 2025), and its H3K9me3 heterochromatinization again weakens in younger insertions according to our current results.

HERV-E is a full length provirus whose overexpression and autoantibodies against its proteins were associated with autoimmune diseases (Mustelin and Ukadike 2020), whereas its TFBS regulate expression of human saliva amylase and apolipoprotein-C1 genes (Bannert and Kurth 2004). It is of particular interest that its H3K4me1 enhancer histone mark enrichment levels are increased in Caco-2 colorectal cell line precisely in the young copies, indicating a possible evolutionary arms-race scenario.

LTR22C was previously found downregulated in small cell lung cancer and their expression was associated with innate immune response (Russo, Morelli, and Capranico 2023). LTR5_Hs, an evolutionarily young HERV-K (HML-2) LTR (Padmanabhan Nair *et al*. 2021), was earlier reported to cause epigenomic differences in H3K4me3 of human and chimpanzee iPSC (Hirata *et al*. 2022) and to activate KRAB–ZNF ZNF75D in lung adenocarcinoma cells (Ito *et al*. 2020). We now show that its younger copies of less than 5% Kimura divergence harbor elevated H3K27ac enrichment signal in HAP-1, which could implicate cell type specific manifestation of evolutionary arms race in bone marrow. HERVIP10FH-int, which is an intact but nonautonomous insertion of ERV-1 (Shah *et al*. 2022), has not previously been discussed in the context of TE–host evolutionary conflict. Given the fact that LTR elements in general are evolving to confront the defence systems pressure (Thomas and Schneider 2011), our results represent the novel evidence implicating these specific subfamilies in an active evolutionary arms race.

### 3.4. Relative impact of distinct epigenomic modalities in the TE–host evolutionary arms race

By jointly analyzing seven epigenomic modalities across TE classes, families, and subfamilies, we demonstrate that repressive chromatin states dominate the evolutionary arms race. The near-perfect co-clustering of H3K27me3 and H3K9me3 profiles associated of CTCF profiles across cell lines, as well as the prevalence of H3K9me3 among significant cases of evolutionary arms-race, indicates that TEs primarily evolve either by evading heterochromatinization or by participating in chromatin architecture. In contrast, enhancer-associated (H3K27ac, H3K4me1), promoter-associated (H3K4me3), and gene body associated (H3K36me3) dimensions appear weaker or highly cell-type specific. The lack of clustering by modalities in DNA elements could be explained by their high evolutionary age and loss of transpositional competency due to serial point mutations (Matsushima, Planet, and Trono 2024), further exemplifying the fact that retroelements could have active epigenetic avoidance strategies.

Although the KRAB–ZNF pathway imposing H3K9me3 is the most extensively studied TE defense system, multiple additional host pathways operate against TEs, including deamination, where there is a documented arms race between APOBEC3 and L1 and LTR elements (Modenini, Abondio, and Boattini 2022b), RNA interference (Cornec and Poirier 2023), the HUSH (Human silencing hub) complex which imposes H3K9me3 (Seczynska and Lehner 2023), DNA methylation, PRC2-mediated H3K27 trimethylation (Déléris, Berger, and Duharcourt 2021, Hisanaga *et al*. 2023), MORC proteins (Yang F and Wang 2016) etc. Each of these mechanisms can potentially be evaded by TE counter-adaptation, producing intragenomic selection detectable as divergence-dependent epigenetic remodeling.

For example, Polycomb repressive complex 2 (PRC2) was shown to repress TEs in a diverse range of eukaryotes depositing H3K27me3 (Hisanaga *et al*. 2023). Although it is hypothesised to have an ancestral eukaryotic transposon-repressing role (Déléris, Berger, and Duharcourt 2021), there is no published evidence of active ongoing evolutionary arms race of PRC2 and TEs, where certain subfamilies are initially repressed and then evade the repression acquiring new mutations. Our results indicate that human TEs can selectively evade PRC2-mediated H3K27 trimethylation, suggesting the existence of an unrecognized evolutionary conflict. These bioinformatic findings may guide future mechanistic studies aimed at resolving PRC2–TE co-evolution.

H3K36me3, transcriptionally active gene body mark, was represented in a small subset of top families and subfamilies in the current study: HERVK11D-int, HERV-Fc1_LTR3 and hAT-hAT19. Integration and co-opting within H3K36me3-marked regions could represent an adaptive TE strategy to evade the host repression rather than a passive insertional bias. Since H3K36me3-enriched regions undergo enhanced DNA damage repair (Li, Ahn, and Wang 2019), they may provide selective protection to resident TEs. This hypothesis could be tested by precise reconstruction of phylogenetic trees of those TE subfamilies which show high correlation of H3K36me3, other epigenetic marks of transcription and CAGE (Cap Analysis of Gene Expression) peaks, with divergence. Detailed studies of KRAB-ZNF and other repressive factors binding in these TEs in connection with active transcription tags could shed light on the previously unexplored ways of TEs/host genome evolutionary arms race. Recently, a preprint epigenomic study identified enrichment of H3K36me3 in Alu elements (Hyacinthe and Bourque 2024), and this epigenomic mark was enriched on duplicated genes with rich TE neighbourhood (Lannes, Rizzon, and Lerat 2019), but these observations alone cannot resolve whether this reflects adaptive co-evolution.

### 3.5. Relevance of the current findings in the context of cancer and therapeutic resistance

Cancer is fundamentally an evolutionary disease arising from conflict between cellular-level selection and organismal fitness (Casás-Selves and Degregori 2011). The TE vs host genome arms race is fundamentally similar to cancer in this manner, since both TEs and cancer cells are prompting the host to evolve defense mechanisms in an ongoing cycle of counter-adaptation (Casás-Selves and Degregori 2011). Consistent with this parallel, TEs are frequently reactivated in dysregulated cancer transcriptomes (Solovyeva *et al*. 2024). Epigenetic dysregulation combined with global DNA hypomethylation (Ehrlich 2009) leads to TE reactivation, which in turn destabilizes the genome through insertional mutagenesis, aberrant recombination, and oncogene activation, thereby accelerating malignant progression(Anwar, Wulaningsih, and Lehmann 2017, Jovčevska *et al*. 2019, Shtam *et al*. 2019).

Current treatment strategies rely increasingly on complex molecular biomarkers (Sorokin Maksim *et al*. 2021, Gudkov *et al*. 2022, Sorokin Maxim *et al*. 2022, Fiore *et al*. 2025), yet resistance still emerges through tumor heterogeneity, co-mutations, and acquired genetic alterations (Sorokin Maxim *et al*. 2020, Vladimirova *et al*. 2021, Tufail *et al*. 2024). Emerging therapeutic strategies may increasingly exploit endogenous TE defense systems which are in an endless arms race with genomic parasites: epigenetic silencing (Song *et al*. 2025), deamination (Frances and Cordelier 2020) and RNA interference (Singh *et al*. 2025).

## 4. CONCLUSIONS

Using T2T-lifted epigenomic profiles and high-resolution repeat annotations, we quantified the evolutionary arms race between transposable elements and human genome defense systems across diverse chromatin states and cellular contexts. By applying Kimure 2 parameters divergence from a derived consensus sequence as a proxy for evolutionary age, we demonstrate that correlations between epigenetic activity and TE divergence are generally weak but highly structured.

We systematically evaluated the relative contributions of seven epigenomic modalities to the TE–host evolutionary conflict and demonstrate that TEs primarily evade H3K9me3- and secondarily H3K27me3-mediated repression, while integrating into chromatin structural CTCF-marked regions. In contrast, enhancer, promoter, and actively transcribing gene body dimensions of TE evolution appear comparatively weaker. These findings provide strong new evidence into the understanding of epigenetic mechanisms of TE and host genome coevolution.

Collectively, our results identify SVA elements as the most dynamic TE class in recent human evolution, both in proliferation history and regulatory potential, while AluYb8, AluYb9 and seven LTR groups (HERV16-int, MER11C, LTR43-int, HERVE-int, LTR22C, LTR5_Hs, HERVIP10FH-int). are the most dynamic TE subfamilies. By linking TE evolutionary age with epigenetic activity at class, family, and subfamily scales using T2T-resolved repeat annotations, this study provides a multi-scale, high-resolution view of TE–host coevolution and underscores the central role of young retroelements in shaping ongoing regulatory innovation in the human genome. The current results establish a framework for future investigations into genome defense, regulatory innovation, and disease evolution.

## 5. MATERIALS AND METHODS

### 5.1. Genome reference and repeats coordinates

We performed co-mapping analysis using genomic annotations and coordinates derived from the T2T complete human genome assembly (CHM13), which enabled the inclusion of the previously unresolved ∼8% of the human genome and approximately 1% additional TE content (Nurk *et al*. 2022).

Coordinates and class/family annotations for human TE were obtained from the RepeatMasker track (Tarailo-Graovac and Chen 2009) of the same assembly and via the same *Table Browser* tool (Group *Variation and Repeats*, Track *RepeatMasker*, Table *T2T RepeatMasker*). The T2T RepeatMasker track was derived from (Hoyt *et al*. 2022).

Each TE was classified by RepeatMasker into class, family and subfamily (Tarailo-Graovac and Chen 2009):

- Class: the highest classification level based on mechanism of movement.
- Family: the medium level classification based on shared, similar ancestry, often sharing similar structures and transposition mechanisms (such as Alu and L1 elements).
- Subfamily: the lowest classification level, specific, often recently evolved, distinct lineage within a family (such AluY, AluS, AluJ).

We used the T2T genome RepeatMasker classification without any amendments to comply with RepeatMasker widely accepted classification.

The dataset included elements classified as LINE, SINE, LTR, SVA (listed as *Retroposon* in the table), Helitron (noted as *RC, Rolling Circle*), and DNA transposons, comprising a total of 3,709,429 entries: 1,706,485 SINEs, 1,005,214 LINEs, 531,410 LTRs, 458,177 DNA elements, 6,274 SVAs, and 1,869 Helitrons.

Despite the functional and evolutionary connections between SVA and Alu elements (Jacobs *et al*. 2014), SINEs and SVAs were treated as different classes in the current research according to the RepeatMasker nomenclature. In the same way, Helitrons were considered as a separate TE class albeit mechanistically they can be considered as a subgroup of DNA transposons due to their DNA-based rolling circle replication (Barro-Trastoy and Köhler 2024).

### 5.2. Evolutionary age assessment of repeats

Kimura divergence scores were obtained through the Kimura 2-parameters (K2P) value, which is available in the .align outputs of the Repeat Masker standard runs via the link (https://www.repeatmasker.org/genomes/hs1/rmsk4.2.2_dfam3.9_rb20181026_rmb/hs1.fa.align.gz). For each hit, the “align” file showed a line like the standard UCSC Repeat Masker track, followed by the alignment followed by statistics including a line “Kimura (with divCpGMod) = XXX”. In case multiple alignments segments of a given TE insertion exist in the “align” file intersecting with its coordinates by the UCSC Repeat Masker table, the divergence K2P value was calculated by computing the weighted average, using the length of each sub hit (each segment) as a weight. Since LINE elements in the human lineages have been split in two to three segments (5’ end, 3’ end and orf2) in order to improve the precision of Repeat Masker (RepeatMasker Frequently Asked Questions, n.d.), the individual LINE element hits matching the 3’ end and orf2 (perhaps 5’ end if full-length hit) in the .align file were again averaged using their lengths as weight. Generally, 157,858 TEs out of 3,709,429 (4.26%) lacked K2P divergence assessment and were not investigated in the further analysis.

### 5.3. Epigenome data

We performed a tiered ChIP-seq–based analysis using datasets generated by the ENCODE consortium under the T2T ENCODE initiative (Gershman *et al*. 2022). Epigenomic enrichment profiles were obtained from the UCSC Genome Browser download repository at (GBDB T2T ENCODE, n.d.). We used enrichment tracks computed as log ratios of experimental versus control coverage following the standard T2T ENCODE ChIP-seq pipeline (Schema for T2T Encode - T2T Encode Reanalysis, n.d.).

In order to get the final enrichment tracks, sequencing reads were aligned to the T2T-CHM13v2.0 human reference genome using Bowtie2 (v2.4.1) in paired-end mode with the following parameters:

--no-discordant --no-mixed --very-sensitive --no-unal --omit-sec-seq --xeq --reorder.

Alignments were filtered with SAMtools (v1.10) using the flags -F 1804 -f 2 -q 2 to remove unmapped, improperly paired, or low-quality (MAPQ < 2) reads. PCR duplicates were identified and removed using Picard Tools (“MarkDuplicates”, v2.22.1) with the parameters VALIDATION_STRINGENCY=LENIENT ASSUME_SORT_ORDER=queryname REMOVE_DUPLICATES=true.

Filtered alignments were subsequently screened for unique k-mers using the *minUniqueKmer* utility (Msauria/MinUniqueKmer) and merged across biological replicates. BigWig coverage tracks were generated with deepTools bamCoverage (v3.4.3) using a bin size of 1 bp and default settings for all other parameters. Enrichment tracks were computed using deepTools bamCompare with a bin size of 50 bp, a pseudo-count of 1, and excluding bins with zero coverage in both target and control datasets.

For downstream analyses, we utilized continuous enrichment signal tracks rather than discrete peak calls, as this approach provided more comprehensive coverage of RE-associated loci. Peak-calling software such as MACS typically identifies fewer than 120,000 peaks (the maximum number within the T2T ENCODE cohort was 118,387 as of October 21, 2025), which would have limited the scope of RE-mapping resolution.

The epigenomic enrichment files were downloaded for the following 12 cell lines from 6 organs/tissues (Figure 1A):

- 22Rv1, prostate carcinoma
- BE2C, neuroblastoma
- C4-2B, epithelial-like morphology, isolated from prostate
- Caco-2, colorectal adenocarcinoma
- HAP-1, chronic myelogenous leukemia
- HL-60, peripheral blood, acute promyelocytic leukemia
- MG63, bone, osteosarcoma
- RWPE1 - epithelial cell from prostate
- RWPE2 - epithelial cell from prostate
- SJCRH30 - fibroblast-like cell line from muscle
- SJSA1 - primitive multipotential sarcoma of the femur
- VcaP - epithelial cell from prostate

Epigenomic datasets were obtained for CTCF and six histone modifications (H3K4me1, H3K9me3, H3K36me3, H3K27me3, H3K4me3, and H3K27ac). In total, 39 chromatin profiles were analyzed, reflecting the fact that not every histone mark was available for each cell line within the T2T ENCODE resource. All epigenomic files used in this study, along with their corresponding download links, are listed in Supplementary File 2.

### 5.4. Mapping of epigenetic signal on TEs

ChIP–seq enrichment bedGraph files were intersected with TE genomic coordinates using bedtools (Quinlan and Hall 2010). For each TE, enrichment values overlapping its genomic interval were extracted and averaged.

### 5.5. Drawing pictures

Iconographic representations of organs and tissues were made using BioRender (Scientific Image and Illustration Software | BioRender, n.d.). All diagrams were generated using the Python libraries matplotlib, matplotlib-venn, and seaborn (Hunter 2007, Waskom 2021). Supervenn plots were drawn using the Python library supervenn (GitHub - gecko984/supervenn: supervenn: precise and easy-to-read multiple sets visualization in Python · GitHub, n.d.). Intermediate SVG files were rendered and refined in Figma (Figma Downloads | Web Design App for Desktops & Mobile, n.d.). Annotation of boxplots with Mann-Whitney p-values was done using statannotations python module (GitHub - trevismd/statannotations, n.d.).

### 5.6. Statistical tests, correlations and data analysis

All computational analyses were conducted in Python using the pandas and numpy libraries for data processing and numerical calculations (Harris et al., 2020; Pandas development team, n.d.). Correlations between TE divergence and epigenetic enrichment scores were computed using Spearman’s correlation coefficient (Rodgers and Nicewander 1988), selected for its property of weighting covariance by the ranks of observations, which is appropriate for detecting non-linear potential dependencies between evolutionary age and epigenetic activity.

Associated *p*-values for Spearman correlations were obtained using the scipy.stats module (Virtanen *et al*. 2020). To evaluate the statistical properties of the correlation coefficients, we applied Fisher’s z-transformation (Fisher 1915), implemented in numpy. Ninety-five percent confidence intervals (CIs) were derived from the Fisher-transformed values using the standard error of the transformed correlations and the 95% quantiles of the corresponding normal approximation (Tian and Wilding 2008), as provided by scipy.stats.

Distributional normality was tested with the Shapiro–Wilk test (Shapiro and Wilk 1965) (scipy.stats). Kurtosis values and their significance (Mardia 1970) were likewise computed using scipy.stats. Group comparisons were performed using the two-sided Mann–Whitney–Wilcoxon test (Wilcoxon 1945, Mann and Whitney 1947) (for median differences) and the Levene test (for variance differences) (Gastwirth, Gel, and Miao 2009). Adjustment for multiple hypothesis testing was applied using the Benjamini–Hochberg false discovery rate (Benjaminit and Hochberg 1995) procedure as implemented in the statsmodels Python package (Seabold and Perktold 2010).

Clustering of correlation coefficients was done using python seaborn library (Waskom 2021) in Euclidean metrics space with average linkage method. Permutation test of clustering trees was done using permutational ANOVA test (Permanova) with 1 mln iterations implemented in python skbio library (GitHub - scikit-bio, n.d.). Similarity between dendrograms was assessed using Spearman cophenetic correlation (Shamsuri and Sharif 2024).

### 5.7. Sigmoid approximations and their significance

Sigmoid models describing the relationship between correlation strength and the number of elements within a TE family or subfamily were fitted using the standard logistic function (Sun 2025):

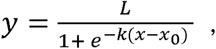

where *x* denotes the family or subfamily size, *y* represents the mean absolute Pearson correlation between divergence and epigenetic enrichment, and *L*, *k*, and *x₀* are parameters to be estimated. Initial parameter guesses were set to the maximum observed *y* value (for *L*), 1 (for *k*), and the median of *x* (for *x₀*). Parameter optimization was carried out using non-linear least squares as implemented in scipy.optimize (Virtanen *et al*. 2020), with a maximum of 5000 iterations. Coefficient of determination (*R²*) and associated *p*-values for the fitted curve were computed using Pearson correlation, following the scipy.stats implementation.

To assess whether the empirical sigmoid trend deviated significantly from random expectations, we performed a permutation-based background analysis (John, Korte, and Grimm 2024). For each TE family or subfamily and each cell line/epigenetic modality pair, values of ChIP-seq enrichment and divergence were independently shuffled in 100 iterations, followed by recalculation of the Pearson correlation coefficient. The resulting random and observed relationships between TE count and mean absolute correlation were compared using the nested-model F-test framework (El-Horbaty 2018).

For TE subfamilies, we additionally constructed a one-sided 95% prediction interval for the random sigmoid fit. To account for heteroscedasticity (You *et al*. 2023), this interval was computed using relative errors: residuals were expressed as (observed – predicted) / predicted, and the 95th percentile of these relative residuals was used as a multiplicative factor to scale the random-fit curve. The scaled curve served as the upper prediction boundary (Hazra 2017); subfamilies with observed values exceeding this threshold were considered to show statistically significant deviation from the random expectation.

### 5.8. AI usage

Gemini 2.5 Pro (Comanici *et al*. 2025) was employed for code refinement and the implementation of statistical methods within the Jupyter notebook environment. ChatGPT-5 (Introducing GPT-5 | OpenAI, n.d.) was used exclusively for orthographic, grammatical, and punctuation review.

### 5.9. Code availability

Jupyter notebooks used for all data analyses, Python and Bash wrapper scripts employed for epigenomic data acquisition, format conversion, bedtools-based genomic mapping, and correlation analyses have been uploaded to a public GitHub repository and are accessible via the link https://github.com/Nikit357/T2T_transposons_evolution/.

## 6. DATA AVAILABILITY

The data underlying this article are available in the article and in its online supplementary material. Intermediate analysis files, large Jupyter notebooks containing figure outputs, and additional plots generated during exploratory analyses but not included in the article or supplementary files have been deposited in a public GitHub repository and are accessible via the link https://github.com/Nikit357/T2T_transposons_evolution/.

## 7. ETHICAL STATEMENT

This study constitutes purely computational research utilizing publicly available epigenomic and genomic datasets. All analyzed data were derived from established human cell lines (22Rv1, BE2C, C4-2B, Caco-2, HAP-1, HL-60, MG63, RWPE1, RWPE2, SJCRH30, SJSA1, VcaP) originally generated and characterized by the ENCODE Consortium (GBDB T2T ENCODE, n.d.) and available via public repositories. As the research did not involve any new primary collection of human or animal tissue, or any direct involvement of human participants or live animals, formal ethical approval and informed consent were not required. All data acquisition and processing adhered to the standard ethical guidelines for the use of secondary biological data and the T2T ENCODE initiative (Gershman *et al*. 2022).

## 9. AUTHORS CONTRIBUTION

**Daniil Nikitin**: Conceptualization, data curation, formal Analysis, funding acquisition, investigation, methodology, project administration, resources, software, supervision, validation, visualization, writing – original draft, writing – review & editing.

As the sole author, I meet all the required criteria for authorship: I made substantial contributions to the conception, design, acquisition, analysis, and interpretation of the data; I drafted and critically revised the manuscript for important intellectual content; I have given final approval of the version to be published; and I agree to be accountable for all aspects of the work.

## 10. ACKNOWLEDGEMENTS

My profound and heartfelt thanks go to my wife, Irina Nikitina. It was her gentle encouragement and unwavering belief in me that transformed an evolutionary biology hobby into the series of research articles, including the current one. Her love and support were my anchor, preserving my passion and courage through the dark times of injuries, illnesses, and the many challenges of our relocation to Armenia with young sons. I also thank the anonymous reviewers of Molecular Biology and Evolution journal who helped me to obtain reliable approximations of TEs evolutionary age and to improve the statistical robustness of the analysis.

This research received no specific grant from any funding agency in the public, commercial, or not-for-profit sectors.

## 11. SUPPLEMENTARY MATERIAL

**Supplementary Figure 1**. (A) Joint scatter plots showing correlation of Kimura CpG adjusted divergence (the main evolutionary age metric in the current study) with other evolutionary metrics: divergence by UCSC Genome Browser tables, percent divergence by T2T RepeatMasker, transitions to transversions ratio by T2T RepeatMasker, Kimura CpG unadjusted divergence by RepeatMasker. The main diagonal shows histograms of the metrics, the lower part visualizes scatter plots colored by TE class, the upper part shows joint density plots by all classes. (B) Scatter plots of TE length by Kimura divergence visualized separately for individual TE classes, with Spearman correlation coefficients and p-values.

**Supplementary Figure 2**. Clustermaps of Spearman correlation coefficients between ChIP-seq enrichment signals and Kimura divergence scores across TE classes, cell lines, and epigenomic modalities, stratified by divergence clipping. Prior to correlation calculation, all TEs with divergence above the clipping threshold were removed. (A) 20%, (B) 22.5%, (C) 25%.

**Supplementary Figure 3**. Clustermap of Spearman correlation coefficients between ChIP-seq enrichment signals and divergence scores across TE families, cell lines, and epigenomic modalities. Sidebar bar plots indicate the number of copies and the mean divergence score for each TE family.

**Supplementary Figure 4**. Clustermaps of Spearman correlation coefficients between ChIP-seq enrichment signals and Kimura divergence scores across TE families, cell lines, and epigenomic modalities, stratified by divergence clipping. Prior to correlation calculation, all TEs with divergence above the clipping threshold were removed. (A) 20%, (B) 22.5%.

**Supplementary Figure 5**. (A) Clustermap of Spearman correlation coefficients between ChIP-seq enrichment signals and Kimura divergence scores across TE families, cell lines, and epigenomic modalities. Prior to correlation calculation, all TEs with divergence above the clipping threshold of 25% were removed. (B) Comparison of Spearman cophenetic correlation of the two dendrograms against the random background distances obtained from 500 random permutations of these two dendrograms. The first one was derived from Spearman correlation coefficients between ChIP-seq enrichment signals and Kimura divergence scores by TE families, cell lines, and epigenomic modalities, whereas the second one was derived from clustermap of the same correlations calculated under the 20% divergence clipping. Empirical p-value was calculated as fraction of random distances greater that the relevant one. (C) Comparison of the same first dendrogram without divergence clipping against the 22.5% divergence clipped dendrogram. (D) Comparison of the same first dendrogram without divergence clipping against the 25% divergence clipped dendrogram.

**Supplementary Figure 6**. (A) Scatter plot showing average absolute Spearman correlation of ChIP-Seq enrichment signal and Kimura divergence against TE count in a family for a dataset of 100 random permutations of epigenetic signal by TEs within each family. Sigmoid equation is added, as well as R^2^ of the sigmoid fit. (B) Scatter plot of average length in a TE family versus average Kimura divergence. (C) Scatter plot visualizing average absolute Spearman correlation and average length in a TE family. (D) Box plots comparing observed and random (by 100 permutations) average absolute Spearman correlation in TE families by TE class. (E) Supervenn plot of TE families, epigenomic modalities and cell lines that were selected as top (lower CI bound above 0.2) or bottom (higher CI bound below -0.2) significantly correlating in the dataset of all divergence values and in those clipped by 20%, 22.5% and 25% divergence.

**Supplementary Figure 7**. (A) Scatter plots showing average absolute Spearman correlation of ChIP-Seq enrichment signal and Kimura divergence against average length in a subfamily by TE classes. Colors denote subfamilies of the same family on each plot. (B) Density distribution showing average absolute Spearman correlation against TE count in a subfamily for a dataset of 100 random permutations of epigenetic signal by TEs within each subfamily. Sigmoid equation is added, as well as R^2^ of the sigmoid fit. (C) Supervenn plot of TE subfamilies, epigenomic modalities and cell lines that were selected as top (lower CI bound above 0.4) or bottom (higher CI bound below -0.4) significantly correlating in the dataset of all divergence values and in those clipped by 20%, 22.5% and 25% divergence.

**Supplementary Figure 8**. (A) Box plots comparing observed and random (by 100 permutations) average absolute Spearman correlation in TE subfamilies by TE class. (B) Pairwise comparison of Spearman cophenetic correlation of the 4 dendrograms against the random background distances obtained from 500 random permutations of these two dendrograms. The “All” one was derived from Spearman correlation coefficients between ChIP-seq enrichment signals and Kimura divergence scores by TE subfamilies, cell lines, and epigenomic modalities, whereas the “cut-off” ones were derived from clustermap of the same correlations calculated under the 20%, 22.5% and 25% divergence clipping. Empirical p-value was calculated as fraction of random distances greater that the relevant one.

**Supplementary Figure 9**. Clustermaps of Spearman correlation coefficients between ChIP-seq enrichment signals and divergence scores across TE subfamilies, cell lines, and epigenomic modalities, stratified by TE class: (A) LINEs, (B) SINEs, (C) LTRs and (D) DNA transposons.

**Supplementary File 1**. Tables containing Spearman correlation coefficients between ChIP-seq enrichment signals and divergence scores at the level of individual cell lines and epigenomic modalities. Separate sheets report correlations for TE classes and families, as well as results from divergence clipping by 20%, 22.5% and 25% thresholds. Spearman correlations for subfamilies and the randomly shuffled background control correlations can be found in the GitHub repository via the link https://github.com/Nikit357/T2T_transposons_evolution/.

**Supplementary File 2**. GBDB download links for files containing ChIP-seq enrichment signals for each epigenomic modality (CTCF and histone modifications) and cell line used in the present study.

**Supplementary File 3**. Jupyter notebook containing all data processing, statistical analyses, and figure generation performed in this study.

