## Supplementary figures and images for "Transposable element–host genome evolutionary arms race revealed by multi-modal epigenomic profiling in a telomere-to-telomere human genome reference"

### Supplementary Figure 1

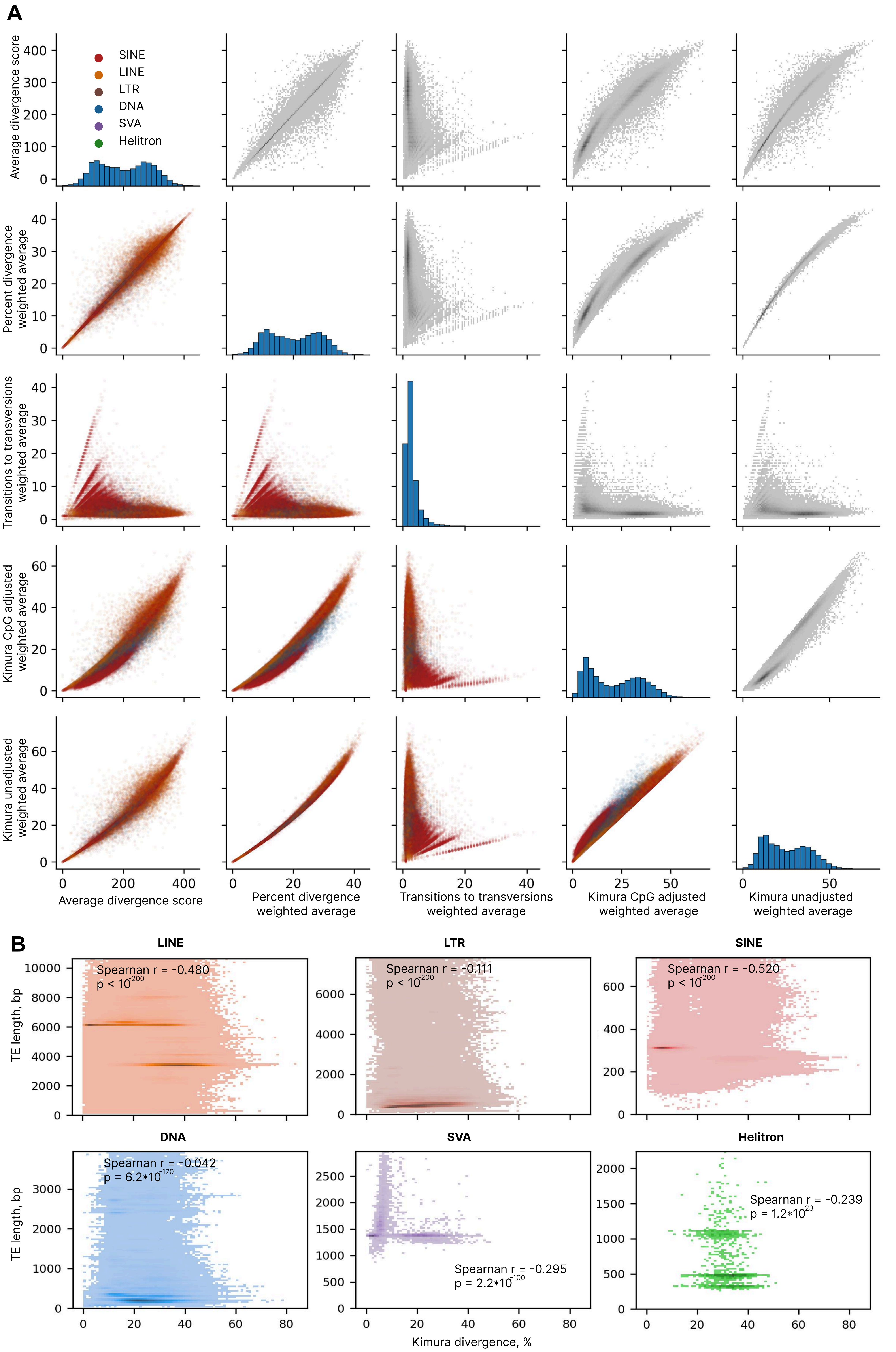

### Supplementary Figure 2

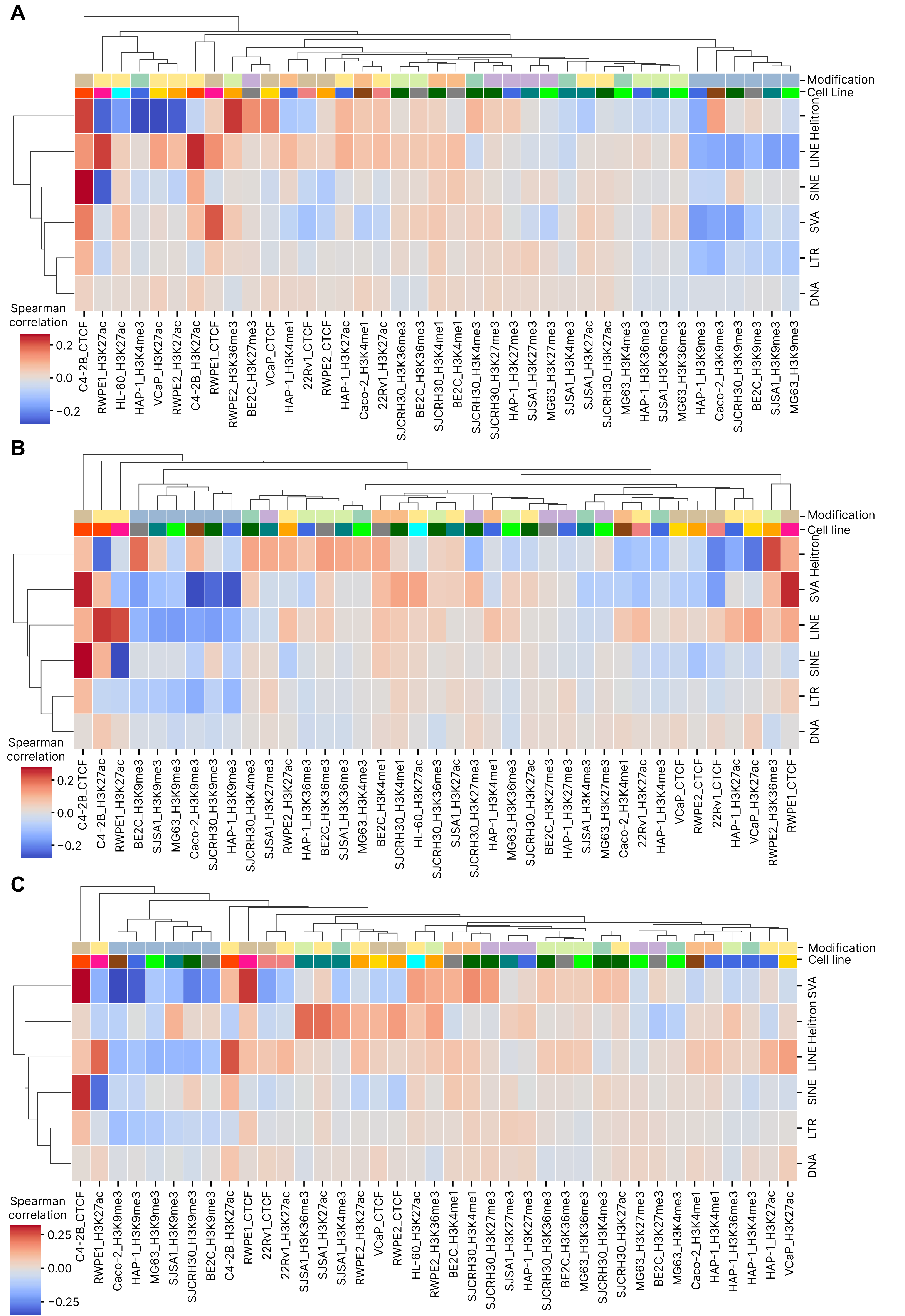

### Supplementary Figure 3

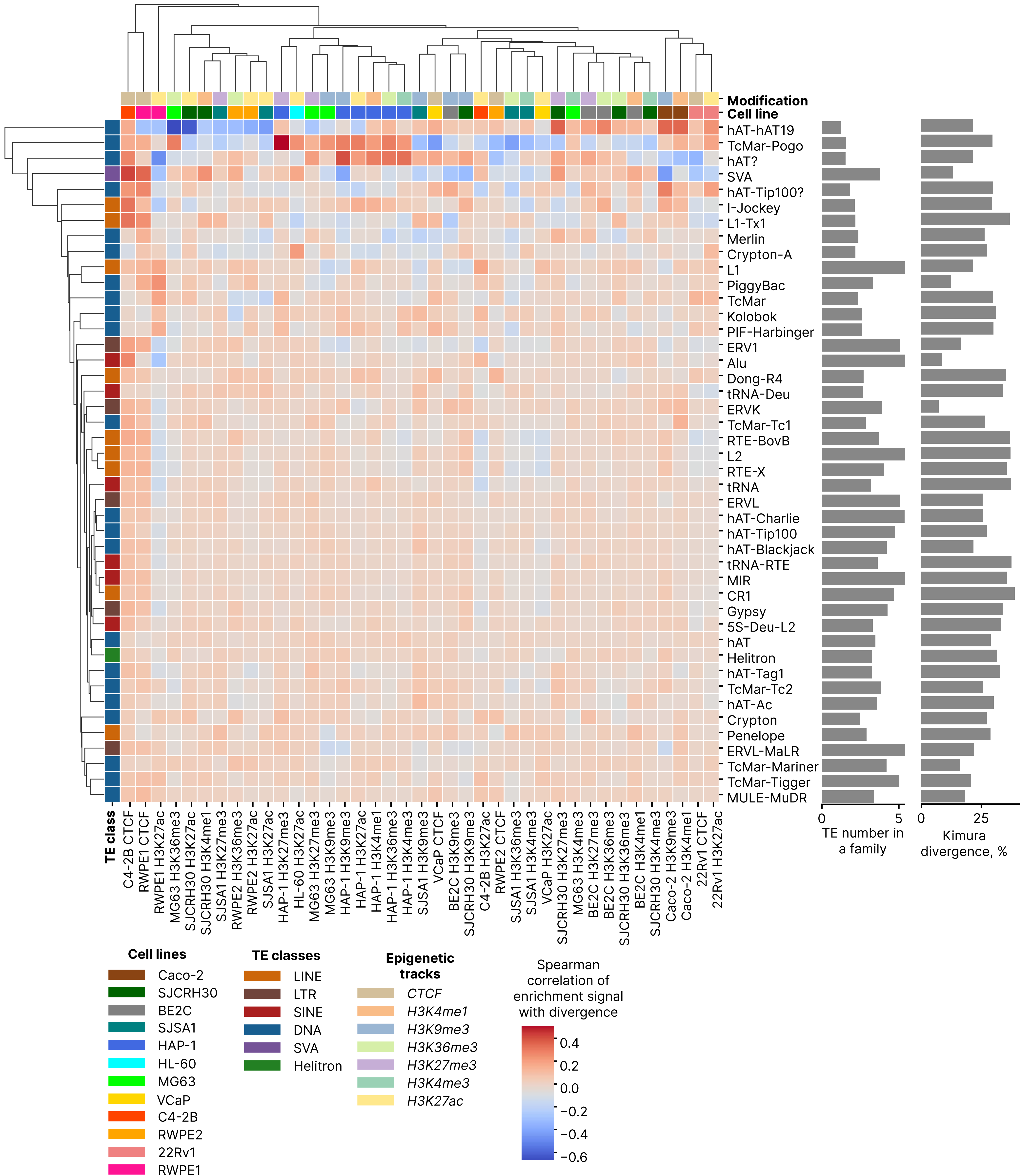

### Supplementary Figure 4

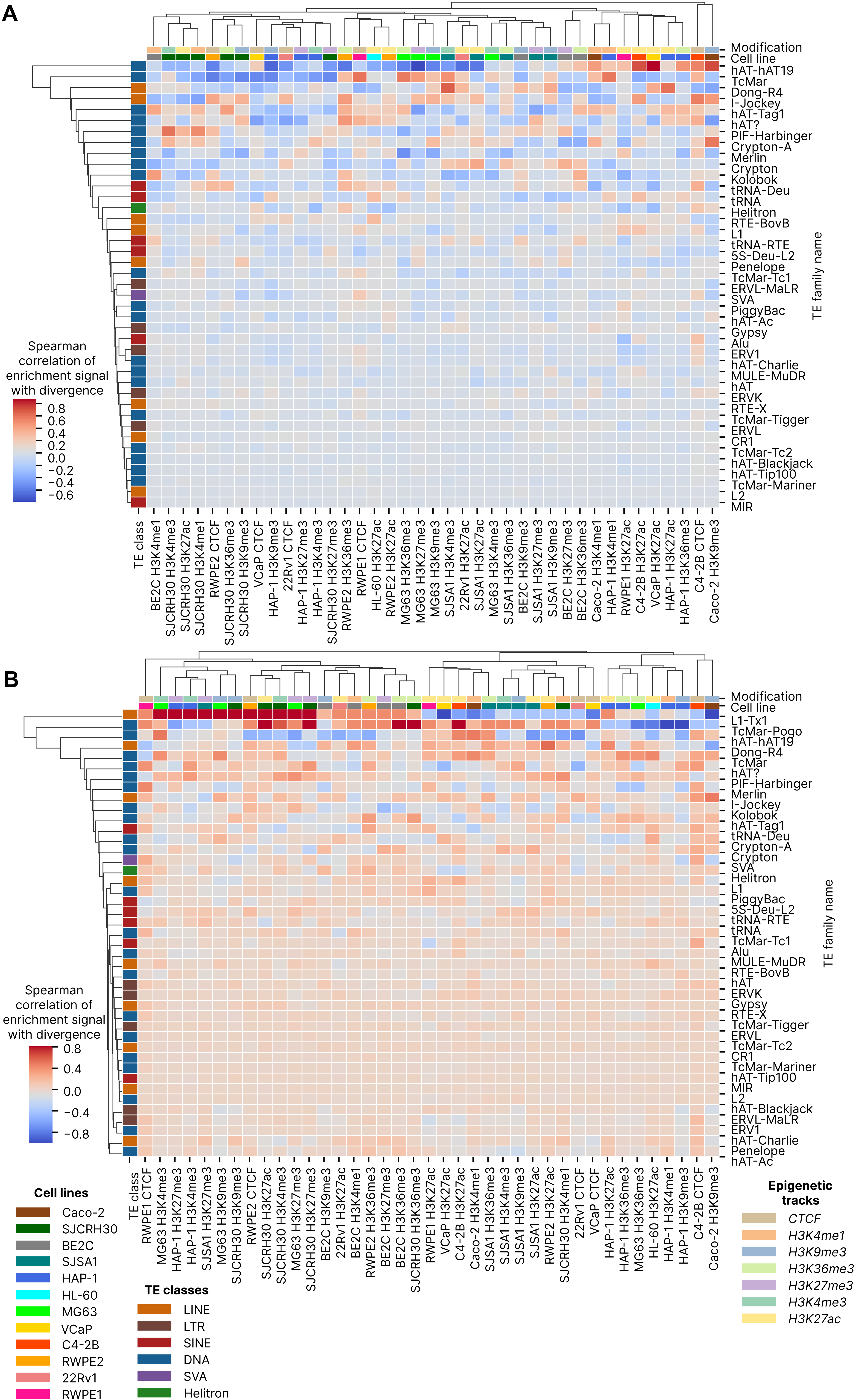

### Supplementary Figure 5

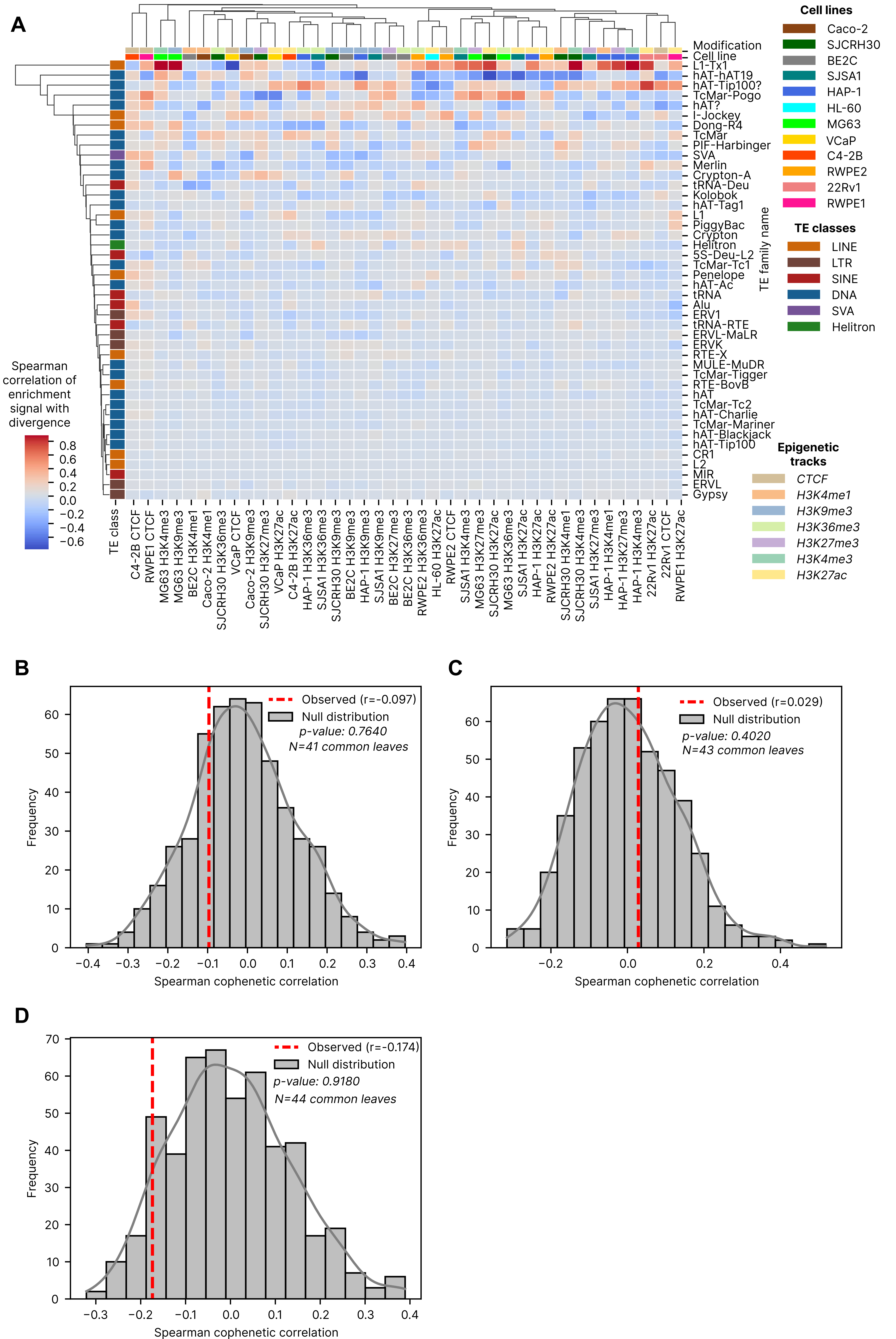

### Supplementary Figure 6

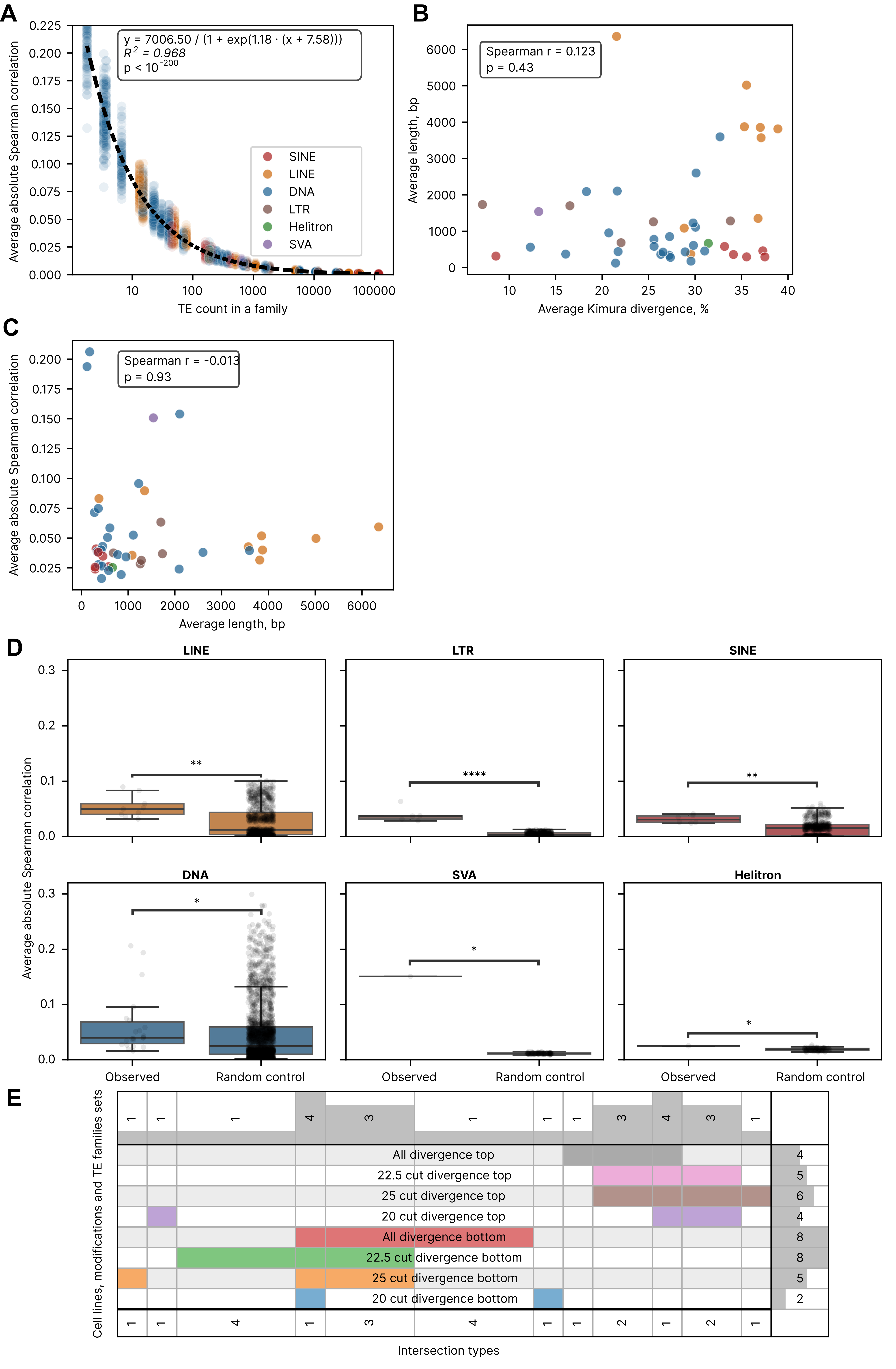

### Supplementary Figure 7

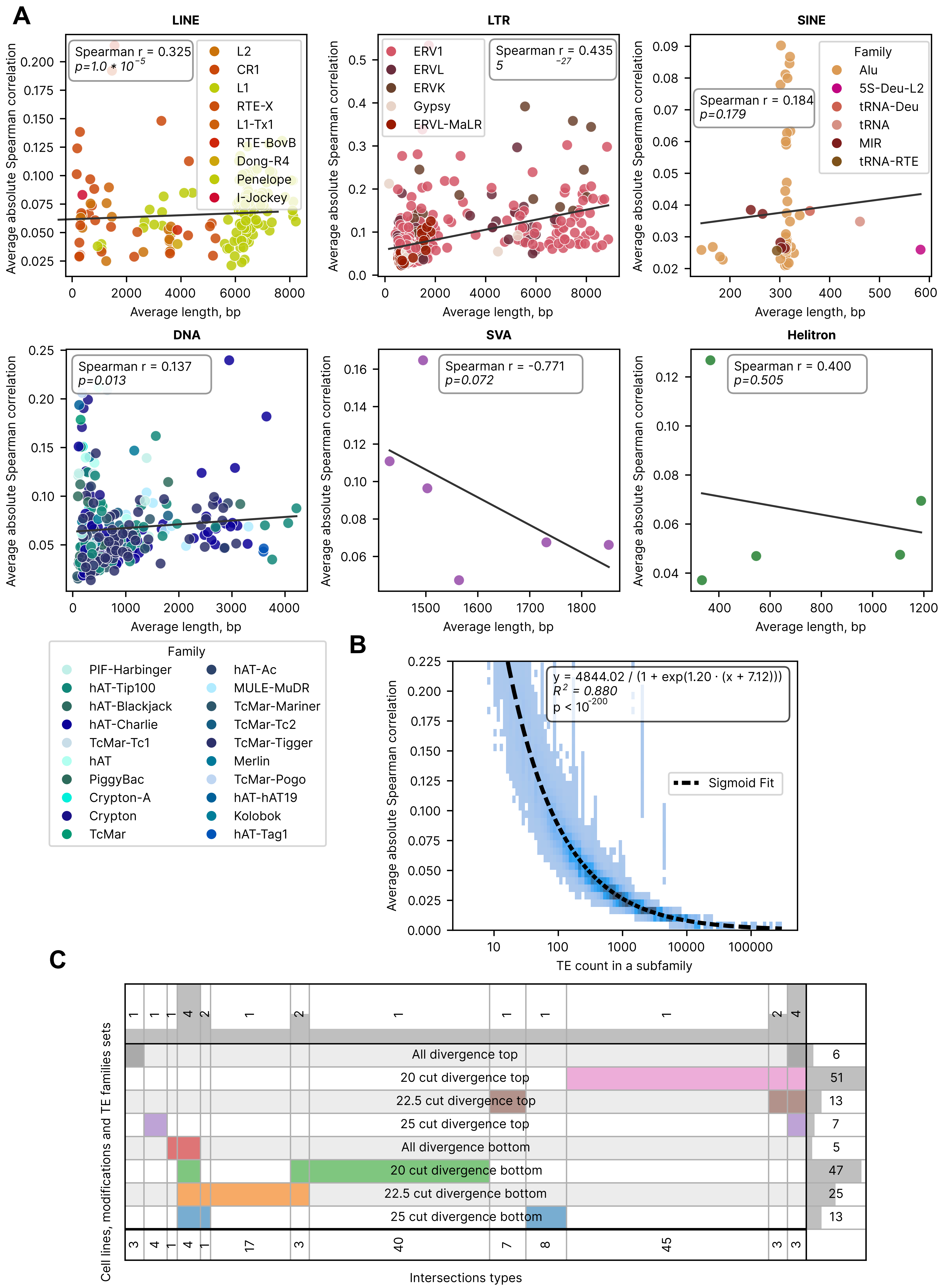

### Supplementary Figure 8

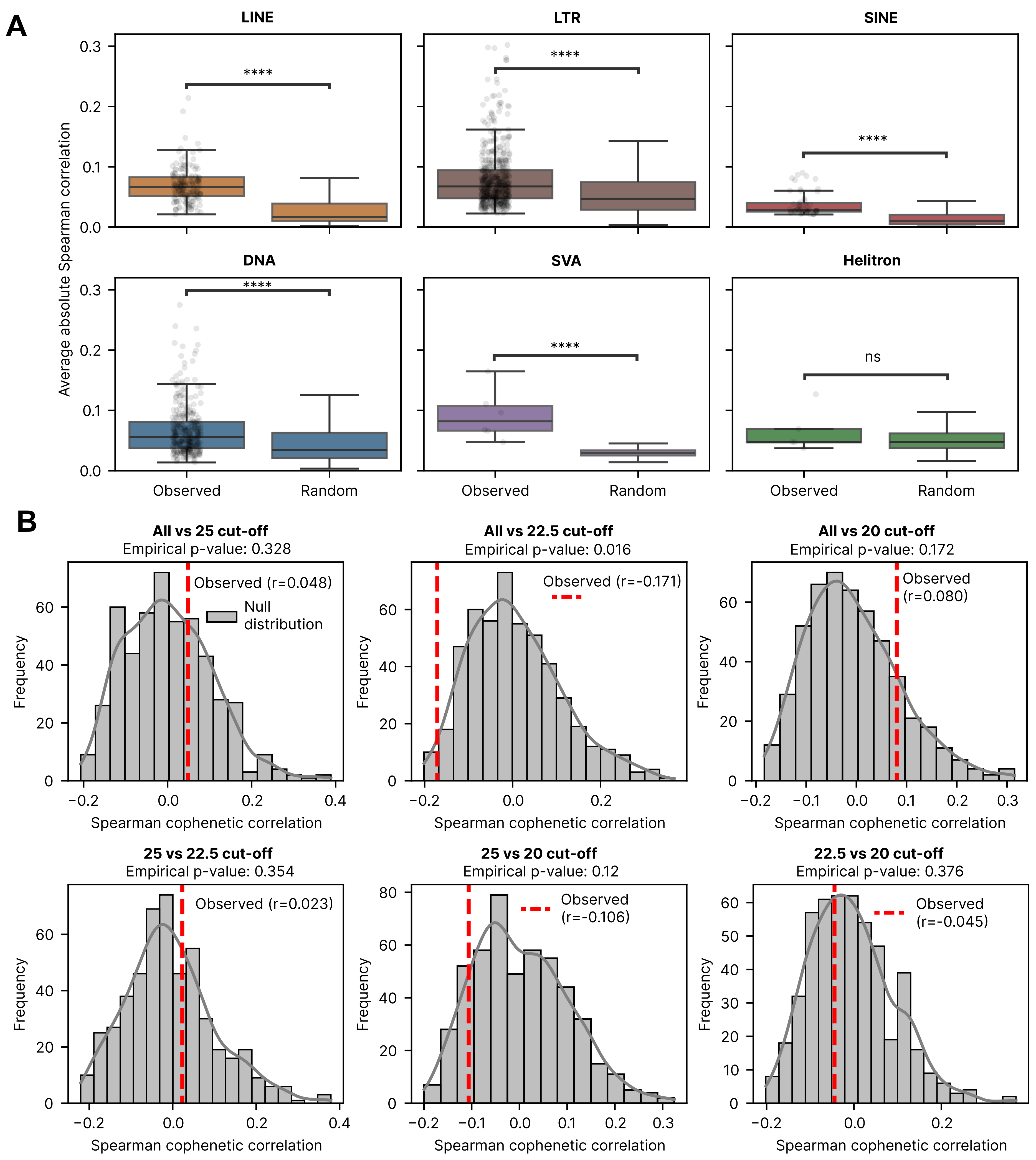

### Supplementary Figure 9

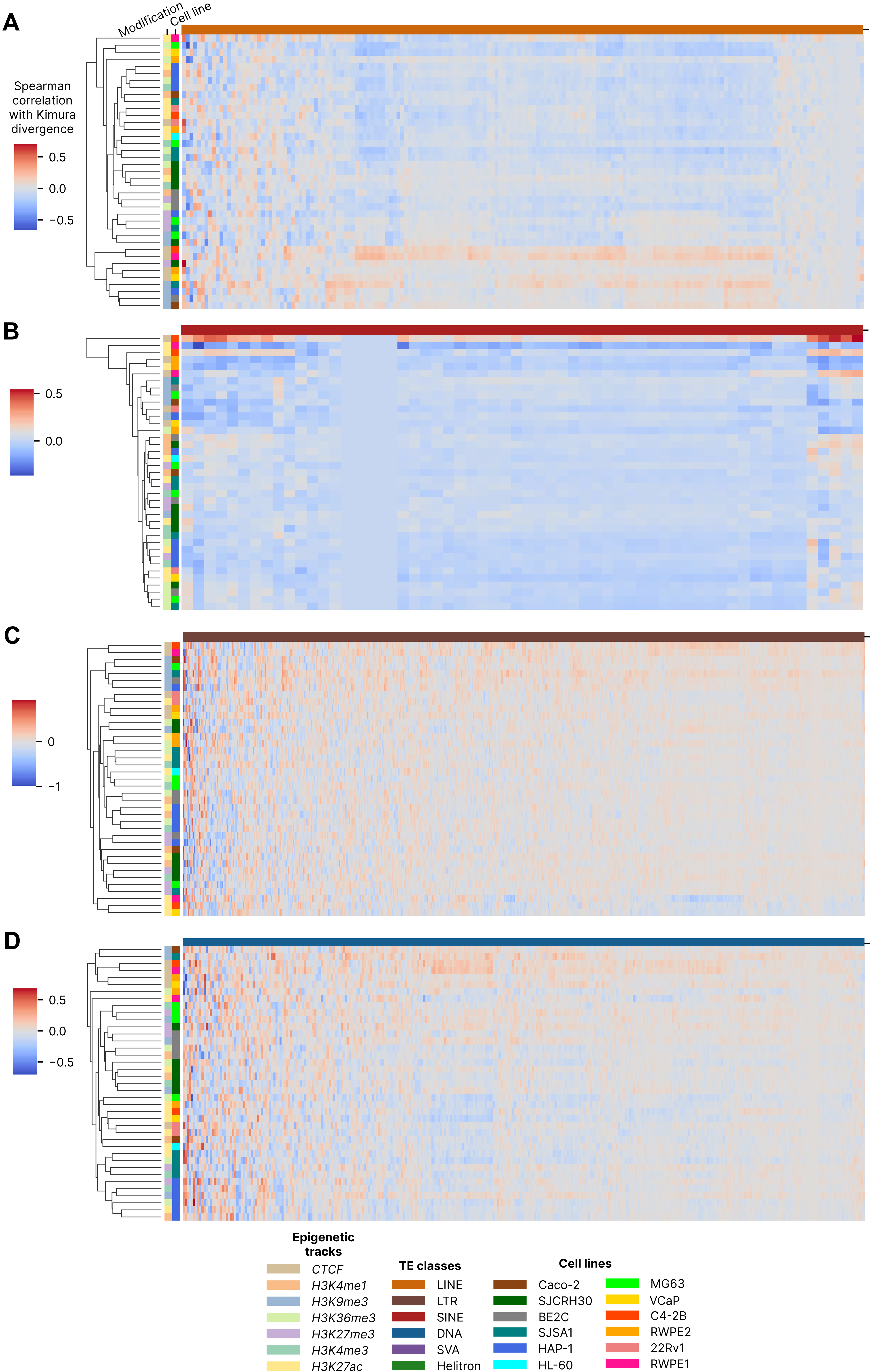
