## Supplementary File 3 for "Transposable element–host genome evolutionary arms race revealed by multi-modal epigenomic profiling in a telomere-to-telomere human genome reference": Supplementary File 3.ipynb

```
# %% [code]
import matplotlib
import matplotlib.pyplot as plt
import numpy as np
import pandas as pd
import seaborn as sns
from matplotlib.gridspec import GridSpec, GridSpecFromSubplotSpec
from scipy import stats
from tqdm import tqdm

plt.rcParams["svg.fonttype"] = "none"
plt.rcParams.update({'font.size': 10})

# %% [code]
plt.rcParams

# %% [markdown]
# Palettes

# %% [code]
cell_line_p = {
    "Caco-2": "saddlebrown",
    "SJCRH30": "darkgreen",
    "BE2C": "grey",
    "SJSA1": "teal",
    "HAP-1": "royalblue",
    "HL-60": "cyan",
    "MG63": "lime",
    "VCaP": "gold",
    "C4-2B": "orangered",
    "RWPE2": "orange",
    "22Rv1": "lightcoral",
    "RWPE1": "deeppink",
}
modification_p = {
    "CTCF": "#e5d8bd",
    "H3K4me1": "#fdcdac",
    "H3K9me3": "#b3cde3",
    "H3K36me3": "#e6f5c9",
    "H3K27me3": "#decbe4",
    "H3K4me3": "#b3e2cd",
    "H3K27ac": "#fff2ae",
}

modification_p_richer = {
    "CTCF": "#d3c09b",
    "H3K4me1": "#fabd88",
    "H3K9me3": "#9ab7d3",
    "H3K36me3": "#d7f0a8",
    "H3K27me3": "#c7aed6",
    "H3K4me3": "#9ad1b5",
    "H3K27ac": "#ffe98c",
}

class_names_p = {
    "LINE": "#cc660b",
    "LTR": "#70453c",
    "SINE": "#ab1f20",
    "DNA": "#195f90",
    "Retroposon": "#765297",
    "RC": "#238023",
}

# %% [code]


# %% [code]


# %% [markdown]
# Repeat masker table analysis

# %% [code]
repeat_masker = pd.read_csv("T2T_repeat_masker.csv")
splitted = repeat_masker["name"].str.split("#").str
repeat_masker["individual_name"] = splitted[0]
repeat_masker["family_name_agg"] = splitted[1]
repeat_masker["family_name_agg"][
    ~(repeat_masker["family_name_agg"].str.contains("/"))
] = (repeat_masker["family_name_agg"] + "/")
splitted_2 = repeat_masker["family_name_agg"].str.split("/").str
repeat_masker["family_name"] = splitted_2[1]
repeat_masker["class_name"] = splitted_2[0]
repeat_masker

# %% [code]
repeat_masker["class_name"].unique()

# %% [code]
repeat_masker["family_name"].unique()

# %% [code]
splitted = repeat_masker["name"].str.split("#").str
repeat_masker["individual_name"] = splitted[0]
repeat_masker["family_name"] = splitted[1]
repeat_masker

# %% [code]
repeat_masker['family_name'][!(repeat_masker['family_name'].str.contains('/'))] = repeat_masker['family_name'] + '/'

# %% [code]
repeat_masker = pd.read_csv("T2T_repeat_masker_processed.csv")

# %% [code]
repeat_masker.class_name.value_counts()

# %% [code]
class_query = 'class_name in ["LINE", "LTR", "SINE", "DNA", "Retroposon", "RC"]'
repeat_masker = repeat_masker.query(class_query)
repeat_masker.to_csv("T2T_repeat_masker_processed.csv")
repeat_masker

# %% [markdown]
## Divergence analysis

# %% [code]
repeat_masker = pd.read_csv("T2T_repeat_masker_processed.csv")

# %% [code]
# repeat_masker.to_csv("T2T_repeat_masker_processed.bed", sep="\t", header=False, index=False)

# %% [code]
# repeat_masker[['#"chrom"', 'chromStart', 'chromEnd', 'score', 'individual_name','family_name','class_name']].to_csv('T2T_repeat_masker_processed.csv')

# %% [code]
repeat_masker

# %% [code]
repeat_masker.score.max()

# %% [code]
(repeat_masker["class_name"] + " / " + repeat_masker["family_name"]).value_counts()

# %% [code]
pd.crosstab(repeat_masker["class_name"], repeat_masker["family_name"])

# %% [code]
repeat_masker[["class_name", "family_name"]].drop_duplicates().to_csv(
    "families_by_classes_TE.csv"
)

# %% [code]
repeat_masker[["class_name", "individual_name"]].drop_duplicates().to_csv(
    "individuals_by_classes_TE.csv"
)

# %% [code]
repeat_masker["individual_name"].value_counts().to_csv(
    "individuals_counts.csv", index=True
)
repeat_masker["family_name"].value_counts().to_csv("families_counts.csv", index=True)

# %% [code]


# %% [code]
class_names = ["LINE", "LTR", "SINE", "DNA", "Retroposon", "RC"]

figure, axs = plt.subplots(
    nrows=2,
    ncols=3,
    sharex=True,
    sharey=False,
    squeeze=True,
    # width_ratios=None,
    # height_ratios=[1, 1, 1, 1, 1, 1, 0.3, 0.3],
    figsize=(28, 16),
)

# Use zip to iterate over class names and the flattened axes array
# axs.flatten() converts the 2x3 array into a 1D array of 6 axes
for class_name, ax in zip(class_names, axs.flatten()):

    # Filter the data
    query_local = f'class_name == "{class_name}"'
    repeat_masker_local = repeat_masker.query(query_local)

    # Use sns.histplot (axes-level) instead of sns.displot (figure-level)
    sns.histplot(repeat_masker_local.score, ax=ax, bins=100)

    # Use ax.set_title() to set the title for the subplot
    ax.set_title(class_name)

# Save the final figure
# plt.savefig("plots/Divergence_score_dist_by_class.svg")

# %% [code]
g = sns.FacetGrid(repeat_masker, col="class_name")
p1 = g.map(sns.distplot, "score", hist=False, rug=True).add_legend()

# %% [markdown]
### Concordance between divergence and evolutionary age

# %% [code]
from supervenn import supervenn

# %% [code]
individuals_by_classes = pd.read_csv("individuals_by_classes_TE.csv").dropna()

repeat_masker = pd.read_csv("T2T_repeat_masker_processed.csv")
divergence_average = (
    repeat_masker[["score", "individual_name"]]
    .groupby("individual_name")
    .mean()["score"]
)


public_age = pd.read_excel('41598_2024_73752_MOESM2_ESM.xlsx', sheet_name='TE subfamily')  
individuals_by_classes = individuals_by_classes.set_index("individual_name").reindex(
    public_age['repeat subfamily name']
)

public_age['average_divergence'] = list(divergence_average.reindex(public_age['repeat subfamily name']))
public_age['class_name'] = list(individuals_by_classes['class_name'])
public_age['class_name_random'] = public_age['class_name'].sample(frac=1, random_state=1).set_axis(public_age.index)
public_age.dropna(subset='average_divergence')

# %% [code]
public_age['class_name']

# %% [code]
public_age['class_name'].sample(frac=1, random_state=1).reset_index(drop=True)

# %% [code]
from scipy import stats
public_age_young = public_age[public_age['evolutionary age'] < 150][['evolutionary age','average_divergence']].dropna()

stats.pearsonr(public_age_young['evolutionary age'], public_age_young['average_divergence'])


# %% [code]
import matplotlib.pyplot as plt
from matplotlib_venn import venn2, venn2_circles
import pandas as pd
import numpy as np
import os


divergence_average = (
    repeat_masker[["score", "family_name"]]
    .groupby("family_name")
    .mean()["score"]
)


# --- 2. Define your sets and labels ---
sets = [
    set(divergence_average.index), 
    set(public_age['repeat family name'])
]
labels = ('Current study, T2T genome', 'Kosuge et al., 2024, GRCh38 genome') # Use a tuple for labels

# --- 3. Create the plot with configurable figsize ---
print("INFO: Generating Venn diagram...")
# You can change the figsize here
fig, ax = plt.subplots(figsize=(8, 6))

# --- 4. Plot the Venn diagram ---
v = venn2(
    subsets=sets,
    set_labels=labels,
    ax=ax,
    # Customize colors
    set_colors=('#1f77b4', '#ff7f0e'), # Blue and Orange
    alpha=0.7
)

# Add circles with black outlines
venn2_circles(subsets=sets, linestyle='solid', linewidth=1.0, color='black', ax=ax)

# Customize label fonts
for text in v.set_labels:
    text.set_fontsize(10)
for text in v.subset_labels:
    text.set_fontsize(10)

# Add a title
ax.set_title("Overlap of TE Names", fontsize=16)

# --- 5. Save to SVG ---
save_path = "plots/TE_name_overlap_families.svg"
print(f"INFO: Saving plot to {save_path}...")

# Ensure the directory exists
os.makedirs(os.path.dirname(save_path), exist_ok=True)

# Save with tight_layout and transparent background
fig.savefig(
    save_path, 
    format="svg", 
    bbox_inches="tight",
    transparent=True
)

plt.show()
print("SUCCESS: Plot saved.")

# %% [code]
sets = [set(divergence_average.index), set(public_age['repeat subfamily name'])]
labels = ['individual name', 'repeat subfamily name']
supervenn(sets, labels)

# %% [code]
set(divergence_average.index) - set(public_age['repeat subfamily name'])

# %% [code]
set(public_age['repeat subfamily name']) - set(divergence_average.index)

# %% [code]
public_age = pd.read_excel('41598_2024_73752_MOESM2_ESM.xlsx', sheet_name='TE subfamily')
public_age[['repeat class', 'repeat copy count']].groupby('repeat class').sum()

# %% [markdown]
# Gene table analysis

# %% [code]
gene_table = pd.read_csv("T2T_refseq_all_genes.csv")
gene_table = gene_table[
    (gene_table.geneName.str.startswith("NM"))
    | (gene_table.geneName.str.startswith("NR"))
    | (gene_table.geneName.str.startswith("NP"))
    | (gene_table.geneName.str.startswith("NP"))
]
gene_table["thick_length"] = gene_table["thickEnd"] - gene_table["thickStart"]
gene_table["TSS_start"] = gene_table["chromStart"] - 5000
gene_table["TSS_end"] = gene_table["chromStart"] + 5000

gene_table["TSS_start"][gene_table["strand"] == "-"] = (
    gene_table["chromEnd"][gene_table["strand"] == "-"] - 5000
)
gene_table["TSS_end"][gene_table["strand"] == "-"] = (
    gene_table["chromEnd"][gene_table["strand"] == "-"] + 5000
)

gene_table[['#"chrom"', "TSS_start", "TSS_end", "geneName2"]].to_csv(
    "TSS.bed", sep="\t", header=False, index=False
)

gene_table["TSS_start"][gene_table.TSS_start < 0] = 0

gene_table  # [(gene_table['thick_length'] > 3000) & (gene_table['strand'] == '+')]
# gene_table[(gene_table['thick_length'] > 3000) & (gene_table['strand'] == '+')]

# %% [code]
gene_table[gene_table.TSS_start < 0]

# %% [code]
gene_table[['#"chrom"', "TSS_start", "TSS_end", "geneName2"]].drop_duplicates().to_csv(
    "T2T_genes.bed", index=False, header=False, sep="\t"
)

# %% [code]
len(gene_table.geneName2.unique())

# %% [code]
gene_table.type.unique()

# %% [code]
sns.histplot(gene_table["thick_length"])

# %% [markdown]
# Analysis of epigenetic profiles mapped on retroelements

# %% [markdown]
## Analysis of a sample bedGraph file

# %% [code]
sample_mapping = pd.read_csv(
    "22Rv1.CTCF.chm13v2.0.sample.bedGraph", sep="\t", header=None
)
sample_mapping.columns = ["Chromosome", "Start", "End", "Signal"]
sample_mapping

# %% [code]
sns.histplot(sample_mapping.Signal)

# %% [markdown]
## Analysis of sample mapping on retroelements

# %% [code]
sample_enrichment = pd.read_csv(
    "./epigenomic_files/RWPE1.H3K27ac.chm13v2.0.mapped_on_repeat_masker.bedGraph",
    sep="\t",
    header=None,
)
sample_enrichment.columns = [
    "Chromosome",
    "Start",
    "End",
    "Divergence",
    "Name",
    "Family",
    "Class",
    "Signal",
]
sample_enrichment["Signal"] = pd.to_numeric(
    sample_enrichment["Signal"], errors="coerce"
)
sample_enrichment["Divergence"] = pd.to_numeric(
    sample_enrichment["Divergence"], errors="coerce"
)

sample_enrichment

# %% [code]
sample_enrichment["Signal"].mean()

# %% [code]
sample_enrichment = pd.read_csv(
    "22Rv1.CTCF.chm13v2.0.repeats_mapped.bed", sep="\t", header=None
)
sample_enrichment.columns = [
    "Chromosome",
    "Start",
    "End",
    "Divergence",
    "Name",
    "Family",
    "Class",
    "Signal",
]
sample_enrichment["Signal"] = pd.to_numeric(
    sample_enrichment["Signal"], errors="coerce"
)
sample_enrichment["Divergence"] = pd.to_numeric(
    sample_enrichment["Divergence"], errors="coerce"
)

sample_enrichment

# %% [code]
a = sample_enrichment[["Class", "Signal"]].groupby("Class").mean()["Signal"].to_dict()
a.update({"a": 1, "b": 2})
a

# %% [code]
1

# %% [code]
# non_zero_query = 'Signal != 0'
# sample_enrichment_non_zero = sample_enrichment.query(non_zero_query)

sns.histplot(sample_enrichment.Signal)

# %% [markdown]
This image displays a leptokurtic (sharply peaked, heavy-tailed) and slightly right-skewed distribution. The vast majority of the "Signal" values are centered at 0, with long tails of less frequent values on both the positive and negative sides.This type of distribution is most likely derived from a Log-fold Change (logFC) calculation, a standard normalization method for epigenomic enrichment data.What the Distribution Shape MeansThe Peak at 0: This sharp, massive peak represents the genomic "background." It's the vast majority of the genome where there is no significant difference in the signal (e.g., no enrichment or depletion) compared to the control.The Right Tail (Positive Values): This tail represents regions of enrichment. These are the areas where the epigenomic mark (like a histone modification or transcription factor binding) is significantly more present than in the control sample.The Left Tail (Negative Values): This tail represents regions of depletion, where the mark is significantly less present than in the control.The Skew: The right tail extends further than the left tail, which is common for "enrichment" tracks. It indicates that the high-enrichment signals are stronger (further from 0) than the depletion signals.How This Normalization is DerivedThis "Signal" value is almost certainly a log-fold change (or a very similar log-ratio), which normalizes a "treatment" sample against a "control" sample.The calculation is typically:$$Signal = \log_{2} \left( \frac{\text{Signal in Treatment Sample}}{\text{Signal in Control Sample}} \right)$$Here's how this calculation creates the distribution you see:If Treatment = Control: (No change) The ratio is 1. $\log_{2}(1) = 0$. This creates the massive peak at 0.If Treatment > Control: (Enrichment) The ratio is > 1. $\log_{2}(>1)$ gives a positive value. This creates the right tail.If Treatment < Control: (Depletion) The ratio is < 1. $\log_{2}(<1)$ gives a negative value. This creates the left tail.This log-ratio is the most common way to represent "enrichment" data from sources like ChIP-seq or ATAC-seq because it provides a single, intuitive metric centered at zero, showing both the direction (enrichment/depletion) and magnitude of the change.Note: A related possibility is a log-likelihood ratio, which also produces positive and negative values centered around zero and is used in some signal-processing pipelines (like MACS2). However, log-fold change is the most straightforward and common interpretation.

# %% [code]
sns.jointplot(data=sample_enrichment, x="Divergence", y="Signal", kind="hex")

# %% [markdown]
### Good plot of Signal vs Divergence correlation by classes of TE

# %% [code]
sample_enrichment

# %% [code]
import matplotlib.pyplot as plt
import numpy as np
import pandas as pd
import seaborn as sns
from matplotlib.gridspec import GridSpec, GridSpecFromSubplotSpec

# --- Main Plotting Logic ---
figure = plt.figure(figsize=(16, 10))
# Create the main 2x3 grid for the entire figure
outer_grid = GridSpec(2, 3, figure=figure, wspace=0.2, hspace=0.2)
class_names = ["LINE", "LTR", "SINE", "DNA", "Retroposon", "RC"]

for i, class_name in enumerate(class_names):
    print(class_name)
    # For each cell in the outer grid, create a nested GridSpec for the jointplot
    inner_grid = GridSpecFromSubplotSpec(
        2,
        2,
        subplot_spec=outer_grid[i],
        width_ratios=[4, 1],
        height_ratios=[1, 4],
        wspace=0.05,
        hspace=0.05,
    )

    # Create the axes for the main plot and the two marginal plots
    ax_main = figure.add_subplot(inner_grid[1, 0])
    ax_hist_x = figure.add_subplot(inner_grid[0, 0], sharex=ax_main)
    ax_hist_y = figure.add_subplot(inner_grid[1, 1], sharey=ax_main)

    # Filter the data for the current class
    query_local = f'Class == "{class_name}"'
    sample_enrichment_local = sample_enrichment.query(query_local)

    # --- Plot the data on the created axes ---
    # 1. Main hexbin plot (axes-level equivalent of jointplot's kind="hex")
    sns.histplot(
        data=sample_enrichment_local,
        x="Divergence",
        y="Signal",
        bins=50,
        pmax=0.9,
        cmap="viridis",
        ax=ax_main,
    )

    # 2. Top marginal histogram for 'Divergence'
    sns.histplot(data=sample_enrichment_local, x="Divergence", ax=ax_hist_x, bins=50)

    # 3. Right marginal histogram for 'Signal'
    sns.histplot(data=sample_enrichment_local, y="Signal", ax=ax_hist_y, bins=50)

    # --- Tidy up the plot ---
    # Remove ticks and labels from the marginal plots for a cleaner look
    ax_hist_x.tick_params(axis="x", labelbottom=False)
    ax_hist_x.tick_params(axis="y", labelbottom=False)
    ax_hist_x.set_ylabel("")
    ax_hist_y.tick_params(axis="y", labelleft=False)
    ax_hist_y.tick_params(axis="x", labelleft=False)
    ax_hist_y.set_xlabel("")

    corr_coef = np.corrcoef(
        sample_enrichment_local.dropna()["Divergence"],
        sample_enrichment_local.dropna()["Signal"],
    )[0, 1]
    corr_coef = str(round(corr_coef, 3))

    # Set the title for this entire "jointplot" cell
    ax_hist_x.set_title(class_name + f", r={corr_coef}", fontsize=16, pad=10)

# Adjust layout to prevent labels from overlapping
plt.suptitle("Divergence vs. Signal by Repeat Class", fontsize=20, y=0.97)
figure.tight_layout(rect=[0, 0, 1, 0.95])  # Adjust rect to make space for suptitle

# Save the final figure
# plt.savefig("plots/Divergence_vs_Signal_grid.svg")
plt.show()

# %% [code]
import matplotlib.pyplot as plt
import numpy as np
import pandas as pd
import seaborn as sns
from matplotlib.gridspec import GridSpec, GridSpecFromSubplotSpec

# --- Main Plotting Logic ---
figure = plt.figure(figsize=(16, 10))
class_names_grid = np.array(class_names).reshape((2, 3))

# Create the main 2x3 grid for the entire figure
outer_grid = GridSpec(2, 3, figure=figure, wspace=0.05, hspace=0.1)

# --- Store references to the first axis in each row/column ---
shared_x_axes = [None] * 3  # One for each column
shared_y_axes = [None] * 2  # One for each row

for r in range(2):
    for c in range(3):
        class_name = class_names_grid[r, c]
        print(f"Plotting: {class_name} (Row {r}, Col {c})")

        # For each cell in the outer grid, create a nested GridSpec
        inner_grid = GridSpecFromSubplotSpec(
            2,
            2,
            subplot_spec=outer_grid[r, c],
            width_ratios=[5, 1],
            height_ratios=[1, 5],
            wspace=0.05,
            hspace=0.05,
        )

        # --- Create axes with sharing logic ---
        ax_main = figure.add_subplot(
            inner_grid[1, 0], sharex=shared_x_axes[c], sharey=shared_y_axes[r]
        )
        ax_hist_x = figure.add_subplot(inner_grid[0, 0], sharex=ax_main)
        ax_hist_y = figure.add_subplot(inner_grid[1, 1], sharey=ax_main)

        # --- Store the first axis of each row/col as reference ---
        if r == 0:
            shared_x_axes[c] = ax_main
        if c == 0:
            shared_y_axes[r] = ax_main

        # Filter the data for the current class
        query_local = f'Class == "{class_name}"'
        sample_enrichment_local = sample_enrichment.query(query_local)

        # --- Plot the data ---
        sns.histplot(
            data=sample_enrichment_local,
            x="Divergence",
            y="Signal",
            bins=48,
            pmax=0.9,
            cmap="viridis",
            ax=ax_main,
        )
        sns.histplot(
            data=sample_enrichment_local, x="Divergence", ax=ax_hist_x, bins=50
        )
        sns.histplot(data=sample_enrichment_local, y="Signal", ax=ax_hist_y, bins=50)

        # --- START: MODIFIED TIDY-UP BLOCK ---
        # Tidy up: Remove all ticks, labels, and borders from marginal plots

        # Top histogram (ax_hist_x)
        ax_hist_x.set_ylabel("")
        ax_hist_x.set_xlabel("")
        ax_hist_x.tick_params(
            axis="both", bottom=False, left=False, labelbottom=False, labelleft=False
        )
        for spine in ax_hist_x.spines.values():
            spine.set_visible(False)

        # Right histogram (ax_hist_y)
        ax_hist_y.set_xlabel("")
        ax_hist_y.set_ylabel("")
        ax_hist_y.tick_params(
            axis="both", bottom=False, left=False, labelbottom=False, labelleft=False
        )
        for spine in ax_hist_y.spines.values():
            spine.set_visible(False)
        # --- END: MODIFIED TIDY-UP BLOCK ---

        # --- Tidy up: Main plot ticks/labels (for shared axes) ---
        if c > 0:  # Not in the first column
            ax_main.tick_params(axis="y", labelleft=False)
            ax_main.set_ylabel("")
        else:
            ax_main.set_ylabel("Signal", fontsize=12)  # Only show Y label on first col

        if r < 1:  # Not in the bottom row
            ax_main.tick_params(axis="x", labelbottom=False)
            ax_main.set_xlabel("")
        else:
            ax_main.set_xlabel(
                "Divergence", fontsize=12
            )  # Only show X label on bottom row

        # Calculate correlation
        clean_data = sample_enrichment_local.dropna(subset=["Divergence", "Signal"])
        if len(clean_data) > 1:
            corr_coef = np.corrcoef(clean_data["Divergence"], clean_data["Signal"])[
                0, 1
            ]
            corr_str = f"r={corr_coef:.3f}"
        else:
            corr_str = "r=N/A"

        # Set the title for this "jointplot" cell
        ax_hist_x.set_title(f"{class_name}, {corr_str}", fontsize=16, pad=10)

# Adjust layout to prevent labels from overlapping
plt.suptitle("Divergence vs. Signal by Repeat Class (Shared Axes)", fontsize=20, y=0.98)
figure.tight_layout(rect=[0, 0.03, 1, 0.95])  # Adjust rect to make space for suptitle

# Save the final figure
# plt.savefig("plots/Divergence_vs_Signal_grid_shared_axes.svg")
plt.show()

# %% [markdown]
## Build distplots of the TE mapping by modification and cell line

# %% [code]
import os

import matplotlib.pyplot as plt
import numpy as np
import pandas as pd
import seaborn as sns
from tqdm import tqdm  # For progress bars

# Removed unused imports: GridSpec, GridSpecFromSubplotSpec

plt.rcParams["svg.fonttype"] = "none"


# --- Helper Class for Colored Logging ---
class Logger:
    """A simple logger for pretty, colored console output."""

    BLUE = "\033[94m"
    GREEN = "\033[92m"
    YELLOW = "\033[93m"
    ENDC = "\033[0m"

    @staticmethod
    def info(message):
        print(f"{Logger.BLUE}INFO: {message}{Logger.ENDC}")

    @staticmethod
    def success(message):
        print(f"{Logger.GREEN}SUCCESS: {message}{Logger.ENDC}")

    @staticmethod
    def warning(message):
        print(f"{Logger.YELLOW}WARNING: {message}{Logger.ENDC}")

    @staticmethod
    def step(message):
        # Use tqdm.write() to print from within a loop
        tqdm.write(f"  -> {message}")


cell_lines = [
    "Caco-2",
    "SJCRH30",
    "BE2C",
    "SJSA1",
    "HAP-1",
    "HL-60",
    "MG63",
    "VCaP",
    "C4-2B",
    "RWPE2",
    "22Rv1",
    "RWPE1",
]
modifications = [
    "CTCF",
    "H3K27ac",
    "H3K4me1",
    "H3K9me3",
    "H3K36me3",
    "H3K27me3",
    "H3K4me3",
    # "H3K27ac",
]
class_names = ["LINE", "LTR", "SINE", "DNA", "Retroposon", "RC"]


figure, axs = plt.subplots(
    nrows=8,
    ncols=5,
    sharex=False,
    sharey=False,
    squeeze=True,
    figsize=(20, 24),  # Adjusted figsize for the larger grid
)

# --- CORRECTED: Create an iterator from the flattened axes list ---
axes_iter = iter(axs.flatten())

# --- Main Script ---
Logger.info(f"Starting batch plot generation for the 39 plots...")
correlation_df_list = []
# Use tqdm for the outer loop progress bar
for cell_line in tqdm(cell_lines, desc="Processing Cell Lines"):

    # Use a nested tqdm for the inner loop
    for modification in tqdm(
        modifications, desc=f"  Mods for {cell_line}", leave=False
    ):

        filepath = f"./epigenomic_files/{cell_line}.{modification}.chm13v2.0.mapped_on_repeat_masker.bedGraph"
        # --- Data Loading ---
        # Use Logger.step (which uses tqdm.write) for in-loop logging
        Logger.step(f"Loading data from: {filepath}")
        try:
            sample_enrichment = pd.read_csv(
                filepath,
                sep="\t",
                header=None,
            )
            # Logger.success("File loaded successfully.") # Too noisy for a loop
        except FileNotFoundError:
            Logger.warning(f"File not found, skipping: {filepath}")
            continue  # Skip to the next modification
        except Exception as e:
            Logger.warning(
                f"An unexpected error occurred while reading {filepath}: {e}"
            )
            continue

        sample_enrichment.columns = [
            "Chromosome",
            "Start",
            "End",
            "Divergence",
            "Name",
            "Family",
            "Class",
            "Signal",
        ]
        sample_enrichment["Signal"] = pd.to_numeric(
            sample_enrichment["Signal"], errors="coerce"
        )

        # --- CORRECTED: Indentation and Iterator ---
        # Get the next available axis
        try:
            ax = next(axes_iter)
        except StopIteration:
            # This should not happen if grid is sized correctly, but it's safe
            Logger.warning("Ran out of axes, stopping early.")
            break

        # Use sns.histplot (axes-level)
        sns.histplot(sample_enrichment["Signal"].dropna(), ax=ax, bins=100)

        # --- CORRECTED: Use the correct variables for the title ---
        ax.set_title(f"{cell_line}\n{modification}", fontsize=8)

# Clean up any remaining empty axes at the end
# This loop will turn off axes that were never used
while True:
    try:
        ax = next(axes_iter)
        ax.axis("off")
    except StopIteration:
        break

figure.tight_layout(rect=[0, 0.03, 1, 0.95])
plt.savefig("plots/Signal_distribution_all_samples.svg")
Logger.success("All plots saved.")
plt.show()

# %% [markdown]
## Build good plots for all marks and cell lines and record pearson correlation coefficient

# %% [code]
cell_lines = [
    "Caco-2",
    "SJCRH30",
    "BE2C",
    "SJSA1",
    "HAP-1",
    "HL-60",
    "MG63",
    "VCaP",
    "C4-2B",
    "RWPE2",
    "22Rv1",
    "RWPE1",
]
modifications = [
    "CTCF",
    "H3K4me1",
    "H3K9me3",
    "H3K36me3",
    "H3K27me3",
    "H3K4me3",
    "H3K27ac",
]
class_names = ["LINE", "LTR", "SINE", "DNA", "Retroposon", "RC"]

# %% [code]
import os

import matplotlib.pyplot as plt
import numpy as np
import pandas as pd
import seaborn as sns
from matplotlib.gridspec import GridSpec, GridSpecFromSubplotSpec
from tqdm import tqdm  # For progress bars

plt.rcParams["svg.fonttype"] = "none"


# --- Helper Class for Colored Logging ---
class Logger:
    """A simple logger for pretty, colored console output."""

    BLUE = "\033[94m"
    GREEN = "\033[92m"
    YELLOW = "\033[93m"
    ENDC = "\033[0m"

    @staticmethod
    def info(message):
        print(f"{Logger.BLUE}INFO: {message}{Logger.ENDC}")

    @staticmethod
    def success(message):
        print(f"{Logger.GREEN}SUCCESS: {message}{Logger.ENDC}")

    @staticmethod
    def warning(message):
        print(f"{Logger.YELLOW}WARNING: {message}{Logger.ENDC}")

    @staticmethod
    def step(message):
        print(f"  -> {message}")


cell_lines = [
    "Caco-2",
    "SJCRH30",
    "BE2C",
    "SJSA1",
    "HAP-1",
    "HL-60",
    "MG63",
    "VCaP",
    "C4-2B",
    "RWPE2",
    "22Rv1",
    "RWPE1",
]
modifications = [
    "CTCF",
    "H3K27ac",
    "H3K4me1",
    "H3K9me3",
    "H3K36me3",
    "H3K27me3",
    "H3K4me3",
    #    "H3K27ac",
]
class_names = ["LINE", "LTR", "SINE", "DNA", "Retroposon", "RC"]

# --- Main Script ---
Logger.info("Starting batch plot generation...")
correlation_df_list = []
# Use tqdm for the outer loop progress bar
for cell_line in tqdm(cell_lines, desc="Processing Cell Lines"):

    # Use a nested tqdm for the inner loop
    for modification in tqdm(
        modifications, desc=f"  Mods for {cell_line}", leave=False
    ):

        filepath = f"./epigenomic_files/{cell_line}.{modification}.chm13v2.0.mapped_on_repeat_masker.bedGraph"
        save_path_png = f"plots/Epigenetic_signal_vs_divergence_by_class/Divergence_vs_Signal_by_classes_{cell_line}_{modification}_tidy.png"
        save_path_svg = f"plots/Epigenetic_signal_vs_divergence_by_class/Divergence_vs_Signal_by_classes_{cell_line}_{modification}_tidy.svg"
        # --- Data Loading ---
        Logger.step(f"Loading data from: {filepath}")
        try:
            sample_enrichment = pd.read_csv(
                filepath,
                sep="\t",
                header=None,
            )
            Logger.success("File loaded successfully.")
        except FileNotFoundError:
            Logger.warning(f"File not found, skipping: {filepath}")
            continue  # Skip to the next modification
        except Exception as e:
            Logger.warning(
                f"An unexpected error occurred while reading {filepath}: {e}"
            )
            continue

        sample_enrichment.columns = [
            "Chromosome",
            "Start",
            "End",
            "Divergence",
            "Name",
            "Family",
            "Class",
            "Signal",
        ]
        sample_enrichment["Signal"] = pd.to_numeric(
            sample_enrichment["Signal"], errors="coerce"
        )
        sample_enrichment["Divergence"] = pd.to_numeric(
            sample_enrichment["Divergence"], errors="coerce"
        )

        # --- Main Plotting Logic ---
        Logger.step("Generating jointplot grid...")
        correlations = {"cell_line": cell_line, "modification": modification}
        figure = plt.figure(figsize=(16, 10))
        class_names_grid = np.array(class_names).reshape((2, 3))

        # Create the main 2x3 grid for the entire figure
        outer_grid = GridSpec(2, 3, figure=figure, wspace=0.1, hspace=0.1)

        # --- Store references to the first axis in each row/column ---
        shared_x_axes = [None] * 3  # One for each column
        shared_y_axes = [None] * 2  # One for each row

        for r in range(2):
            for c in range(3):
                class_name = class_names_grid[r, c]
                tqdm.write(f"Plotting: {class_name} (Row {r}, Col {c})")

                # For each cell in the outer grid, create a nested GridSpec
                inner_grid = GridSpecFromSubplotSpec(
                    2,
                    2,
                    subplot_spec=outer_grid[r, c],
                    width_ratios=[5, 1],
                    height_ratios=[1, 5],
                    wspace=0.05,
                    hspace=0.05,
                )

                # --- Create axes with sharing logic ---
                ax_main = figure.add_subplot(
                    inner_grid[1, 0], sharex=shared_x_axes[c], sharey=shared_y_axes[r]
                )
                ax_hist_x = figure.add_subplot(inner_grid[0, 0], sharex=ax_main)
                ax_hist_y = figure.add_subplot(inner_grid[1, 1], sharey=ax_main)

                # --- Store the first axis of each row/col as reference ---
                if r == 0:
                    shared_x_axes[c] = ax_main
                if c == 0:
                    shared_y_axes[r] = ax_main

                # Filter the data for the current class
                query_local = f'Class == "{class_name}"'
                sample_enrichment_local = sample_enrichment.query(query_local)

                # --- Plot the data ---
                sns.histplot(
                    data=sample_enrichment_local,
                    x="Divergence",
                    y="Signal",
                    bins=48,
                    pmax=0.9,
                    cmap="viridis",
                    ax=ax_main,
                )
                sns.histplot(
                    data=sample_enrichment_local, x="Divergence", ax=ax_hist_x, bins=50
                )
                sns.histplot(
                    data=sample_enrichment_local, y="Signal", ax=ax_hist_y, bins=50
                )

                # --- START: MODIFIED TIDY-UP BLOCK ---
                # Tidy up: Remove all ticks, labels, and borders from marginal plots

                # Top histogram (ax_hist_x)
                ax_hist_x.set_ylabel("")
                ax_hist_x.set_xlabel("")
                ax_hist_x.tick_params(
                    axis="both",
                    bottom=False,
                    left=False,
                    labelbottom=False,
                    labelleft=False,
                )
                for spine in ax_hist_x.spines.values():
                    spine.set_visible(False)

                # Right histogram (ax_hist_y)
                ax_hist_y.set_xlabel("")
                ax_hist_y.set_ylabel("")
                ax_hist_y.tick_params(
                    axis="both",
                    bottom=False,
                    left=False,
                    labelbottom=False,
                    labelleft=False,
                )
                for spine in ax_hist_y.spines.values():
                    spine.set_visible(False)
                # --- END: MODIFIED TIDY-UP BLOCK ---

                # --- Tidy up: Main plot ticks/labels (for shared axes) ---
                if c > 0:  # Not in the first column
                    ax_main.tick_params(axis="y", labelleft=False)
                    ax_main.set_ylabel("")
                else:
                    ax_main.set_ylabel(
                        "Signal", fontsize=12
                    )  # Only show Y label on first col

                if r < 1:  # Not in the bottom row
                    ax_main.tick_params(axis="x", labelbottom=False)
                    ax_main.set_xlabel("")
                else:
                    ax_main.set_xlabel(
                        "Divergence", fontsize=12
                    )  # Only show X label on bottom row

                # Calculate correlation
                clean_data = sample_enrichment_local.dropna(
                    subset=["Divergence", "Signal"]
                )
                if len(clean_data) > 1:
                    corr_coef = np.corrcoef(
                        clean_data["Divergence"], clean_data["Signal"]
                    )[0, 1]
                    corr_str = f"r={corr_coef:.3f}"
                else:
                    corr_str = "r=N/A"

                # Set the title for this "jointplot" cell
                ax_hist_x.set_title(f"{class_name}, {corr_str}", fontsize=16, pad=10)
                correlations.update({class_name: corr_coef})

        correlation_df_list.append(pd.Series(correlations))

        # Final figure adjustments
        figure.suptitle(
            f"Divergence vs. Signal for {cell_line} - {modification}",
            fontsize=20,
            y=0.98,
        )
        figure.tight_layout(rect=[0, 0.03, 1, 0.95])

        # Save the final figure
        Logger.step(f"Saving plot to: {save_path_png}")
        # Ensure the directory exists
        os.makedirs(os.path.dirname(save_path_png), exist_ok=True)
        plt.savefig(save_path_png)
        plt.savefig(save_path_svg)
        plt.close(figure)  # IMPORTANT: Close the figure to free up memory
        Logger.success("Plot saved.")
# correlation_df = pd.concat(correlation_df_list)
# correlation_df.to_csv('Signal_vs_divergence_correlation_by_lines_mods_classes.csv')
Logger.info("All processing complete!")

# %% [code]
import os

import matplotlib.pyplot as plt
import numpy as np
import pandas as pd
import seaborn as sns
from matplotlib.gridspec import GridSpec, GridSpecFromSubplotSpec
from tqdm import tqdm  # For progress bars


# --- Helper Class for Colored Logging ---
class Logger:
    """A simple logger for pretty, colored console output."""

    BLUE = "\033[94m"
    GREEN = "\033[92m"
    YELLOW = "\033[93m"
    ENDC = "\033[0m"

    @staticmethod
    def info(message):
        print(f"{Logger.BLUE}INFO: {message}{Logger.ENDC}")

    @staticmethod
    def success(message):
        print(f"{Logger.GREEN}SUCCESS: {message}{Logger.ENDC}")

    @staticmethod
    def warning(message):
        print(f"{Logger.YELLOW}WARNING: {message}{Logger.ENDC}")

    @staticmethod
    def step(message):
        print(f"  -> {message}")


cell_lines = [
    "Caco-2",
    "SJCRH30",
    "BE2C",
    "SJSA1",
    "HAP-1",
    "HL-60",
    "MG63",
    "VCaP",
    "C4-2B",
    "RWPE2",
    "22Rv1",
    "RWPE1",
]
modifications = [
    "CTCF",
    "H3K4me1",
    "H3K9me3",
    "H3K36me3",
    "H3K27me3",
    "H3K4me3",
    "H3K27ac",
]
class_names = ["LINE", "LTR", "SINE", "DNA", "Retroposon", "RC"]

# --- Main Script ---
Logger.info("Starting batch plot generation...")
correlation_df_list = []
# Use tqdm for the outer loop progress bar
for cell_line in tqdm(cell_lines, desc="Processing Cell Lines"):

    # Use a nested tqdm for the inner loop
    for modification in tqdm(
        modifications, desc=f"  Mods for {cell_line}", leave=False
    ):

        filepath = f"./epigenomic_files/{cell_line}.{modification}.chm13v2.0.mapped_on_repeat_masker.bedGraph"
        save_path_png = f"plots/Epigenetic_signal_vs_divergence_by_class/Divergence_vs_Signal_by_classes_{cell_line}_{modification}.png"
        save_path_svg = f"plots/Epigenetic_signal_vs_divergence_by_class/Divergence_vs_Signal_by_classes_{cell_line}_{modification}.svg"
        # --- Data Loading ---
        Logger.step(f"Loading data from: {filepath}")
        try:
            sample_enrichment = pd.read_csv(
                filepath,
                sep="\t",
                header=None,
            )
            Logger.success("File loaded successfully.")
        except FileNotFoundError:
            Logger.warning(f"File not found, skipping: {filepath}")
            continue  # Skip to the next modification
        except Exception as e:
            Logger.warning(
                f"An unexpected error occurred while reading {filepath}: {e}"
            )
            continue

        sample_enrichment.columns = [
            "Chromosome",
            "Start",
            "End",
            "Divergence",
            "Name",
            "Family",
            "Class",
            "Signal",
        ]
        sample_enrichment["Signal"] = pd.to_numeric(
            sample_enrichment["Signal"], errors="coerce"
        )
        sample_enrichment["Divergence"] = pd.to_numeric(
            sample_enrichment["Divergence"], errors="coerce"
        )

        # --- Main Plotting Logic ---
        Logger.step("Generating jointplot grid...")
        # --- Main Plotting Logic ---
        figure = plt.figure(figsize=(16, 10))
        class_names_grid = np.array(class_names).reshape((2, 3))

        # Create the main 2x3 grid for the entire figure
        outer_grid = GridSpec(2, 3, figure=figure, wspace=0.1, hspace=0.1)

        # --- Store references to the first axis in each row/column ---
        shared_x_axes = [None] * 3  # One for each column
        shared_y_axes = [None] * 2  # One for each row

        for r in range(2):
            for c in range(3):
                class_name = class_names_grid[r, c]
                print(f"Plotting: {class_name} (Row {r}, Col {c})")

                # For each cell in the outer grid, create a nested GridSpec
                inner_grid = GridSpecFromSubplotSpec(
                    2,
                    2,
                    subplot_spec=outer_grid[r, c],
                    width_ratios=[5, 1],
                    height_ratios=[1, 5],
                    wspace=0.05,
                    hspace=0.05,
                )

                # --- Create axes with sharing logic ---
                ax_main = figure.add_subplot(
                    inner_grid[1, 0], sharex=shared_x_axes[c], sharey=shared_y_axes[r]
                )
                ax_hist_x = figure.add_subplot(inner_grid[0, 0], sharex=ax_main)
                ax_hist_y = figure.add_subplot(inner_grid[1, 1], sharey=ax_main)

                # --- Store the first axis of each row/col as reference ---
                if r == 0:
                    shared_x_axes[c] = ax_main
                if c == 0:
                    shared_y_axes[r] = ax_main

                # Filter the data for the current class
                query_local = f'Class == "{class_name}"'
                sample_enrichment_local = sample_enrichment.query(query_local)

                # --- Plot the data ---
                sns.histplot(
                    data=sample_enrichment_local,
                    x="Divergence",
                    y="Signal",
                    bins=48,
                    pmax=0.9,
                    cmap="viridis",
                    ax=ax_main,
                )
                sns.histplot(
                    data=sample_enrichment_local, x="Divergence", ax=ax_hist_x, bins=50
                )
                sns.histplot(
                    data=sample_enrichment_local, y="Signal", ax=ax_hist_y, bins=50
                )

                # --- START: MODIFIED TIDY-UP BLOCK ---
                # Tidy up: Remove all ticks, labels, and borders from marginal plots

                # Top histogram (ax_hist_x)
                ax_hist_x.set_ylabel("")
                ax_hist_x.set_xlabel("")
                ax_hist_x.tick_params(
                    axis="both",
                    bottom=False,
                    left=False,
                    labelbottom=False,
                    labelleft=False,
                )
                for spine in ax_hist_x.spines.values():
                    spine.set_visible(False)

                # Right histogram (ax_hist_y)
                ax_hist_y.set_xlabel("")
                ax_hist_y.set_ylabel("")
                ax_hist_y.tick_params(
                    axis="both",
                    bottom=False,
                    left=False,
                    labelbottom=False,
                    labelleft=False,
                )
                for spine in ax_hist_y.spines.values():
                    spine.set_visible(False)
                # --- END: MODIFIED TIDY-UP BLOCK ---

                # --- Tidy up: Main plot ticks/labels (for shared axes) ---
                if c > 0:  # Not in the first column
                    ax_main.tick_params(axis="y", labelleft=False)
                    ax_main.set_ylabel("")
                else:
                    ax_main.set_ylabel(
                        "Signal", fontsize=12
                    )  # Only show Y label on first col

                if r < 1:  # Not in the bottom row
                    ax_main.tick_params(axis="x", labelbottom=False)
                    ax_main.set_xlabel("")
                else:
                    ax_main.set_xlabel(
                        "Divergence", fontsize=12
                    )  # Only show X label on bottom row

                # Calculate correlation
                clean_data = sample_enrichment_local.dropna(
                    subset=["Divergence", "Signal"]
                )
                if len(clean_data) > 1:
                    corr_coef = np.corrcoef(
                        clean_data["Divergence"], clean_data["Signal"]
                    )[0, 1]
                    corr_str = f"r={corr_coef:.3f}"
                else:
                    corr_str = "r=N/A"

                # Set the title for this "jointplot" cell
                ax_hist_x.set_title(f"{class_name}, {corr_str}", fontsize=16, pad=10)

        # Adjust layout to prevent labels from overlapping
        plt.suptitle(
            "Divergence vs. Signal by Repeat Class (Shared Axes)", fontsize=20, y=0.98
        )
        figure.tight_layout(
            rect=[0, 0.03, 1, 0.95]
        )  # Adjust rect to make space for suptitle

        # Save the final figure
        Logger.step(f"Saving plot to: {save_path}")
        # Ensure the directory exists
        os.makedirs(os.path.dirname(save_path_png), exist_ok=True)
        plt.savefig(save_path_png)
        plt.savefig(save_path_svg)
        plt.close(figure)  # IMPORTANT: Close the figure to free up memory
        Logger.success("Plot saved.")
correlation_df = pd.concat(correlation_df_list, axis=1)
correlation_df.to_csv("Signal_vs_divergence_correlation_by_lines_mods_classes.csv")
Logger.info("All processing complete!")

# %% [code]
pd.concat([pd.Series({"a": 1, "b": 1}), pd.Series({"a": 1, "b": 1})], axis=1)

# %% [code]
list(sample_enrichment.Signal[0:100000])

# %% [markdown]
### Set up cell lines and modifications palettes, save as legends for figures

# %% [code]
import matplotlib.colors as mcolors

cmap = matplotlib.cm.get_cmap("Pastel1")
# Access the .colors attribute (a list of RGBA tuples)
rgba_colors = cmap.colors

# Convert each RGBA tuple to a hex string
hex_colors = [mcolors.to_hex(c) for c in rgba_colors]

# Print the list of hex color strings
print(hex_colors)

# %% [code]
cmap

# %% [code]
cmap = matplotlib.cm.get_cmap("Pastel2")
# Access the .colors attribute (a list of RGBA tuples)
rgba_colors = cmap.colors

# Convert each RGBA tuple to a hex string
hex_colors = [mcolors.to_hex(c) for c in rgba_colors]

# Print the list of hex color strings
print(hex_colors)

# %% [code]
cmap

# %% [code]
cell_line_p = {
    "Caco-2": "saddlebrown",
    "SJCRH30": "darkgreen",
    "BE2C": "grey",
    "SJSA1": "teal",
    "HAP-1": "royalblue",
    "HL-60": "cyan",
    "MG63": "lime",
    "VCaP": "gold",
    "C4-2B": "orangered",
    "RWPE2": "orange",
    "22Rv1": "lightcoral",
    "RWPE1": "deeppink",
}
modification_p = {
    "CTCF": "#e5d8bd",
    "H3K4me1": "#fdcdac",
    "H3K9me3": "#b3cde3",
    "H3K36me3": "#e6f5c9",
    "H3K27me3": "#decbe4",
    "H3K4me3": "#b3e2cd",
    "H3K27ac": "#fff2ae",
}

modification_p_richer = {
    "CTCF": "#d3c09b",
    "H3K4me1": "#fabd88",
    "H3K9me3": "#9ab7d3",
    "H3K36me3": "#d7f0a8",
    "H3K27me3": "#c7aed6",
    "H3K4me3": "#9ad1b5",
    "H3K27ac": "#ffe98c",
}

# %% [code]
import os

import matplotlib.patches as mpatches
import matplotlib.pyplot as plt

# 1. Create a list of 'Patch' objects for the legend handles
# A Patch is just a colored rectangle
patches = [
    mpatches.Patch(color=color, label=label) for label, color in cell_line_p.items()
]

# 2. Create a figure and a single axis
# We can make the figure small, as we will crop it tightly
fig, ax = plt.subplots(figsize=(4, 2))

# 3. Create the legend on the axis
# We place it in the 'center' of the invisible axis
legend = ax.legend(
    handles=patches,
    loc="center",
    frameon=False,  # Remove the legend box frame
    fontsize=12,
    ncol=1,  # Arrange in a single column
)

# 4. Turn off the axis (spines, ticks, etc.)
ax.axis("off")

# 5. Define the save path
save_path = "./plots/cell_line_legend.svg"

# Ensure the 'plots' directory exists (if you want to save it there)
# os.makedirs(os.path.dirname(save_path), exist_ok=True)

# 6. Save the figure as an SVG
# bbox_inches='tight' is the key to cropping the output
# to the legend's bounding box.
plt.savefig(
    save_path,
    format="svg",
    bbox_inches="tight",
    pad_inches=0.1,  # Add a small amount of padding
    transparent=True,  # Make the background transparent
)

plt.close(fig)  # Close the figure to free memory

print(f"Legend saved successfully to {save_path}")

# %% [code]


# %% [code]


# %% [markdown]
## Analyze correlation coefficients of signal vs divergence by TE class, line and modification

# %% [code]
corr_coefs = pd.read_csv("Signal_vs_divergence_correlation_by_lines_mods_classes.csv")
class_names = ["LINE", "LTR", "SINE", "DNA", "Retroposon", "RC"]

correlation_df_list = []
for i in range(0, 39):
    correlation_df_list.append(corr_coefs[8 * i : 8 * i + 8].set_index("Unnamed: 0"))

correlation_df = pd.concat(correlation_df_list, axis=1).T
correlation_df.index = correlation_df.cell_line + "_" + correlation_df.modification
correlation_df[class_names] = correlation_df[class_names].apply(
    pd.to_numeric, errors="coerce"
)
correlation_df.to_csv('./supplementary/Supplementary table 3.csv')
correlation_df  # [correlation_df.modification == 'H3K9me3']

# %% [code]
correlation_df.reindex(
    ["RWPE1_CTCF", "SJCRH30_H3K27me3", "SJCRH30_H3K4me1", "BE2C_H3K4me1"]
)

# %% [code]
correlation_df.drop(["C4-2B_CTCF", "RWPE1_H3K27ac"]).min()

# %% [code]
correlation_df.drop(["C4-2B_CTCF", "RWPE1_H3K27ac"]).max()

# %% [code]
correlation_df[class_names].min()

# %% [code]
# --- 1. Prepare Row Colors ---
# Map the 'cell_line' and 'modification' columns to their respective colors
mapped_cell_colors = correlation_df["cell_line"].map(cell_line_p)
mapped_mod_colors = correlation_df["modification"].map(modification_p_richer)

# Create a DataFrame for the row_colors argument
# The column names (e.g., 'Cell Line') will appear as labels.
row_colors_df = pd.DataFrame(
    {"Cell Line": mapped_cell_colors, "Modification": mapped_mod_colors}
)
# Important: The index must align with the data being plotted
row_colors_df.index = correlation_df.index


# --- 2. Plot the Clustermap ---
data_to_plot = correlation_df[class_names].T


g = sns.clustermap(
    data_to_plot,
    cmap="coolwarm",
    # Goal 1: Customize figsize
    figsize=(12, 8),
    # Goal 2 & 3: Add row colors using the mapped palettes
    # We use row_colors, as 'cell_line' and 'modification' describe the rows
    col_colors=row_colors_df[["Modification", "Cell Line"]],
    # Other aesthetic improvements
    # vmin=-1, vmax=1,          # Set colorbar limits for correlation
    linewidths=0.5,
    cbar_kws={"label": "Pearson Correlation"},
    # metric='correlation'      # Use correlation for clustering distance
)

# --- 3. Save the Figure ---
save_path = "./plots/signal_to_divergence_corr_clustermap.svg"


# Goal 4: Save as SVG
g.savefig(save_path, format="svg")


plt.show()

# %% [code]
class_names = ["LINE", "LTR", "SINE", "DNA", "Retroposon", "RC"]
sns.clustermap(
    correlation_df[class_names],
    cmap="coolwarm",
)

# %% [code]
corr_coefs[0:8].set_index("Unnamed: 0")

# %% [markdown]
### Test for significant differences between correlation coefficients with divergence of different families 

# %% [markdown]
That's an excellent question. The "pattern" you're looking for can be defined in a few different statistical ways.The most important first step, before any test, is to transform your correlation coefficients ($r$).The Prerequisite: Fisher $z$-TransformationYou should not perform statistical tests directly on the raw Pearson $r$ values. The distribution of $r$ is not normal and is bounded between -1 and 1 (it "bunches up" as it gets close to the ends).You must first convert every $r$ value in your set to a $z$-score using the Fisher $z$-transformation. This transformation "stretches" the values so they are approximately normally distributed, which is an assumption for many tests.The formula is:$$z = 0.5 \times \ln\left(\frac{1+r}{1-r}\right)$$In most software (like Python/Numpy or R), this is simply z = np.arctanh(r).From now on, all tests are performed on your set of $z$-scores, not the $r$-values.Here are the most common "patterns" you can test for, from simplest to most complex.1. Does the distribution follow a specific shape (e.g., normal)?This tests if the set itself has a recognizable structure.Your Question: "Is my set of $z$-scores normally distributed?" (This would imply the correlations are clustered around a single mean value in a standard way).How to Test:Visually: Plot a histogram or a Q-Q plot of your $z$-scores. This is often the most insightful step. A Q-Q plot will show you if the data follows a straight line (normal) or where it deviates (e.g., "heavy tails").Formally: Use a Shapiro-Wilk Test. The null hypothesis is that the data is normally distributed. A low p-value (e.g., $< 0.05$) suggests it is not normal.Interpretation: If it's not normal, the visual plot tells you the pattern. Is it bimodal (two peaks)? Is it skewed? This is a pattern in itself.2. Does the set of correlations relate to another variable?This is often the most powerful and common question. You have a list of correlations, $r_1, r_2, \dots, r_n$. Do you have another property for each of those $n$ vectors?Your Question: "Do the vectors that correlate highly with my target vector also share some other property?"Example:Your vectors are Gene 1, Gene 2, ... Gene 1000.Your target vector is Disease_Severity.Your set of $r$-values is [Corr(Gene1, Disease), Corr(Gene2, Disease), ...].Your other variable could be the "Average Expression Level" for each gene: [Avg(Gene1), Avg(Gene2), ...].The test answers: "Do genes that strongly correlate with the disease also have high average expression?"How to Test:Create your list of $z$-scores: $Z = [z_1, z_2, \dots, z_n]$.Create your list of the other property: $P = [p_1, p_2, \dots, p_n]$.Calculate a Spearman Rank Correlation between $Z$ and $P$.Interpretation: A significant Spearman correlation (e.g., $p < 0.05$) shows a monotonic pattern: as one value increases, the other tends to increase (or decrease). This is a very strong and common way to find a "pattern."3. Is the average correlation of the set different from zero?This tests if your "many vectors" have any relationship at all (on average) with the target vector.Your Question: "Is the mean of my correlation set significantly positive or negative?"How to Test:Get your set of $z$-scores.Run a one-sample t-test with a null hypothesis (mu) of 0.Interpretation: A low p-value means the average correlation is not zero. This confirms a general, set-wide pattern of a positive or negative relationship.4. Are the correlations different between groups?If your "many vectors" can be divided into pre-defined categories, you can test if the correlation pattern is different between them.Your Question: "Is the average correlation to the target vector different for Group A vs. Group B?"Example: "Is the correlation of 'kinase' genes with the target different from the correlation of 'transcription factor' genes with the target?"How to Test:Get your $z$-scores and label them by group (e.g., 'Group A', 'Group B').For 2 groups: Run a Welch's t-test (or a Mann-Whitney U test if the $z$-scores are still not normal) between the two groups.For 3+ groups: Run an ANOVA (or a Kruskal-Wallis test) across all groups.Interpretation: A low p-value shows that the "pattern" of correlation is not the same across your defined categories.

# %% [code]
correlation_df

# %% [code]
corr_coefs = pd.read_csv("Signal_vs_divergence_correlation_by_lines_mods_classes.csv")
class_names = ["LINE", "LTR", "SINE", "DNA", "Retroposon", "RC"]

correlation_df_list = []
for i in range(0, 39):
    correlation_df_list.append(corr_coefs[8 * i : 8 * i + 8].set_index("Unnamed: 0"))

correlation_df = pd.concat(correlation_df_list, axis=1).T
correlation_df.index = correlation_df.cell_line + "_" + correlation_df.modification
correlation_df[class_names] = correlation_df[class_names].apply(
    pd.to_numeric, errors="coerce"
)

corrs_for_test = correlation_df.melt(
    id_vars=["cell_line", "modification"], value_vars=class_names
)
corrs_for_test.columns = ["cell_line", "modification", "TE_class", "correlation"]
corrs_for_test["transformed_corr"] = np.arctanh(corrs_for_test.correlation)
corrs_for_test

# %% [code]
sns.histplot(corrs_for_test.correlation)

# %% [code]
sns.histplot(corrs_for_test.transformed_corr)  # Fisher z-transformation

# %% [code]
stats.shapiro(corrs_for_test.transformed_corr)

# %% [code]
stats.kurtosis(corrs_for_test.transformed_corr)

# %% [code]
stats.kurtosistest(corrs_for_test.transformed_corr)

# %% [code]
figure, ax = plt.subplots(
    figsize=(7, 4),
)
sns.boxplot(
    data=corrs_for_test,
    x="TE_class",
    y="transformed_corr",
    ax=ax,
    # hue='TE_class'
)

# %% [code]
figure, ax = plt.subplots(
    figsize=(9, 4),
)
sns.boxplot(
    data=corrs_for_test,
    x="modification",
    y="transformed_corr",
    ax=ax,
    # hue='TE_class'
)

# %% [code]
figure, ax = plt.subplots(
    figsize=(9, 4),
)
sns.boxplot(
    data=corrs_for_test,
    x="cell_line",
    y="transformed_corr",
    ax=ax,
    # hue='TE_class'
)

# %% [code]
np.arctanh

# %% [markdown]
#### Test for equality of variances

# %% [code]
corrs_for_test[]

# %% [code]
class_names

# %% [code]
from scipy.stats import levene
??levene

# %% [code]
from scipy.stats import levene

LINE = corrs_for_test["transformed_corr"][corrs_for_test["TE_class"] == "LINE"]
LTR = corrs_for_test["transformed_corr"][corrs_for_test["TE_class"] == "LTR"]
SINE = corrs_for_test["transformed_corr"][corrs_for_test["TE_class"] == "SINE"]
DNA = corrs_for_test["transformed_corr"][corrs_for_test["TE_class"] == "DNA"]
Retroposon = corrs_for_test["transformed_corr"][
    corrs_for_test["TE_class"] == "Retroposon"
]
RC = corrs_for_test["transformed_corr"][corrs_for_test["TE_class"] == "RC"]
non_sva = corrs_for_test["transformed_corr"][corrs_for_test["TE_class"] != "Retroposon"]


levene(LINE, LTR, SINE, DNA, Retroposon, RC)

# %% [code]
levene(Retroposon, non_sva)

# %% [code]
from statsmodels.stats.oneway import anova_oneway

# Perform the Brown-Forsythe test
anova_oneway(
    data=corrs_for_test["transformed_corr"],
    groups=corrs_for_test["TE_class"],
    use_var="equal",
)

# %% [code]
??anova_oneway

# %% [code]


# %% [markdown]
### Attempts of plot of signal vs Divergence correlation

# %% [code]
from scipy import stats

# %% [code]
class_names = ["LINE", "LTR", "SINE", "DNA", "Retroposon", "RC"]

figure, axs = plt.subplots(
    nrows=2,
    ncols=3,
    sharex=True,
    sharey=False,
    squeeze=True,
    # width_ratios=None,
    # height_ratios=[1, 1, 1, 1, 1, 1, 0.3, 0.3],
    figsize=(14, 8),
)

# Use zip to iterate over class names and the flattened axes array
# axs.flatten() converts the 2x3 array into a 1D array of 6 axes
for class_name, ax in zip(class_names, axs.flatten()):
    print(class_name)
    # Filter the data
    query_local = f'Class == "{class_name}"'
    sample_enrichment_local = sample_enrichment.query(query_local).dropna()
    corr_coef = np.corrcoef(
        sample_enrichment_local["Divergence"], sample_enrichment_local["Signal"]
    )[0, 1]
    # Use sns.histplot (axes-level) instead of sns.displot (figure-level)

    sns.kdeplot(  #
        data=sample_enrichment_local,
        x="Divergence",
        y="Signal",
        # alpha=0.01,
        ax=ax,
        fill=True,
        thresh=0,
        levels=100,
        cmap="mako",
    )

    # Use ax.set_title() to set the title for the subplot
    ax.set_title(class_name)

plt.show()
# Save the final figure
# plt.savefig("plots/Divergence_score_dist_by_class.svg")

# %% [code]
class_names = ["LINE", "LTR", "SINE", "DNA", "Retroposon", "RC"]

figure, axs = plt.subplots(
    nrows=2,
    ncols=3,
    sharex=True,
    sharey=False,
    squeeze=True,
    # width_ratios=None,
    # height_ratios=[1, 1, 1, 1, 1, 1, 0.3, 0.3],
    figsize=(14, 8),
)

# Use zip to iterate over class names and the flattened axes array
# axs.flatten() converts the 2x3 array into a 1D array of 6 axes
for class_name, ax in zip(class_names, axs.flatten()):
    print(class_name)
    # Filter the data
    query_local = f'Class == "{class_name}"'
    sample_enrichment_local = sample_enrichment.query(query_local).dropna()

    # Use sns.histplot (axes-level) instead of sns.displot (figure-level)

    sns.scatterplot(  # kdeplot
        data=sample_enrichment_local,
        x="Divergence",
        y="Signal",
        alpha=0.01,
        ax=ax,
        # fill=True, thresh=0, levels=100, cmap="mako",
    )

    # Use ax.set_title() to set the title for the subplot
    ax.set_title(class_name)

plt.show()
# Save the final figure
# plt.savefig("plots/Divergence_score_dist_by_class.svg")

# %% [markdown]
## Record correlation coefficients of inividual families of TEs

# %% [code]
repeat_masker = pd.read_csv("T2T_repeat_masker_processed.csv")

# %% [code]
repeat_masker

# %% [code]
repeat_masker.family_name.value_counts().to_csv("repeat_masker_family_name_counts.csv")

# %% [code]
repeat_masker.individual_name.value_counts().to_csv(
    "repeat_masker_individual_name_counts.csv"
)

# %% [code]
repeat_masker_family_name_counts = pd.read_csv("repeat_masker_family_name_counts.csv")[
    "Unnamed: 0"
]
repeat_masker_family_name_counts

# %% [code]
import os

import numpy as np
import pandas as pd
from scipy import stats
from tqdm import tqdm  # For progress bars


def pearsonr_ci(x, y, alpha=0.05):
    """
    Calculates Pearson r, p-value, and 95% CI using Fisher z-transformation.
    This works on all versions of SciPy.
    """
    # Calculate r and p-value
    r, p = stats.pearsonr(x, y)
    n = len(x)

    # Fisher z-transformation
    z = np.arctanh(r)

    # Standard error of z
    se_z = 1.0 / np.sqrt(n - 3)

    # Get the z-score for the desired confidence level
    z_crit = stats.norm.ppf(1 - alpha / 2)  # This is 1.96 for 95% CI

    # Calculate CI for z
    z_low = z - z_crit * se_z
    z_high = z + z_crit * se_z

    # Back-transform z CI to r CI
    r_low = np.tanh(z_low)
    r_high = np.tanh(z_high)

    return r, p, r_low, r_high


# --- Helper Class for Colored Logging ---
class Logger:
    """A simple logger for pretty, colored console output."""

    BLUE = "\033[94m"
    GREEN = "\033[92m"
    YELLOW = "\033[93m"
    ENDC = "\033[0m"

    @staticmethod
    def info(message):
        print(f"{Logger.BLUE}INFO: {message}{Logger.ENDC}")

    @staticmethod
    def success(message):
        print(f"{Logger.GREEN}SUCCESS: {message}{Logger.ENDC}")

    @staticmethod
    def warning(message):
        print(f"{Logger.YELLOW}WARNING: {message}{Logger.ENDC}")

    @staticmethod
    def step(message):
        print(f"  -> {message}")


cell_lines = [
    "Caco-2",
    "SJCRH30",
    "BE2C",
    "SJSA1",
    "HAP-1",
    "HL-60",
    "MG63",
    "VCaP",
    "C4-2B",
    "RWPE2",
    "22Rv1",
    "RWPE1",
]
modifications = [
    "CTCF",
    "H3K27ac",
    "H3K4me1",
    "H3K9me3",
    "H3K36me3",
    "H3K27me3",
    "H3K4me3",
]

family_names = pd.read_csv("repeat_masker_family_name_counts.csv")["Unnamed: 0"]
individual_names = pd.read_csv("repeat_masker_individual_name_counts.csv")["Unnamed: 0"]

# Name	Family	Class
# --- Main Script ---
Logger.info("Starting batch plot generation...")
families_stats_list = []

individuals_stats_list = []
# Use tqdm for the outer loop progress bar
for cell_line in tqdm(cell_lines, desc="Processing Cell Lines"):

    # Use a nested tqdm for the inner loop
    for modification in tqdm(
        modifications, desc=f"  Mods for {cell_line}", leave=False
    ):

        filepath = f"./epigenomic_files/{cell_line}.{modification}.chm13v2.0.mapped_on_repeat_masker.bedGraph"
        # --- Data Loading ---
        Logger.step(f"Loading data from: {filepath}")
        try:
            sample_enrichment = pd.read_csv(
                filepath,
                sep="\t",
                header=None,
            )
            Logger.success("File loaded successfully.")
        except FileNotFoundError:
            Logger.warning(f"File not found, skipping: {filepath}")
            continue  # Skip to the next modification
        except Exception as e:
            Logger.warning(
                f"An unexpected error occurred while reading {filepath}: {e}"
            )
            continue

        sample_enrichment.columns = [
            "Chromosome",
            "Start",
            "End",
            "Divergence",
            "Name",
            "Family",
            "Class",
            "Signal",
        ]
        sample_enrichment["Signal"] = pd.to_numeric(
            sample_enrichment["Signal"], errors="coerce"
        )
        sample_enrichment["Divergence"] = pd.to_numeric(
            sample_enrichment["Divergence"], errors="coerce"
        )
        for family_name in tqdm(family_names, desc="Processing Family Names"):

            # Filter the data for the current family
            query_local = f'Family == "{family_name}"'
            sample_enrichment_local = sample_enrichment.query(query_local)

            # Calculate correlation
            clean_data = sample_enrichment_local.dropna(subset=["Divergence", "Signal"])
            if len(clean_data) > 3:
                corr_coef, p_val, ci_low, ci_high = pearsonr_ci(
                    clean_data["Divergence"], clean_data["Signal"]
                )
            else:
                corr_coef, p_val, ci_low, ci_high = None, None, None, None

            families_stats_list.append(
                {
                    "cell_line": cell_line,
                    "modification": modification,
                    "family_name": family_name,
                    "r": corr_coef,
                    "p_value": p_val,
                    "ci_low": ci_low,
                    "ci_high": ci_high,
                }
            )

        for individual_name in tqdm(
            individual_names, desc="Processing Individual Names"
        ):

            # Filter the data for the current name
            query_local = f'Name == "{individual_name}"'
            sample_enrichment_local = sample_enrichment.query(query_local)

            # Calculate correlation
            clean_data = sample_enrichment_local.dropna(subset=["Divergence", "Signal"])
            if len(clean_data) > 3:
                corr_coef, p_val, ci_low, ci_high = pearsonr_ci(
                    clean_data["Divergence"], clean_data["Signal"]
                )
            else:
                corr_coef, p_val, ci_low, ci_high = None, None, None, None

            individuals_stats_list.append(
                {
                    "cell_line": cell_line,
                    "modification": modification,
                    "individual_name": individual_name,
                    "r": corr_coef,
                    "p_value": p_val,
                    "ci_low": ci_low,
                    "ci_high": ci_high,
                }
            )


correlation_df_families = pd.DataFrame(families_stats_list)
correlation_df_individuals = pd.DataFrame(individuals_stats_list)

correlation_df_families.to_csv(
    "Signal_vs_divergence_correlation_by_lines_mods_families_detailed.csv", index=True
)


correlation_df_individuals.to_csv(
    "Signal_vs_divergence_correlation_by_lines_mods_individuals_detailed.csv",
    index=True,
)
Logger.info("All processing complete!")

# %% [markdown]
### Record correlation coefficients with random control

# %% [code]
import os

import numpy as np
import pandas as pd
from scipy import stats
from tqdm import tqdm  # For progress bars

import warnings
warnings.filterwarnings('ignore')
def pearsonr_ci(x, y, alpha=0.05):
    """
    Calculates Pearson r, p-value, and 95% CI using Fisher z-transformation.
    This works on all versions of SciPy.
    """
    # Calculate r and p-value
    r, p = stats.pearsonr(x, y)
    n = len(x)

    # Fisher z-transformation
    z = np.arctanh(r)

    # Standard error of z
    se_z = 1.0 / np.sqrt(n - 3)

    # Get the z-score for the desired confidence level
    z_crit = stats.norm.ppf(1 - alpha / 2)  # This is 1.96 for 95% CI

    # Calculate CI for z
    z_low = z - z_crit * se_z
    z_high = z + z_crit * se_z

    # Back-transform z CI to r CI
    r_low = np.tanh(z_low)
    r_high = np.tanh(z_high)

    return r, p, r_low, r_high


# --- Helper Class for Colored Logging ---
class Logger:
    """A simple logger for pretty, colored console output."""

    BLUE = "\033[94m"
    GREEN = "\033[92m"
    YELLOW = "\033[93m"
    ENDC = "\033[0m"

    @staticmethod
    def info(message):
        print(f"{Logger.BLUE}INFO: {message}{Logger.ENDC}")

    @staticmethod
    def success(message):
        print(f"{Logger.GREEN}SUCCESS: {message}{Logger.ENDC}")

    @staticmethod
    def warning(message):
        print(f"{Logger.YELLOW}WARNING: {message}{Logger.ENDC}")

    @staticmethod
    def step(message):
        print(f"  -> {message}")


cell_lines = [
    "Caco-2",
    "SJCRH30",
    "BE2C",
    "SJSA1",
    "HAP-1",
    "HL-60",
    "MG63",
    "VCaP",
    "C4-2B",
    "RWPE2",
    "22Rv1",
    "RWPE1",
]
modifications = [
    "CTCF",
    "H3K27ac",
    "H3K4me1",
    "H3K9me3",
    "H3K36me3",
    "H3K27me3",
    "H3K4me3",
]

family_names = pd.read_csv("repeat_masker_family_name_counts.csv")["Unnamed: 0"]
individual_names = pd.read_csv("repeat_masker_individual_name_counts.csv")["Unnamed: 0"]

# Name	Family	Class
# --- Main Script ---
Logger.info("Starting batch plot generation...")
families_stats_list = []

individuals_stats_list = []
# Use tqdm for the outer loop progress bar
for cell_line in tqdm(cell_lines, desc="Processing Cell Lines"):

    # Use a nested tqdm for the inner loop
    for modification in tqdm(
        modifications, desc=f"  Mods for {cell_line}", leave=False
    ):

        filepath = f"./epigenomic_files/{cell_line}.{modification}.chm13v2.0.mapped_on_repeat_masker.bedGraph"
        # --- Data Loading ---
        Logger.step(f"Loading data from: {filepath}")
        try:
            sample_enrichment = pd.read_csv(
                filepath,
                sep="\t",
                header=None,
            )
            Logger.success("File loaded successfully.")
        except FileNotFoundError:
            Logger.warning(f"File not found, skipping: {filepath}")
            continue  # Skip to the next modification
        except Exception as e:
            Logger.warning(
                f"An unexpected error occurred while reading {filepath}: {e}"
            )
            continue

        sample_enrichment.columns = [
            "Chromosome",
            "Start",
            "End",
            "Divergence",
            "Name",
            "Family",
            "Class",
            "Signal",
        ]
        sample_enrichment["Signal"] = pd.to_numeric(
            sample_enrichment["Signal"], errors="coerce"
        )
        sample_enrichment["Divergence"] = pd.to_numeric(
            sample_enrichment["Divergence"], errors="coerce"
        )
        for family_name in tqdm(family_names, desc="Processing Family Names"):

            # Filter the data for the current family
            query_local = f'Family == "{family_name}"'
            sample_enrichment_local = sample_enrichment.query(query_local)

            # Calculate correlation
            clean_data = sample_enrichment_local.dropna(subset=["Divergence", "Signal"])
            clean_data['Signal'] = clean_data['Signal'].sample(frac=1, random_state=1).set_axis(clean_data.index)
            if len(clean_data) > 3:
                corr_coef, p_val, ci_low, ci_high = pearsonr_ci(
                    clean_data["Divergence"], clean_data["Signal"]
                )
            else:
                corr_coef, p_val, ci_low, ci_high = None, None, None, None

            families_stats_list.append(
                {
                    "cell_line": cell_line,
                    "modification": modification,
                    "family_name": family_name,
                    "r": corr_coef,
                    "p_value": p_val,
                    "ci_low": ci_low,
                    "ci_high": ci_high,
                }
            )

        for individual_name in tqdm(
            individual_names, desc="Processing Individual Names"
        ):

            # Filter the data for the current name
            query_local = f'Name == "{individual_name}"'
            sample_enrichment_local = sample_enrichment.query(query_local)

            # Calculate correlation
            clean_data = sample_enrichment_local.dropna(subset=["Divergence", "Signal"])
            clean_data['Signal'] = clean_data['Signal'].sample(frac=1, random_state=1).set_axis(clean_data.index)
            if len(clean_data) > 3:
                corr_coef, p_val, ci_low, ci_high = pearsonr_ci(
                    clean_data["Divergence"], clean_data["Signal"]
                )
            else:
                corr_coef, p_val, ci_low, ci_high = None, None, None, None

            individuals_stats_list.append(
                {
                    "cell_line": cell_line,
                    "modification": modification,
                    "individual_name": individual_name,
                    "r": corr_coef,
                    "p_value": p_val,
                    "ci_low": ci_low,
                    "ci_high": ci_high,
                }
            )


correlation_df_families = pd.DataFrame(families_stats_list)
correlation_df_individuals = pd.DataFrame(individuals_stats_list)

correlation_df_families.to_csv(
    "Signal_vs_divergence_correlation_by_lines_mods_families_detailed_random.csv", index=True
)


correlation_df_individuals.to_csv(
    "Signal_vs_divergence_correlation_by_lines_mods_individuals_detailed_random.csv",
    index=True,
)
Logger.info("All processing complete!")

# %% [code]
clean_data = sample_enrichment_local.dropna(subset=["Divergence", "Signal"])
clean_data['Signal'] = clean_data['Signal'].sample(frac=1, random_state=1).set_axis(clean_data.index)
clean_data

# %% [code]
clean_data = sample_enrichment_local.dropna(subset=["Divergence", "Signal"])

clean_data

# %% [code]
clean_data['Signal'].sample(frac=1, random_state=1).reset_index(drop=True)

# %% [markdown]
## Record averages and SDs of epigenetic signal by TE classes, families and individuals 

# %% [code]
import os

import numpy as np
import pandas as pd
from tqdm import tqdm  # For progress bars


# --- Helper Class for Colored Logging ---
class Logger:
    """A simple logger for pretty, colored console output."""

    BLUE = "\033[94m"
    GREEN = "\033[92m"
    YELLOW = "\033[93m"
    ENDC = "\033[0m"

    @staticmethod
    def info(message):
        print(f"{Logger.BLUE}INFO: {message}{Logger.ENDC}")

    @staticmethod
    def success(message):
        print(f"{Logger.GREEN}SUCCESS: {message}{Logger.ENDC}")

    @staticmethod
    def warning(message):
        print(f"{Logger.YELLOW}WARNING: {message}{Logger.ENDC}")

    @staticmethod
    def step(message):
        print(f"  -> {message}")


cell_lines = [
    "Caco-2",
    "SJCRH30",
    "BE2C",
    "SJSA1",
    "HAP-1",
    "HL-60",
    "MG63",
    "VCaP",
    "C4-2B",
    "RWPE2",
    "22Rv1",
    "RWPE1",
]
modifications = [
    "CTCF",
    "H3K27ac",
    "H3K4me1",
    "H3K9me3",
    "H3K36me3",
    "H3K27me3",
    "H3K4me3",
]

family_names = pd.read_csv("repeat_masker_family_name_counts.csv")["Unnamed: 0"]
individual_names = pd.read_csv("repeat_masker_individual_name_counts.csv")["Unnamed: 0"]


# Name	Family	Class
# --- Main Script ---
Logger.info("Starting batch statistics calculation...")
means_df_list_classes = []
means_df_list_families = []
means_df_list_individuals = []

sds_df_list_classes = []
sds_df_list_families = []
sds_df_list_individuals = []

# Use tqdm for the outer loop progress bar
for cell_line in tqdm(cell_lines, desc="Processing Cell Lines"):

    # Use a nested tqdm for the inner loop
    for modification in tqdm(
        modifications, desc=f"  Mods for {cell_line}", leave=False
    ):

        filepath = f"./epigenomic_files/{cell_line}.{modification}.chm13v2.0.mapped_on_repeat_masker.bedGraph"
        # --- Data Loading ---
        Logger.step(f"Loading data from: {filepath}")
        try:
            sample_enrichment = pd.read_csv(
                filepath,
                sep="\t",
                header=None,
            )
            Logger.success("File loaded successfully.")
        except FileNotFoundError:
            Logger.warning(f"File not found, skipping: {filepath}")
            continue  # Skip to the next modification
        except Exception as e:
            Logger.warning(
                f"An unexpected error occurred while reading {filepath}: {e}"
            )
            continue

        sample_enrichment.columns = [
            "Chromosome",
            "Start",
            "End",
            "Divergence",
            "Name",
            "Family",
            "Class",
            "Signal",
        ]
        sample_enrichment["Signal"] = pd.to_numeric(
            sample_enrichment["Signal"], errors="coerce"
        )
        sample_enrichment["Divergence"] = pd.to_numeric(
            sample_enrichment["Divergence"], errors="coerce"
        )

        # --- Means ---
        class_means = sample_enrichment.groupby("Class")["Signal"].mean().to_dict()
        class_means.update({"cell_line": cell_line, "modification": modification})
        means_df_list_classes.append(class_means)

        family_means = sample_enrichment.groupby("Family")["Signal"].mean().to_dict()
        family_means.update({"cell_line": cell_line, "modification": modification})
        means_df_list_families.append(family_means)

        individual_means = sample_enrichment.groupby("Name")["Signal"].mean().to_dict()
        individual_means.update({"cell_line": cell_line, "modification": modification})
        means_df_list_individuals.append(individual_means)

        # --- Standard Deviations ---
        class_sds = sample_enrichment.groupby("Class")["Signal"].std().to_dict()
        class_sds.update({"cell_line": cell_line, "modification": modification})
        sds_df_list_classes.append(class_sds)

        family_sds = sample_enrichment.groupby("Family")["Signal"].std().to_dict()
        family_sds.update({"cell_line": cell_line, "modification": modification})
        sds_df_list_families.append(family_sds)

        individual_sds = sample_enrichment.groupby("Name")["Signal"].std().to_dict()
        individual_sds.update({"cell_line": cell_line, "modification": modification})
        sds_df_list_individuals.append(individual_sds)

# --- CORRECTED DataFrame construction ---
# Use pd.DataFrame() to build from a list of dicts (rows)
# Use index=False so you don't save the 0, 1, 2... row numbers
Logger.info("Assembling and saving DataFrames...")

pd.DataFrame(means_df_list_classes).to_csv(
    "Average_signal_by_lines_mods_TE_classes.csv", index=False
)

pd.DataFrame(means_df_list_families).to_csv(
    "Average_signal_by_lines_mods_TE_families.csv", index=False
)

pd.DataFrame(means_df_list_individuals).to_csv(
    "Average_signal_by_lines_mods_TE_individuals.csv", index=False
)

pd.DataFrame(sds_df_list_classes).to_csv(
    "SD_signal_by_lines_mods_TE_classes.csv", index=False
)

pd.DataFrame(sds_df_list_families).to_csv(
    "SD_signal_by_lines_mods_TE_families.csv", index=False
)

pd.DataFrame(sds_df_list_individuals).to_csv(
    "SD_signal_by_lines_mods_TE_individuals.csv", index=False
)

Logger.success("All processing complete!")

# %% [markdown]
## Analyze correlation coefficients of inividual families of TEs

# %% [code]
correlation_df_families = pd.read_csv(
    "Signal_vs_divergence_correlation_by_lines_mods_families.csv", index_col=0
).T
family_names = list(correlation_df_families.columns[2:])
correlation_df_families[family_names] = correlation_df_families[family_names].apply(
    pd.to_numeric, errors="coerce"
)
correlation_df_families

# %% [markdown]
### TE classes palette creation

# %% [code]
class_names = ["LINE", "LTR", "SINE", "DNA", "Retroposon", "RC"]
class_names_p = {
    "LINE": "#cc660b",
    "LTR": "#70453c",
    "SINE": "#ab1f20",
    "DNA": "#195f90",
    "Retroposon": "#765297",
    "RC": "#238023",
}

cell_line_p = {
    "Caco-2": "saddlebrown",
    "SJCRH30": "darkgreen",
    "BE2C": "grey",
    "SJSA1": "teal",
    "HAP-1": "royalblue",
    "HL-60": "cyan",
    "MG63": "lime",
    "VCaP": "gold",
    "C4-2B": "orangered",
    "RWPE2": "orange",
    "22Rv1": "lightcoral",
    "RWPE1": "deeppink",
}

["#195f90", "#cc660b", "#238023", "#ab1f20", "#765297", "#70453c"]

# %% [code]
import os

import matplotlib.patches as mpatches
import matplotlib.pyplot as plt

# 1. Create a list of 'Patch' objects for the legend handles
# A Patch is just a colored rectangle
patches = [
    mpatches.Patch(color=color, label=label) for label, color in class_names_p.items()
]

# 2. Create a figure and a single axis
# We can make the figure small, as we will crop it tightly
fig, ax = plt.subplots(figsize=(4, 2))

# 3. Create the legend on the axis
# We place it in the 'center' of the invisible axis
legend = ax.legend(
    handles=patches,
    loc="center",
    frameon=False,  # Remove the legend box frame
    fontsize=12,
    ncol=1,  # Arrange in a single column
)

# 4. Turn off the axis (spines, ticks, etc.)
ax.axis("off")

# 5. Define the save path
save_path = "./plots/class_names_legend.svg"

# Ensure the 'plots' directory exists (if you want to save it there)
# os.makedirs(os.path.dirname(save_path), exist_ok=True)

# 6. Save the figure as an SVG
# bbox_inches='tight' is the key to cropping the output
# to the legend's bounding box.
plt.savefig(
    save_path,
    format="svg",
    bbox_inches="tight",
    pad_inches=0.1,  # Add a small amount of padding
    transparent=True,  # Make the background transparent
)

plt.close(fig)  # Close the figure to free memory

print(f"Legend saved successfully to {save_path}")

# %% [code]
1072 / 756

# %% [code]
import matplotlib.colors as mcolors

cmap = matplotlib.cm.get_cmap("tab10")
# Access the .colors attribute (a list of RGBA tuples)
rgba_colors = cmap.colors

# Convert each RGBA tuple to a hex string
hex_colors = [mcolors.to_hex(c) for c in rgba_colors]

# Print the list of hex color strings
print(hex_colors)

# %% [code]
families_by_classes = pd.read_csv("families_by_classes_TE.csv").dropna()

families_by_classes = families_by_classes.set_index("family_name").reindex(family_names)
# families_by_classes = families_by_classes[families_by_classes.index!='NaN']
#
families_by_classes

# %% [markdown]
### Clustermap of TE families epigenetic signal correlation with divergence

# %% [code]
correlation_df_families = pd.read_csv(
    "Signal_vs_divergence_correlation_by_lines_mods_families.csv", index_col=0
).T
family_names = list(correlation_df_families.columns[2:])
correlation_df_families[family_names] = correlation_df_families[family_names].apply(
    pd.to_numeric, errors="coerce"
)
correlation_df_families.index = (
    correlation_df_families.cell_line + " " + correlation_df_families.modification
)

correlation_df_families

# %% [code]
plt.rcParams["svg.fonttype"] = "none"

# --- 1. Prepare Row Colors ---
# Map the 'cell_line' and 'modification' columns to their respective colors


mapped_cell_colors = correlation_df_families["cell_line"].map(cell_line_p)
mapped_mod_colors = correlation_df_families["modification"].map(modification_p_richer)

families_counts = pd.read_csv("repeat_masker_family_name_counts.csv")
# families_counts

families_by_classes = pd.read_csv("families_by_classes_TE.csv").dropna()

families_by_classes = families_by_classes.set_index("family_name").reindex(family_names)
mapped_families_colors = families_by_classes["class_name"].map(class_names_p)

# Create a DataFrame for the row_colors argument
# The column names (e.g., 'Cell Line') will appear as labels.
row_colors_df = pd.DataFrame(
    {"Cell Line": mapped_cell_colors, "Modification": mapped_mod_colors}
)
# Important: The index must align with the data being plotted
row_colors_df.index = correlation_df_families.index


# --- 2. Plot the Clustermap ---
data_to_plot = correlation_df_families[family_names].T
data_to_plot

g = sns.clustermap(
    data_to_plot,
    cmap="coolwarm",
    # Goal 1: Customize figsize
    figsize=(11, 13),
    # Goal 2 & 3: Add row colors using the mapped palettes
    # We use row_colors, as 'cell_line' and 'modification' describe the rows
    col_colors=row_colors_df[["Modification", "Cell Line"]],
    row_colors=mapped_families_colors,
    # Other aesthetic improvements
    # vmin=-1, vmax=1,          # Set colorbar limits for correlation
    linewidths=0.5,
    cbar_kws={"label": "Pearson Correlation"},
    # metric='correlation'      # Use correlation for clustering distance
)

# --- 3. Save the Figure ---
save_path = "./plots/signal_to_divergence_corr_clustermap_families.svg"


# Goal 4: Save as SVG
g.savefig(save_path, format="svg")


plt.show()

# %% [code]
import os

import matplotlib.pyplot as plt
import numpy as np
import pandas as pd
import seaborn as sns
from matplotlib.gridspec import GridSpec, GridSpecFromSubplotSpec


# --- Setup: Helper Class for Colored Logging ---
class Logger:
    """A simple logger for pretty, colored console output."""

    BLUE = "\033[94m"
    GREEN = "\033[92m"
    YELLOW = "\033[93m"
    ENDC = "\033[0m"

    @staticmethod
    def info(message):
        print(f"{Logger.BLUE}INFO: {message}{Logger.ENDC}")

    @staticmethod
    def success(message):
        print(f"{Logger.GREEN}SUCCESS: {message}{Logger.ENDC}")

    @staticmethod
    def warning(message):
        print(f"{Logger.YELLOW}WARNING: {message}{Logger.ENDC}")

    @staticmethod
    def step(message):
        print(f"  -> {message}")


Logger.info("Setting up the data and palettes...")

families_counts = pd.read_csv("repeat_masker_family_name_counts.csv")
families_counts.columns = ["family_name", "count"]
families_counts["count"] = np.log10(families_counts["count"])

families_by_classes = pd.read_csv("families_by_classes_TE.csv").dropna()

# --- 1. Prepare Row Colors ---
Logger.step("Preparing row and column colors...")
# Map the 'cell_line' and 'modification' columns to their respective colors
mapped_cell_colors = correlation_df_families["cell_line"].map(cell_line_p)
mapped_mod_colors = correlation_df_families["modification"].map(modification_p_richer)

families_by_classes = families_by_classes.set_index("family_name").reindex(family_names)
mapped_families_colors = families_by_classes["class_name"].map(class_names_p)

# Create a DataFrame for the row_colors argument
row_colors_df = pd.DataFrame(
    {"Cell Line": mapped_cell_colors, "Modification": mapped_mod_colors}
)
row_colors_df.index = correlation_df_families.index

# --- 2. Plot the Clustermap ---
Logger.step("Generating initial clustermap...")
data_to_plot = correlation_df_families.set_index(["cell_line", "modification"])[
    family_names
].T

g = sns.clustermap(
    data_to_plot,
    cmap="coolwarm",
    figsize=(12, 16),
    # Column colors (for cell line and modification)
    col_colors=row_colors_df[["Modification", "Cell Line"]],
    # Row colors (for repeat class)
    row_colors=mapped_families_colors,
    linewidths=0.5,
    cbar_kws={"label": "Pearson Correlation"},
    # vmin=-1, vmax=1 # Good practice for correlation
)

# --- 3. Prepare Bar Plot Data ---
Logger.step("Preparing data for bar plot...")
# Get the reordered family names from the clustermap's row dendrogram
reordered_indices = g.dendrogram_row.reordered_ind
reordered_families = data_to_plot.index[reordered_indices]

# Load and prepare the counts data
# Set index to 'family_name' to allow reindexing
families_counts = families_counts.set_index("family_name")

# Reorder the counts to match the clustermap's row order
counts_reordered = families_counts.reindex(reordered_families)

# --- 4. Resize Axes and Add Bar Plot ---
Logger.step("Resizing axes and adding bar plot...")

# Get current positions
heatmap_pos = g.ax_heatmap.get_position()
cbar_pos = g.cax.get_position()

# Define new widths
bar_width = 0.08  # 8% of figure width for bars
space = 0.01

# Shrink heatmap to make space
new_heatmap_width = heatmap_pos.width - bar_width - space
g.ax_heatmap.set_position(
    [heatmap_pos.x0, heatmap_pos.y0, new_heatmap_width, heatmap_pos.height]
)

# Add new axis for the bar plot
bar_ax_left = heatmap_pos.x0 + new_heatmap_width + space
bar_ax = g.fig.add_axes([bar_ax_left, heatmap_pos.y0, bar_width, heatmap_pos.height])

# Move colorbar to the right of the new bar plot
new_cbar_left = bar_ax_left + bar_width + space
g.cax.set_position(
    [new_cbar_left, cbar_pos.y0, cbar_pos.width, cbar_pos.height]
)  # Keep original width

# --- 5. Plot the horizontal bar chart ---
sns.barplot(
    y=counts_reordered.index,
    x=counts_reordered["count"],  # Assumes 'count' column
    orient="h",
    ax=bar_ax,
    color="#888888",  # Neutral gray
)

# --- 6. Tidy up all axes ---
Logger.step("Tidying up final plot...")

# Tidy the new bar plot axis
bar_ax.set_yticks([])  # Remove y-tick labels
bar_ax.set_ylabel("")  # Remove y-axis label
bar_ax.set_xlabel("Copy Count")
bar_ax.spines["top"].set_visible(False)
bar_ax.spines["right"].set_visible(False)
bar_ax.spines["left"].set_visible(False)

# Tidy the main heatmap
g.ax_heatmap.set_yticklabels([])  # Remove family names (now redundant)
g.ax_heatmap.set_ylabel("")  # Remove row label

# --- 7. Save the Figure ---
save_path = "./plots/signal_to_divergence_corr_clustermap_families_with_barplot.svg"
os.makedirs(os.path.dirname(save_path), exist_ok=True)  # Ensure dir exists

# Save as SVG
g.savefig(save_path, format="svg")
Logger.success(f"Clustermap saved to {save_path}")

plt.show()

# %% [code]
(correlation_df_families[family_names].abs() > 0.3).sum().sum()

# %% [code]
families_counts = pd.read_csv("repeat_masker_family_name_counts.csv")
families_counts

# %% [code]
1164096

# %% [code]
correlation_df_individuals = pd.read_csv(
    "Signal_vs_divergence_correlation_by_lines_mods_individuals.csv", index_col=0
).T
individual_names = list(correlation_df_individuals.columns[2:])
individuals_counts = pd.read_csv("individuals_counts.csv", index_col=0)
individual_names = individuals_counts[individuals_counts > 1000].dropna().index

correlation_df_individuals[individual_names] = correlation_df_individuals[
    individual_names
].apply(pd.to_numeric, errors="coerce")
correlation_df_individuals.index = (
    correlation_df_individuals.cell_line + " " + correlation_df_individuals.modification
)
correlation_df_individuals

# %% [markdown]
### Clustering vs average divergence barplot

# %% [code]
reordered_indices = g.dendrogram_row.reordered_ind
reordered_families = data_to_plot.index[reordered_indices]
reordered_families

# %% [code]
repeat_masker = pd.read_csv("T2T_repeat_masker_processed.csv")
divergence_average = (
    repeat_masker[["score", "family_name"]].groupby("family_name").mean()
)
divergence_average

# %% [code]
# Get the reordered family names from the clustermap's row dendrogram
reordered_indices = g.dendrogram_row.reordered_ind
reordered_families = data_to_plot.index[reordered_indices]

# Load and prepare the counts data
# Set index to 'family_name' to allow reindexing


# Reorder the counts to match the clustermap's row order
divergence_average = divergence_average.reindex(reordered_families)

# Define new widths
bar_width = 0.08  # 8% of figure width for bars
space = 0.01

fig, ax = plt.subplots(figsize=(1, 10))
# --- 5. Plot the horizontal bar chart ---
sns.barplot(
    y=divergence_average.index,
    x=divergence_average["score"],  # Assumes 'count' column
    orient="h",
    # ax=bar_ax,
    color="#888888",  # Neutral gray
)

save_path = "./plots/average_divergence_families_barplot.svg"
# Save as SVG
plt.savefig(save_path, format="svg")

plt.show()

# %% [markdown]
### Connection of correlation with average divergence and count by family

# %% [code]
correlation_df_families = pd.read_csv(
    "Signal_vs_divergence_correlation_by_lines_mods_families_detailed.csv", index_col=0
)
correlation_df_families['abs_r'] = correlation_df_families.r.abs()
corrs_average = correlation_df_families[['cell_line', 'modification', 'family_name', 'abs_r']].groupby(['family_name']).mean()


repeat_masker = pd.read_csv("T2T_repeat_masker_processed.csv")
divergence_average = (
    repeat_masker[["score", "family_name"]]
    .groupby("family_name")
    .mean()["score"]
    .reindex(corrs_average.index)
)

families_counts = pd.read_csv("repeat_masker_family_name_counts.csv")
families_counts.columns = ["family_name", "count"]
families_counts["log10_count"] = np.log10(families_counts["count"])
families_counts = families_counts.set_index("family_name").reindex(corrs_average.index)


families_by_classes = pd.read_csv("families_by_classes_TE.csv").dropna()

families_by_classes = families_by_classes.set_index("family_name").reindex(
    corrs_average.index
)

data_to_plot = pd.concat(
    [
        corrs_average,
        divergence_average,
        families_counts[["count", "log10_count"]],
        families_by_classes["class_name"],
    ],
    axis=1,
)
data_to_plot.columns = [
    "average_abs_corr",
    "average_divergence",
    "count",
    "log10_count",
    "class_name",
]
data_relevant = data_to_plot.copy()
data_to_plot

# %% [code]
sns.scatterplot(
    data=data_to_plot,
    hue="class_name",
    x="log10_count",
    y="average_abs_corr",
    palette=class_names_p,
)

# %% [code]
sns.pairplot(
    data=data_to_plot[
        [
            "average_abs_corr",
            "average_divergence",
            "log10_count",
            "class_name",
        ]
    ],
    hue="class_name",
)

# %% [code]
?curve_fit

# %% [code]
 data_to_plot[["log10_count", "average_abs_corr"]]

# %% [code]
import matplotlib.pyplot as plt
import numpy as np
import pandas as pd
import seaborn as sns
from scipy import stats
from scipy.optimize import curve_fit


# %% [code]
??curve_fit

# %% [code]
import matplotlib.pyplot as plt
import numpy as np
import pandas as pd
import seaborn as sns
from scipy import stats
from scipy.optimize import curve_fit

plt.rcParams["svg.fonttype"] = "none"  # For clean SVG text


# --- 2. Define the sigmoidal function ---
def sigmoid(x, L, k, x0):
    """
    Sigmoidal (logistic) function.
    L: maximum value (the asymptote)
    k: steepness of the curve
    x0: the x-value of the sigmoid's midpoint
    """
    return L / (1 + np.exp(-k * (x - x0)))


# --- 3. Prepare data and perform the fit ---
print("Fitting sigmoidal curve...")

# Drop NaNs to ensure alignment and prevent errors
fit_data = data_to_plot[["log10_count", "average_abs_corr"]].dropna()
x_data = fit_data["log10_count"]
y_data = fit_data["average_abs_corr"]

# Provide initial guesses for the parameters [L, k, x0]
# L: guess the max value of y
# k: guess 1 (steepness)
# x0: guess the median value of x
p0 = [y_data.max(), 1, x_data.median()]

try:
    # Fit the curve
    popt, pcov = curve_fit(sigmoid, x_data, y_data, p0=p0, maxfev=5000)
    L, k, x0 = popt
    fit_successful = True

    # --- 4. Calculate Statistics (R-squared and p-value) ---
    # Get the predicted y-values from the fit
    y_pred = sigmoid(x_data, *popt)

    # Calculate R-squared
    # (using Pearson correlation between actual and predicted)
    corr, p_value = stats.pearsonr(y_data, y_pred)
    r_squared = corr**2

    # Format p-value for display
    if p_value < 0.001:
        p_val_str = "p < 0.001"
    else:
        p_val_str = f"p = {p_value:.3f}"

    # Create the equation string for the plot
    equation_text = (
        f"y = {L:.2f} / (1 + exp(-{k:.2f} $\\cdot$ (x - {x0:.2f})))\n"
        f"$R^2$ = {r_squared:.3f}\n"
        f"{p_val_str}"
    )

except RuntimeError:
    # This happens if the curve fit fails to converge
    print("Sigmoidal curve fit failed to converge.")
    fit_successful = False
    equation_text = "Fit failed to converge"


# --- 5. Create the Plot ---
print("Generating plot...")
# Create the figure and axes (for figsize customization)
fig, ax = plt.subplots(figsize=(7, 4))

# Draw the scatterplot onto the axis
sns.scatterplot(
    data=data_to_plot,
    hue="class_name",
    x="log10_count",
    y="average_abs_corr",
    palette=class_names_p,
    ax=ax,
    alpha=0.7,
)

if fit_successful:
    # --- 6. Plot the Fitted Curve ---
    # Create a smooth x-axis for the curve
    x_fit = np.linspace(x_data.min(), x_data.max(), 200)
    y_fit = sigmoid(x_fit, *popt)

    # Plot the curve
    ax.plot(x_fit, y_fit, color="black", ls="--", lw=2.5, label="Sigmoid Fit")

    # --- 7. Add the Equation Textbox ---
    ax.text(
        0.05,
        0.95,  # Position in (x, y) coordinates (0.05=5% from left, 0.95=95% from top)
        equation_text,
        transform=ax.transAxes,  # Use axis-relative coordinates
        fontsize=10,
        verticalalignment="top",
        ha="left",
        # Add a light background box
        bbox=dict(facecolor="white", alpha=0.7, boxstyle="round,pad=0.5"),
    )

# --- 8. Tidy Up and Save ---
ax.legend(loc="center left", bbox_to_anchor=(1, 0.5))
ax.set_title("Sigmoidal Fit of Correlation vs. Count", fontsize=16)
ax.set_xlabel("Count", fontsize=12)
ax.set_ylabel("Average Absolute Correlation", fontsize=12)
ax.set_ylim((0, 0.225))

plt.tight_layout(rect=[0, 0, 0.85, 1])  # Make space for the legend

# Save to SVG
save_path = "./plots/abs_average_corr_vs_count_TE_random.svg"
fig.savefig(save_path, format="svg")
print(f"Plot saved to {save_path}")

plt.show()

# %% [markdown]
#### Random correlations

# %% [code]
correlation_df_families = pd.read_csv(
    "Signal_vs_divergence_correlation_by_lines_mods_families_detailed_random.csv", index_col=0
)
correlation_df_families['abs_r'] = correlation_df_families.r.abs()
corrs_average = correlation_df_families[['cell_line', 'modification', 'family_name', 'abs_r']].groupby(['family_name']).mean()


repeat_masker = pd.read_csv("T2T_repeat_masker_processed.csv")
divergence_average = (
    repeat_masker[["score", "family_name"]]
    .groupby("family_name")
    .mean()["score"]
    .reindex(corrs_average.index)
)

families_counts = pd.read_csv("repeat_masker_family_name_counts.csv")
families_counts.columns = ["family_name", "count"]
families_counts["log10_count"] = np.log10(families_counts["count"])
families_counts = families_counts.set_index("family_name").reindex(corrs_average.index)


families_by_classes = pd.read_csv("families_by_classes_TE.csv").dropna()

families_by_classes = families_by_classes.set_index("family_name").reindex(
    corrs_average.index
)

data_control = pd.concat(
    [
        corrs_average,
        divergence_average,
        families_counts[["count", "log10_count"]],
        families_by_classes["class_name"],
    ],
    axis=1,
)
data_control.columns = [
    "average_abs_corr",
    "average_divergence",
    "count",
    "log10_count",
    "class_name",
]
data_control

# %% [markdown]
#### Compare random and relevant via F-Test for Nested Models

# %% [code]
import numpy as np
import pandas as pd
from scipy.optimize import curve_fit
from scipy import stats

def sigmoid_decay(x, L, k, x0):
    """
    Sigmoidal decay (inverse logistic) function.
    L: maximum value (the asymptote)
    k: steepness of the curve
    x0: the x-value of the sigmoid's midpoint
    """
    return L / (1 + np.exp(k * (x - x0)))

def get_rss(y_true, y_pred):
    """Calculates the Residual Sum of Squares."""
    return np.sum((y_true - y_pred)**2)

def fit_sigmoid(x_data, y_data):
    """
    Fits a sigmoid_decay curve to X and Y data.
    Returns the optimal parameters and the RSS.
    """
    # Provide initial guesses for [L, k, x0]
    # L: guess the max value of y
    # k: guess 1.0 (steepness)
    # x0: guess the median value of x
    p0 = [np.max(y_data), 1.0, np.median(x_data)]
    
    try:
        popt, _ = curve_fit(sigmoid_decay, x_data, y_data, p0=p0, maxfev=5000)
        # Calculate RSS
        y_pred = sigmoid_decay(x_data, *popt)
        rss = get_rss(y_data, y_pred)
        return popt, rss
    except RuntimeError as e:
        print(f"Error: Curve fitting failed - {e}")
        return None, np.inf


# --- 2. Combine Datasets ---
data_relevant['group'] = 'relevant'
data_control['group'] = 'control'
data_all = pd.concat([data_relevant, data_control]).reset_index(drop=True)

# Define X and Y for all datasets
x_all = data_all['log10_count']
y_all = data_all['average_abs_corr']

x_rel = data_relevant['log10_count']
y_rel = data_relevant['average_abs_corr']

x_con = data_control['log10_count']
y_con = data_control['average_abs_corr']

N_all = len(data_all)
N_rel = len(data_relevant)
N_con = len(data_control)

# --- 3. Fit the Null Model (H₀) ---
# H₀: One single curve fits all data.
print("--- Fitting Null Model (H₀) ---")
popt_null, RSS_null = fit_sigmoid(x_all, y_all)
if RSS_null == np.inf:
    print("FATAL: Could not fit the Null Model. Stopping.")
else:
    K_null = 3 # 3 parameters (L, k, x0)
    df_null = N_all - K_null
    print(f"  - RSS_null: {RSS_null:.4f}")
    print(f"  - df_null (N - K_null): {df_null}")


    # --- 4. Fit the Alternative Model (H₁) ---
    # H₁: Two separate curves (one for each group) fit the data.
    print("\n--- Fitting Alternative Model (H₁) ---")
    popt_rel, RSS_relevant = fit_sigmoid(x_rel, y_rel)
    popt_con, RSS_control = fit_sigmoid(x_con, y_con)
    
    if RSS_relevant == np.inf or RSS_control == np.inf:
        print("FATAL: Could not fit one of the alternative models. Stopping.")
    else:
        RSS_alt = RSS_relevant + RSS_control
        K_alt = 6 # 3 params for 'relevant' + 3 params for 'control'
        df_alt = N_all - K_alt  # (or (N_rel - 3) + (N_con - 3))
        
        print(f"  - RSS_relevant: {RSS_relevant:.4f}")
        print(f"  - RSS_control: {RSS_control:.4f}")
        print(f"  - RSS_alt (combined): {RSS_alt:.4f}")
        print(f"  - df_alt (N - K_alt): {df_alt}")

        
        # --- 5. Perform the F-Test ---
        print("\n--- F-Test Results ---")
        
        # Degrees of freedom for the test
        df1 = df_null - df_alt  # Number of *extra* parameters (K_alt - K_null = 3)
        df2 = df_alt            # Degrees of freedom of the full (alternative) model
        
        # Calculate the F-statistic
        F_statistic = ((RSS_null - RSS_alt) / df1) / (RSS_alt / df2)
        
        # Get the p-value from the F-distribution
        # stats.f.sf is the survival function (1 - cdf), which is what we want
        p_value = stats.f.sf(F_statistic, df1, df2)
        
        print(f"  - F-statistic: {F_statistic:.4f}")
        print(f"  - p-value: {p_value:.4e}") # Scientific notation
        
        print("\n--- Conclusion ---")
        if p_value < 0.05:
            print(f"p < 0.05. We REJECT the null hypothesis.")
            print("The two sigmoid curves are STATISTICALLY SIGNIFICANTLY DIFFERENT.")
        else:
            print(f"p >= 0.05. We FAIL to reject the null hypothesis.")
            print("The two curves are NOT statistically different.")

# %% [markdown]
This is an excellent statistical question. The situation you're in—where even your "random" control shows a pattern—is a very common and important finding.Your images are perfect examples. The "Sigmoidal Fit" (Image 2, $R^2 = 0.989$) fits beautifully, while your "Biologically Relevant" data (Image 1, $R^2 = 0.579$) is much, much noisier.The core question is: "Is the relationship between Count and Correlation in my relevant data significantly different from the relationship in my control data?"You can't just compare p-values or $R^2$ values directly. You need to test if the parameters that define the curves are statistically different.Here is the most rigorous method to do this, often called a Sum-of-Squares F-test for non-linear model comparison.The "Gold Standard" Method: F-Test for Nested ModelsThis method tests whether fitting two separate sigmoid curves (one for "Relevant" and one for "Control") is significantly better than fitting just one curve to all the data combined.Here is the step-by-step logic:1. Combine Your DataFirst, combine both datasets into a single, larger dataset. You must add a "dummy" or "indicator" variable to keep track of which group each data point belongs to.log10_countaverage_abs_corrgroup2.10.125'relevant'3.40.045'relevant'.........2.20.115'control'3.50.030'control'.........2. Fit the "Null" Model (H₀)The "Null Hypothesis" (H₀) is that there is no difference between the groups.To model this, you fit one single sigmoid curve to all the data combined, ignoring the group column.y = L / (1 + exp(-k * (x - x0)))After fitting, you calculate the Residual Sum of Squares (RSS) for this one model. Let's call it RSS_null.You also record the degrees of freedom: df_null = (total_N) - 3 (where N is the total number of points, and 3 is the number of parameters: L, k, x0).3. Fit the "Alternative" Model (H₁)The "Alternative Hypothesis" (H₁) is that the groups are different.To model this, you fit two separate sigmoid curves:Fit one curve only to the 'relevant' data. Get its RSS_relevant.Fit another curve only to the 'control' data. Get its RSS_control.Your total residual sum of squares is the sum of these two: RSS_alt = RSS_relevant + RSS_control.You record the degrees of freedom: df_alt = (n_relevant - 3) + (n_control - 3) or simply (total_N) - 6.4. Compare the Models with an F-TestYou now test if the reduction in error (from RSS_null to RSS_alt) is statistically significant.You calculate an F-statistic:$$F = \frac{(RSS_{null} - RSS_{alt}) / (df_{null} - df_{alt})}{(RSS_{alt}) / (df_{alt})}$$RSS_null - RSS_alt = The improvement in fit.df_null - df_alt = The number of extra parameters you used. (e.g., 6 parameters in the 'alt' model - 3 in the 'null' model = 3).(RSS_alt) / (df_alt) = The variance of the full model.You then calculate a p-value from this F-statistic (using the F-distribution with df1 = (df_null - df_alt) and df2 = df_alt).InterpretationIf p-value < 0.05: You have a statistically significant result. You can reject the null hypothesis. This proves that fitting two separate curves is significantly better than fitting one. You can conclude the two sigmoid curves are statistically different.If p-value > 0.05: You cannot conclude the curves are different. The extra complexity of the two-curve model was not justified by the data.

# %% [markdown]
MOVE TO METHODS

# %% [markdown]
### Analyze correlation coefficients with limits

# %% [code]
correlation_df_families = pd.read_csv(
    "Signal_vs_divergence_correlation_by_lines_mods_families_detailed.csv", index_col=0
)
correlation_df_families

# %% [code]
from statsmodels.stats.multitest import multipletests

reject, p_adjusted, _, _ = multipletests(
    correlation_df_families["p_value"], alpha=0.05, method="fdr_bh"
)

correlation_df_families["q_value"] = p_adjusted
p_adjusted

# %% [code]
correlation_df_families[correlation_df_families["ci_low"] > 0.2]

# %% [code]
correlation_df_families[correlation_df_families["ci_high"] < -0.2]

# %% [code]


# %% [code]
len(correlation_df_families[correlation_df_families["r"].abs() > 0.2])

# %% [code]
len(correlation_df_families)

# %% [code]
76 / 1716

# %% [markdown]
### Visualize top and bottom correlations for families, marks and cell lines

# %% [code]
correlation_df_families = pd.read_csv(
    "Signal_vs_divergence_correlation_by_lines_mods_families_detailed.csv", index_col=0
)

top_corrs = pd.concat(
    [
        correlation_df_families[correlation_df_families["ci_low"] > 0.2],
        correlation_df_families[correlation_df_families["ci_high"] < -0.2],
    ]
)
top_corrs = top_corrs[
    ~((top_corrs["cell_line"] == "C4-2B") & (top_corrs["modification"] == "CTCF") & (top_corrs["family_name"] == "SVA"))
]

top_corrs = top_corrs[
    ~((top_corrs["cell_line"] == "RWPE1") & (top_corrs["modification"] == "H3K27ac") & (top_corrs["family_name"] == "SVA"))
]

top_corrs

# %% [code]
def pearsonr_ci(x, y, alpha=0.05):
    """
    Calculates Pearson r, p-value, and 95% CI using Fisher z-transformation.
    This works on all versions of SciPy.
    """
    # Calculate r and p-value
    r, p = stats.pearsonr(x, y)
    n = len(x)

    # Fisher z-transformation
    z = np.arctanh(r)

    # Standard error of z
    se_z = 1.0 / np.sqrt(n - 3)

    # Get the z-score for the desired confidence level
    z_crit = stats.norm.ppf(1 - alpha / 2)  # This is 1.96 for 95% CI

    # Calculate CI for z
    z_low = z - z_crit * se_z
    z_high = z + z_crit * se_z

    # Back-transform z CI to r CI
    r_low = np.tanh(z_low)
    r_high = np.tanh(z_high)

    return r, p, r_low, r_high

# %% [code]
import os

import matplotlib.pyplot as plt
import numpy as np
import pandas as pd
import seaborn as sns
from matplotlib.gridspec import GridSpec, GridSpecFromSubplotSpec
from tqdm import tqdm  # For progress bars

correlation_df_families = pd.read_csv(
    "Signal_vs_divergence_correlation_by_lines_mods_families_detailed.csv", index_col=0
)

top_corrs = pd.concat(
    [
        correlation_df_families[correlation_df_families["ci_low"] > 0.2],
        correlation_df_families[correlation_df_families["ci_high"] < -0.2],
    ]
)
top_corrs = top_corrs[
    ~((top_corrs["cell_line"] == "C4-2B") & (top_corrs["modification"] == "CTCF") & (top_corrs["family_name"] == "SVA"))
]

top_corrs = top_corrs[
    ~((top_corrs["cell_line"] == "RWPE1") & (top_corrs["modification"] == "H3K27ac") & (top_corrs["family_name"] == "SVA"))
]

plt.rcParams["svg.fonttype"] = "none"


# --- Helper Class for Colored Logging ---
class Logger:
    """A simple logger for pretty, colored console output."""

    BLUE = "\033[94m"
    GREEN = "\033[92m"
    YELLOW = "\033[93m"
    ENDC = "\033[0m"

    # Note: Use tqdm.write() for all logging inside loops
    @staticmethod
    def info(message):
        print(f"{Logger.BLUE}INFO: {message}{Logger.ENDC}")

    @staticmethod
    def success(message):
        print(f"{Logger.GREEN}SUCCESS: {message}{Logger.ENDC}")

    @staticmethod
    def warning(message):
        print(f"{Logger.YELLOW}WARNING: {message}{Logger.ENDC}")

    # Static versions to return the string for tqdm.write()
    @staticmethod
    def s_success(message):
        return f"{Logger.GREEN}SUCCESS: {message}{Logger.ENDC}"

    @staticmethod
    def s_warning(message):
        return f"{Logger.YELLOW}WARNING: {message}{Logger.ENDC}"


save_path_svg = f"plots/Divergence_vs_Signal_by_families_sign_correlations.svg"

figure = plt.figure(figsize=(10, 6))

N_ROWS = 3
N_COLS = 4

# Increased spacing for independent labels
outer_grid = GridSpec(N_ROWS, N_COLS, figure=figure, wspace=0.4, hspace=0.5)

# --- Removed shared axes lists ---

# --- Add a cache for data loading ---
data_cache = {}

# --- Use a more Pythonic loop ---
for i, (idx, row) in enumerate(
    tqdm(top_corrs.iterrows(), total=len(top_corrs), desc="Plotting Top 10")
):

    # Calculate grid position
    r = i // N_COLS
    c = i % N_COLS

    # Get data from the row
    family_name = row["family_name"]
    cell_line = row["cell_line"]
    modification = row["modification"]

    filepath = f"./epigenomic_files/{cell_line}.{modification}.chm13v2.0.mapped_on_repeat_masker.bedGraph"

    # --- Check cache before loading ---
    if filepath in data_cache:
        tqdm.write(f"  -> Found in cache: {filepath}")
        sample_enrichment = data_cache[filepath]
    else:
        # --- Data Loading ---
        tqdm.write(f"  -> Loading data from: {filepath}")
        try:
            sample_enrichment = pd.read_csv(
                filepath,
                sep="\t",
                header=None,
            )
            data_cache[filepath] = sample_enrichment
            tqdm.write(Logger.s_success("File loaded and cached."))

        except FileNotFoundError:
            tqdm.write(Logger.s_warning(f"File not found, skipping: {filepath}"))
            ax = figure.add_subplot(outer_grid[r, c])
            ax.axis("off")
            continue
        except Exception as e:
            tqdm.write(
                Logger.s_warning(
                    f"An unexpected error occurred while reading {filepath}: {e}"
                )
            )
            ax = figure.add_subplot(outer_grid[r, c])
            ax.axis("off")
            continue

        # --- Process data (only needs to be done once, on load) ---
        sample_enrichment.columns = [
            "Chromosome",
            "Start",
            "End",
            "Divergence",
            "Name",
            "Family",
            "Class",
            "Signal",
        ]
        sample_enrichment["Signal"] = pd.to_numeric(
            sample_enrichment["Signal"], errors="coerce"
        )
        sample_enrichment["Divergence"] = pd.to_numeric(
            sample_enrichment["Divergence"], errors="coerce"
        )
        data_cache[filepath] = sample_enrichment

    # --- Query must be done every time ---
    query_local = f'Family == "{family_name}"'
    sample_enrichment_local = sample_enrichment.query(query_local)

    # --- Create *single* axis for this plot ---
    ax_main = figure.add_subplot(outer_grid[r, c])

    # --- Plot the data ---
    sns.histplot(
        data=sample_enrichment_local,
        x="Divergence",
        y="Signal",
        bins=48,
        pmax=0.9,
        cmap="viridis",
        ax=ax_main,
    )

    # --- Tidy up: Main plot ticks/labels ---
    ax_main.set_facecolor("black")
    ax_main.set_xlabel("Divergence", fontsize=10)
    ax_main.set_ylabel(f"{cell_line}\n{modification}", fontsize=10)

    # Set spine colors to white

    # --- Calculate correlation ---
    clean_data = sample_enrichment_local.dropna(subset=["Divergence", "Signal"])
    if len(clean_data) > 3:  # Need n > 3 for the 1/sqrt(n-3) formula
        try:
            # Use the new helper function
            corr_coef, p_val, ci_low, ci_high = pearsonr_ci(
                clean_data["Divergence"], clean_data["Signal"]
            )

            # Create the string for the title
            corr_str = f"r = {corr_coef:.3f} (95% CI: [{ci_low:.3f}, {ci_high:.3f}])"
        except Exception as e:
            # Catch any math errors (e.g., if variance is zero)
            corr_coef = np.nan
            corr_str = "r = N/A (calc error)"
    else:
        corr_coef = np.nan  # Set to nan
        corr_str = "r = N/A (n <= 3)"
    # --- Set the title on the main axis ---
    ax_main.set_title(f"{family_name}\n{corr_str}", fontsize=10, pad=10)


# --- Ensure plots directory exists ---
os.makedirs(os.path.dirname(save_path_svg), exist_ok=True)
figure.tight_layout()  # Add tight_layout to prevent label overlap
figure.savefig(save_path_svg, format="svg", facecolor="white", transparent=True)
plt.show()

Logger.success(f"Plot saved to {save_path_svg}")

# %% [markdown]
### Compare correlations by class, cell line, modification

# %% [code]
families_by_classes = pd.read_csv("families_by_classes_TE.csv").dropna()


correlation_df_families = pd.read_csv(
    "Signal_vs_divergence_correlation_by_lines_mods_families_detailed.csv", index_col=0
).merge(families_by_classes[['class_name', 'family_name']])
correlation_df_families["transformed_corr"] = np.arctanh(correlation_df_families.r)
correlation_df_families["r_abs"] = correlation_df_families.r.abs()
correlation_df_families

# %% [code]
import os
import matplotlib.pyplot as plt
import numpy as np
import pandas as pd
import seaborn as sns
from scipy import stats # We need this for the stats test
from statannotations.Annotator import Annotator

# Set the font type for a clean SVG
plt.rcParams['svg.fonttype'] = 'none'

x_name='cell_line'
# --- 2. Create the Plot ---
figure, ax = plt.subplots(
    figsize=(8, 7), # Increased height for stats
)

# --- 3. Plot the Boxplot ---
sns.boxplot(
    data=correlation_df_families,
    x=x_name,
    y="r_abs",
    ax=ax,
    palette=cell_line_p,
    boxprops={'alpha': 1}, # Make boxes slightly transparent,
    showfliers=False
)

# --- 4. Add the Scatter Plot (Stripplot) ---
# This plots the individual points over the boxes
sns.stripplot(
    data=correlation_df_families,
    x=x_name,
    y="r_abs",
    ax=ax,
    color="black",  # Color for the points
    size=3,         # Size of the points
    alpha=0.5,      # Transparency
    jitter=True, # Spread points horizontally
    #hue='modification',
    #palette=modification_p_richer
)

# Get the order of boxes as they appear on the plot
plot_order = [tick.get_text() for tick in ax.get_xticklabels()]

# Define the pairs you want to compare.
# This example compares all groups to the first group ("CTCF")

box_pairs = [(plot_order[i], plot_order[i+1]) for i in range(0,len(plot_order)-1)]

# You can also define custom pairs, e.g.:
# box_pairs = [
#     ("CTCF", "H3K27ac"), 
#     ("H3K27ac", "H3K4me3")
# ]

annotator = Annotator(
    ax,
    box_pairs,
    data=correlation_df_families,
    x=x_name,
    y="r_abs",
    order=plot_order # Pass the plot order
)

annotator.configure(
    test='Mann-Whitney', # Specify the test
    text_format='star',  # Use stars for significance
    loc='outside',       # Place annotations outside the plot area
    verbose=2            # Show p-values in the console
)

# Apply the test and draw the annotations
annotator.apply_and_annotate()
# --- 6. Tidy Up and Save ---

ax.set_title(f"Absolute Correlation by {x_name}", fontsize=10)
ax.set_xlabel(x_name, fontsize=10)
ax.set_ylabel("Absolute Pearson's r (|r|)", fontsize=10)

# Rotate x-axis labels for better readability
plt.xticks(rotation=0, ha='center')

# Ensure the plot layout is clean
figure.tight_layout()

# Save to SVG
save_path = f"./plots/corrs_boxplot_{x_name}.svg"
figure.savefig(
    save_path, 
    format="svg", 
    bbox_inches="tight",
    transparent=True
)

plt.show()


# %% [code]
1

# %% [code]
figure, ax = plt.subplots(
    figsize=(9, 4),
)
sns.boxplot(
    data=correlation_df_families,
    x="modification",
    y="r_abs",
    ax=ax,
    # hue='TE_class'
)

# %% [code]
figure, ax = plt.subplots(
    figsize=(9, 4),
)
sns.boxplot(
    data=correlation_df_families,
    x="class_name",
    y="r_abs",
    ax=ax,
    # hue='TE_class'
)

# %% [code]
figure, ax = plt.subplots(
    figsize=(9, 4),
)
sns.boxplot(
    data=correlation_df_families,
    x="cell_line",
    y="r_abs",
    ax=ax,
    # hue='TE_class'
)

# %% [code]


# %% [code]
import scipy

scipy.stats.pearsonr(clean_data["Divergence"], clean_data["Signal"])

# %% [markdown]
#### Curtosis and Shapiro test

# %% [code]
correlation_df_families = pd.read_csv(
    "Signal_vs_divergence_correlation_by_lines_mods_families.csv", index_col=0
).T
family_names = list(correlation_df_families.columns[2:])
correlation_df_families[family_names] = correlation_df_families[family_names].apply(
    pd.to_numeric, errors="coerce"
)
corrs_for_test = correlation_df_families.melt(
    id_vars=["cell_line", "modification"], value_vars=family_names
)
corrs_for_test.columns = ["cell_line", "modification", "TE_family", "correlation"]
corrs_for_test["transformed_corr"] = np.arctanh(corrs_for_test.correlation)

corrs_for_test = corrs_for_test.sort_values("correlation")

# %% [code]
sns.histplot(corrs_for_test.transformed_corr)

# %% [code]
stats.shapiro(corrs_for_test.transformed_corr)

# %% [code]
import scipy.stats as stats
??stats.shapiro

# %% [code]
stats.kurtosistest(corrs_for_test.transformed_corr)

# %% [code]
stats.kurtosis(corrs_for_test.transformed_corr)

# %% [markdown]
#### Levene test for comparison of variances

# %% [code]
from scipy.stats import levene

LINE = corrs_for_test["transformed_corr"][corrs_for_test["TE_class"] == "LINE"]
LTR = corrs_for_test["transformed_corr"][corrs_for_test["TE_class"] == "LTR"]
SINE = corrs_for_test["transformed_corr"][corrs_for_test["TE_class"] == "SINE"]
DNA = corrs_for_test["transformed_corr"][corrs_for_test["TE_class"] == "DNA"]
Retroposon = corrs_for_test["transformed_corr"][
    corrs_for_test["TE_class"] == "Retroposon"
]
RC = corrs_for_test["transformed_corr"][corrs_for_test["TE_class"] == "RC"]
non_sva = corrs_for_test["transformed_corr"][corrs_for_test["TE_class"] != "Retroposon"]


levene(LINE, LTR, SINE, DNA, Retroposon, RC)

# %% [code]
correlation_df_families

# %% [code]
from scipy.stats import levene

compare_list = [correlation_df_families["transformed_corr"][correlation_df_families["class_name"] == class_name] for class_name in correlation_df_families["class_name"].unique()] 

levene(*compare_list)

# %% [code]
compare_list = [correlation_df_families["transformed_corr"][correlation_df_families["cell_line"] == cell_line] for cell_line in correlation_df_families["cell_line"].unique()] 

levene(*compare_list)

# %% [code]
compare_list = [correlation_df_families["transformed_corr"][correlation_df_families["modification"] == modification] for modification in correlation_df_families["modification"].unique()] 

levene(*compare_list)

# %% [code]


# %% [markdown]
## Visualize means and SDs for families, their epigenetic signal

# %% [code]
data_by_family = pd.read_csv(
    "Average_signal_by_lines_mods_TE_families.csv", index_col=None
)
family_names = list(data_by_family.columns[:-2])
data_by_family[family_names] = data_by_family[family_names].apply(
    pd.to_numeric, errors="coerce"
)

data_by_family.index = data_by_family.cell_line + " " + data_by_family.modification
data_by_family

# %% [code]
mapped_cell_colors = data_by_family["cell_line"].map(cell_line_p)
mapped_mod_colors = data_by_family["modification"].map(modification_p_richer)

families_by_classes = pd.read_csv("families_by_classes_TE.csv").dropna()

families_by_classes = families_by_classes.set_index("family_name").reindex(family_names)
mapped_families_colors = families_by_classes["class_name"].map(class_names_p)

# Create a DataFrame for the row_colors argument
# The column names (e.g., 'Cell Line') will appear as labels.
row_colors_df = pd.DataFrame(
    {"Cell Line": mapped_cell_colors, "Modification": mapped_mod_colors}
)
# Important: The index must align with the data being plotted
row_colors_df.index = data_by_family.index


# --- 2. Plot the Clustermap ---
data_to_plot = data_by_family[family_names].T
data_to_plot

g = sns.clustermap(
    data_to_plot,
    cmap="coolwarm",
    # Goal 1: Customize figsize
    figsize=(11, 13),
    # Goal 2 & 3: Add row colors using the mapped palettes
    # We use row_colors, as 'cell_line' and 'modification' describe the rows
    col_colors=row_colors_df[["Modification", "Cell Line"]],
    row_colors=mapped_families_colors,
    # Other aesthetic improvements
    # vmin=-1, vmax=1,          # Set colorbar limits for correlation
    linewidths=0.5,
    cbar_kws={"label": "Average signal"},
    # metric='correlation'      # Use correlation for clustering distance
)

# --- 3. Save the Figure ---
save_path = "./plots/signal_clustermap_families.svg"

# Goal 4: Save as SVG
g.savefig(save_path, format="svg")

plt.show()

# %% [code]
data_by_family = pd.read_csv("SD_signal_by_lines_mods_TE_families.csv", index_col=None)
family_names = list(data_by_family.columns[:-2])
data_by_family[family_names] = data_by_family[family_names].apply(
    pd.to_numeric, errors="coerce"
)

data_by_family.index = data_by_family.cell_line + " " + data_by_family.modification
data_by_family

mapped_cell_colors = data_by_family["cell_line"].map(cell_line_p)
mapped_mod_colors = data_by_family["modification"].map(modification_p_richer)

families_by_classes = pd.read_csv("families_by_classes_TE.csv").dropna()

families_by_classes = families_by_classes.set_index("family_name").reindex(family_names)
mapped_families_colors = families_by_classes["class_name"].map(class_names_p)

# Create a DataFrame for the row_colors argument
# The column names (e.g., 'Cell Line') will appear as labels.
row_colors_df = pd.DataFrame(
    {"Cell Line": mapped_cell_colors, "Modification": mapped_mod_colors}
)
# Important: The index must align with the data being plotted
row_colors_df.index = data_by_family.index


# --- 2. Plot the Clustermap ---
data_to_plot = data_by_family[family_names].T
data_to_plot

g = sns.clustermap(
    data_to_plot,
    cmap="coolwarm",
    # Goal 1: Customize figsize
    figsize=(11, 13),
    # Goal 2 & 3: Add row colors using the mapped palettes
    # We use row_colors, as 'cell_line' and 'modification' describe the rows
    col_colors=row_colors_df[["Modification", "Cell Line"]],
    row_colors=mapped_families_colors,
    # Other aesthetic improvements
    # vmin=-1, vmax=1,          # Set colorbar limits for correlation
    linewidths=0.5,
    cbar_kws={"label": "Average signal"},
    # metric='correlation'      # Use correlation for clustering distance
)

# --- 3. Save the Figure ---
save_path = "./plots/signal_clustermap_families.svg"

# Goal 4: Save as SVG
g.savefig(save_path, format="svg")

plt.show()

# %% [code]
data_by_class = pd.read_csv(
    "Average_signal_by_lines_mods_TE_classes.csv", index_col=None
)
class_names = list(data_by_class.columns[:-2])
data_by_class[class_names] = data_by_class[class_names].apply(
    pd.to_numeric, errors="coerce"
)

data_by_class.index = data_by_class.cell_line + " " + data_by_class.modification

mapped_cell_colors = data_by_class["cell_line"].map(cell_line_p)
mapped_mod_colors = data_by_class["modification"].map(modification_p_richer)

# --- 2. Plot the Clustermap ---
data_to_plot = data_by_class[class_names].T
data_to_plot

g = sns.clustermap(
    data_to_plot,
    cmap="coolwarm",
    figsize=(11, 5),
    col_colors=row_colors_df[["Modification", "Cell Line"]],
    linewidths=0.5,
    cbar_kws={"label": "Average signal"},
)

plt.show()

data_by_class = pd.read_csv("SD_signal_by_lines_mods_TE_classes.csv", index_col=None)
class_names = list(data_by_class.columns[:-2])
data_by_class[class_names] = data_by_class[class_names].apply(
    pd.to_numeric, errors="coerce"
)

data_by_class.index = data_by_class.cell_line + " " + data_by_class.modification

mapped_cell_colors = data_by_class["cell_line"].map(cell_line_p)
mapped_mod_colors = data_by_class["modification"].map(modification_p_richer)

# --- 2. Plot the Clustermap ---
data_to_plot = data_by_class[class_names].T
data_to_plot

g = sns.clustermap(
    data_to_plot,
    cmap="coolwarm",
    figsize=(11, 5),
    col_colors=row_colors_df[["Modification", "Cell Line"]],
    linewidths=0.5,
    cbar_kws={"label": "Average signal"},
)

plt.show()

# %% [markdown]
## Analyze correlation coefficients of individual copies of TEs

# %% [markdown]
### Connection of correlation with average divergence and count by individual

# %% [code]
correlation_df_individuals = pd.read_csv(
    "Signal_vs_divergence_correlation_by_lines_mods_individuals_detailed.csv", index_col=0
)
correlation_df_individuals['abs_r'] = correlation_df_individuals.r.abs()
corrs_average = correlation_df_individuals[['cell_line', 'modification', 'individual_name', 'abs_r']].groupby(['individual_name']).mean()

correlation_df_individuals_random = pd.read_csv(
    "Signal_vs_divergence_correlation_by_lines_mods_individuals_detailed_random.csv", index_col=0
)
correlation_df_individuals_random['abs_r'] = correlation_df_individuals_random.r.abs()
corrs_average_random = correlation_df_individuals_random[['cell_line', 'modification', 'individual_name', 'abs_r']].groupby(['individual_name']).mean()


repeat_masker = pd.read_csv("T2T_repeat_masker_processed.csv")
divergence_average = (
    repeat_masker[["score", "individual_name"]]
    .groupby("individual_name")
    .mean()["score"]
    .reindex(corrs_average.index)
)

individuals_counts = pd.read_csv("repeat_masker_individual_name_counts.csv")
individuals_counts.columns = ["individual_name", "count"]
individuals_counts["log10_count"] = np.log10(individuals_counts["count"])
individuals_counts = individuals_counts.set_index("individual_name").reindex(corrs_average.index)


individuals_by_classes = pd.read_csv("individuals_by_classes_TE.csv").dropna()


individuals_by_classes = individuals_by_classes.set_index("individual_name").reindex(
    corrs_average.index
)

data_to_plot = pd.concat(
    [
        corrs_average,
        corrs_average_random,
        divergence_average,
        individuals_counts[["count", "log10_count"]],
        individuals_by_classes["class_name"],
    ],
    axis=1,
)
data_to_plot.columns = [
    "average_abs_corr",
    "average_abs_corr_random",
    "average_divergence",
    "count",
    "log10_count",
    "class_name",
]
data_to_plot

# %% [code]
individuals_by_classes = pd.read_csv("individuals_by_classes_TE.csv").dropna()
individuals_by_classes[individuals_by_classes.class_name == 'Retroposon']

# %% [code]


# %% [code]
sns.pairplot(
    data=data_to_plot[
        [
            "average_abs_corr",
            "average_divergence",
            "log10_count",
            "class_name",
        ]
    ],
    hue="class_name"
    
)

# %% [code]
import matplotlib.pyplot as plt
import numpy as np
import pandas as pd
import seaborn as sns
from scipy import stats
from scipy.optimize import curve_fit

plt.rcParams["svg.fonttype"] = "none"  # For clean SVG text


# --- 2. Define the sigmoidal function ---
def sigmoid_decay(x, L, k, x0):
    """
    Sigmoidal decay function (inverse logistic).
    L: maximum value (the asymptote)
    k: steepness of the curve
    x0: the x-value of the sigmoid's midpoint
    """
    # Note: The sign on k is positive for a decay curve
    return L / (1 + np.exp(k * (x - x0)))


# --- 3. Prepare data and perform the fit ---
print("Fitting sigmoidal curve...")

# Drop NaNs to ensure alignment and prevent errors
fit_data = data_to_plot.dropna() #[["log10_count", "average_abs_corr", "average_abs_corr_random"]]
x_data = fit_data["log10_count"]
y_data = fit_data["average_abs_corr"]
y_data_random = fit_data["average_abs_corr_random"]

# Provide initial guesses for the parameters [L, k, x0]
# L: guess the max value of y
# k: guess 1 (steepness)
# x0: guess the median value of x
p0 = [y_data.max(), 1, x_data.median()]
p0_random = [y_data_random.max(), 1, x_data.median()]
try:
    # Fit the curve
    popt, pcov = curve_fit(sigmoid_decay, x_data, y_data, p0=p0, maxfev=5000)
    L, k, x0 = popt
    popt_random, pcov_random = curve_fit(sigmoid_decay, x_data, y_data_random, p0=p0_random, maxfev=5000)
    L_random, k_random, x0_random = popt_random
    
    fit_successful = True
    y_pred = sigmoid_decay(x_data, *popt)
    corr, p_value = stats.pearsonr(y_data, y_pred)
    r_squared = corr**2
    if p_value < 0.001:
        p_val_str = "p < 0.001"
    else:
        p_val_str = f"p = {p_value:.3f}"
    equation_text = (
        f"y = {L:.2f} / (1 + exp(-{k:.2f} $\\cdot$ (x - {x0:.2f})))\n"
        f"$R^2$ = {r_squared:.3f}\n"
        f"{p_val_str}"
    )

        # --- 5. NEW: Calculate 95% Upper Limit ---
    # Calculate relative residuals: (actual - predicted) / predicted
    # This handles the changing variance (heteroscedasticity)
    y_pred_random = sigmoid_decay(x_data, *popt_random)
    relative_residuals = (y_data_random - y_pred_random) / y_pred_random

    # Find the 95th percentile of these relative residuals
    rel_resid_95th = np.percentile(relative_residuals, 95)
    
    print(f"95th percentile of relative error: +{rel_resid_95th * 100:.2f}%")


except RuntimeError:
    # This happens if the curve fit fails to converge
    print("Sigmoidal curve fit failed to converge.")
    fit_successful = False
    equation_text = "Fit failed to converge"


# --- 5. Create the Plot ---
print("Generating plot...")
# Create the figure and axes (for figsize customization)
fig, ax = plt.subplots(figsize=(10, 5))

# Draw the scatterplot onto the axis
sns.scatterplot(
    data=data_to_plot,
    hue="class_name",
    x="log10_count",
    y="average_abs_corr",
    palette=class_names_p,
    ax=ax,
    alpha=0.5,
)

if fit_successful:

    x_fit = np.linspace(x_data.min(), x_data.max(), 200)
    y_fit = sigmoid_decay(x_fit, *popt)
    ax.plot(x_fit, y_fit, color="black", ls="--", lw=2.5, label="Sigmoid Fit")

    y_fit_random = sigmoid_decay(x_fit, *popt_random)    
    ax.plot(x_fit, y_fit_random, color="red", ls="--", lw=2.5, label="Sigmoid Fit Random")
    # Plot the new 95% upper limit curve
    y_upper_limit_random = y_fit_random * (1 + rel_resid_95th)

    ax.plot(x_fit, y_upper_limit_random, color="red", ls=":", lw=2.5, label="95% Upper Limit")

    fit_data['average_abs_corr_random_95_top'] = sigmoid_decay(fit_data['log10_count'], *popt_random) * (1 + rel_resid_95th)
    
    # --- 7. Add the Equation Textbox ---
    ax.text(
        0.05,
        0.95,  # Position in (x, y) coordinates (0.05=5% from left, 0.95=95% from top)
        equation_text,
        transform=ax.transAxes,  # Use axis-relative coordinates
        fontsize=10,
        verticalalignment="top",
        ha="left",
        # Add a light background box
        bbox=dict(facecolor="white", alpha=0.7, boxstyle="round,pad=0.5"),
    )

# --- 8. Tidy Up and Save ---
ax.legend(loc="center left", bbox_to_anchor=(1, 0.5))
ax.set_title("Sigmoidal Fit of Correlation vs. Count", fontsize=16)
ax.set_xlabel("Count", fontsize=12)
ax.set_ylabel("Average Absolute Correlation", fontsize=12)
#ax.set_ylim((0, 0.225))

plt.tight_layout(rect=[0, 0, 0.85, 1])  # Make space for the legend

# Save to SVG
save_path = "./plots/abs_average_corr_vs_count_TE_individuals.svg"
fig.savefig(save_path, format="svg", transparent=True)
print(f"Plot saved to {save_path}")

plt.show()

# %% [code]
fit_data.sort_values('average_abs_corr')

# %% [code]
fit_data['status_95'] = 'above'
fit_data['status_95'][fit_data['average_abs_corr'] < fit_data['average_abs_corr_random_95_top']] = 'below'
fig, ax = plt.subplots(figsize=(10, 5))
sns.scatterplot(
    data=fit_data,
    hue="status_95",
    x="log10_count",
    y="average_abs_corr",

    ax=ax,
    alpha=0.5,
)


# %% [code]
fit_data.to_csv('average_corrs_divergence_random_status_individuals.csv')

# %% [code]
fig, ax = plt.subplots(figsize=(7, 5))
sns.scatterplot(
    data=data_to_plot,
    hue="class_name",
    x="average_divergence",
    y="average_abs_corr",
    palette=class_names_p,
    ax=ax,
    alpha=0.5,
)
save_path = "./plots/abs_average_corr_vs_divergence_TE_individuals.svg"
fig.savefig(save_path, format="svg", transparent=True)

plt.show()

# %% [markdown]
#### Random correlations

# %% [code]
correlation_df_individuals = pd.read_csv(
    "Signal_vs_divergence_correlation_by_lines_mods_individuals_detailed_random.csv", index_col=0
)
correlation_df_individuals['abs_r'] = correlation_df_individuals.r.abs()
corrs_average = correlation_df_individuals[['cell_line', 'modification', 'individual_name', 'abs_r']].groupby(['individual_name']).mean()

repeat_masker = pd.read_csv("T2T_repeat_masker_processed.csv")
divergence_average = (
    repeat_masker[["score", "individual_name"]]
    .groupby("individual_name")
    .mean()["score"]
    .reindex(corrs_average.index)
)

individuals_counts = pd.read_csv("repeat_masker_individual_name_counts.csv")
individuals_counts.columns = ["individual_name", "count"]
individuals_counts["log10_count"] = np.log10(individuals_counts["count"])
individuals_counts = individuals_counts.set_index("individual_name").reindex(corrs_average.index)


individuals_by_classes = pd.read_csv("individuals_by_classes_TE.csv").dropna()


individuals_by_classes = individuals_by_classes.set_index("individual_name").reindex(
    corrs_average.index
)

data_to_plot = pd.concat(
    [
        corrs_average,
        divergence_average,
        individuals_counts[["count", "log10_count"]],
        individuals_by_classes["class_name"],
    ],
    axis=1,
)
data_to_plot.columns = [
    "average_abs_corr",
    "average_divergence",
    "count",
    "log10_count",
    "class_name",
]

data_to_plot

# %% [code]
import matplotlib.pyplot as plt
import numpy as np
import pandas as pd
import seaborn as sns
from scipy import stats
from scipy.optimize import curve_fit

plt.rcParams["svg.fonttype"] = "none"  # For clean SVG text


# --- 2. Define the *CORRECTED* sigmoidal decay function ---
def sigmoid_decay(x, L, k, x0):
    """
    Sigmoidal decay function (inverse logistic).
    L: maximum value (the asymptote)
    k: steepness of the curve
    x0: the x-value of the sigmoid's midpoint
    """
    # Note: The sign on k is positive for a decay curve
    return L / (1 + np.exp(k * (x - x0)))


# --- 3. Prepare data and perform the fit ---
print("Fitting sigmoidal curve...")

fit_data = data_to_plot[["log10_count", "average_abs_corr"]].dropna()
x_data = fit_data["log10_count"]
y_data = fit_data["average_abs_corr"]

# Provide initial guesses [L, k, x0] for the *decay* curve
p0 = [y_data.max(), 1.0, x_data.median()]

try:
    popt, pcov = curve_fit(sigmoid_decay, x_data, y_data, p0=p0, maxfev=5000)
    L, k, x0 = popt
    popt_random = list(popt)
    fit_successful = True

    # --- 4. Calculate Statistics (R-squared and p-value) ---
    y_pred = sigmoid_decay(x_data, *popt)
    corr, p_value = stats.pearsonr(y_data, y_pred)
    r_squared = corr**2

    if p_value < 0.001:
        p_val_str = "p < 0.001"
    else:
        p_val_str = f"p = {p_value:.3f}"

    # Create the *corrected* equation string for the plot
    equation_text = (
        f"y = {L:.2f} / (1 + exp({k:.2f} $\\cdot$ (x - {x0:.2f})))\n"
        f"$R^2$ = {r_squared:.3f}\n"
        f"{p_val_str}"
    )

    # --- 5. NEW: Calculate 95% Upper Limit ---
    
    # Calculate relative residuals: (actual - predicted) / predicted
    # This handles the changing variance (heteroscedasticity)
    relative_residuals = (y_data - y_pred) / y_pred

    # Find the 95th percentile of these relative residuals
    rel_resid_95th = np.percentile(relative_residuals, 95)
    
    print(f"95th percentile of relative error: +{rel_resid_95th * 100:.2f}%")

except RuntimeError:
    print("Sigmoidal curve fit failed to converge.")
    fit_successful = False
    equation_text = "Fit failed to converge"


# --- 6. Create the Plot ---
print("Generating plot...")
fig, ax = plt.subplots(figsize=(7, 4))

sns.scatterplot(
    data=data_to_plot,
    hue="class_name",
    x="log10_count",
    y="average_abs_corr",
    palette=class_names_p,
    ax=ax,
    alpha=0.3,
)

if fit_successful:
    # --- 7. Plot the Mean Fit and 95% Upper Limit ---
    x_fit = np.linspace(x_data.min(), x_data.max(), 200)
    y_fit_random = sigmoid_decay(x_fit, *popt)

    # Calculate the upper limit curve using the relative residual
    y_upper_limit = y_fit * (1 + rel_resid_95th)

    # Plot the mean curve
    ax.plot(x_fit, y_fit_random, color="black", ls="--", lw=2.5, label="Mean Fit (Control)")
    x_fit_upper = np.array(x_fit)
    # Plot the new 95% upper limit curve
    ax.plot(x_fit, y_upper_limit, color="red", ls=":", lw=2.5, label="95% Upper Limit")
    

    # --- 8. Add the Equation Textbox ---
    ax.text(
        0.05, 0.95,
        equation_text,
        transform=ax.transAxes,
        fontsize=10,
        verticalalignment='top',
        ha='left',
        bbox=dict(facecolor="white", alpha=0.7, boxstyle="round,pad=0.5"),
    )

# --- 9. Tidy Up and Save ---
ax.legend(loc="center left", bbox_to_anchor=(1, 0.5))
ax.set_title("Sigmoidal Fit of Correlation vs. Count", fontsize=16)
ax.set_xlabel("log10(Count)", fontsize=12) # Changed label
ax.set_ylabel("Average Absolute Correlation", fontsize=12)
ax.set_ylim((0, 0.225))

plt.tight_layout(rect=[0, 0, 0.85, 1])

save_path = "./plots/abs_average_corr_vs_count_TE_individuals_random.svg"
fig.savefig(save_path, format="svg")
print(f"Plot saved to {save_path}")

plt.show()

# %% [markdown]
### Visualize top and bottom correlations for individuals, marks and cell lines

# %% [code]
correlation_df_individuals

# %% [code]
correlation_df_individuals = pd.read_csv(
    "Signal_vs_divergence_correlation_by_lines_mods_individuals_detailed.csv", index_col=0
).dropna()

average_corrs_random_check = pd.read_csv(
    "average_corrs_divergence_random_status_individuals.csv", index_col=0
).dropna()
from statsmodels.stats.multitest import multipletests

reject, p_adjusted, _, _ = multipletests(
    correlation_df_individuals["p_value"], alpha=0.05, method="fdr_bh"
)

correlation_df_individuals["q_value"] = p_adjusted
top_corrs = pd.concat(
    [
        correlation_df_individuals[correlation_df_individuals["ci_low"] > 0.4],
        correlation_df_individuals[correlation_df_individuals["ci_high"] < -0.4],
    ]
)
top_corrs=top_corrs[top_corrs['individual_name'].isin(average_corrs_random_check.index[average_corrs_random_check['status_95'] == 'above'])]
top_corrs.reset_index()

# %% [code]
reject, p_adjusted, _, _ = multipletests(
    correlation_df_individuals["p_value"].dropna(), alpha=0.05, method="fdr_bh"
)
p_adjusted

# %% [code]


# %% [code]
p_adjusted

# %% [code]


save_path_svg = f"plots/Divergence_vs_Signal_by_individuals_sign_correlations.svg"

figure = plt.figure(figsize=(10, 5))

N_ROWS = 3
N_COLS = 4

# Increased spacing for independent labels
outer_grid = GridSpec(N_ROWS, N_COLS, figure=figure, wspace=0.4, hspace=0.5)

# --- Removed shared axes lists ---

# --- Add a cache for data loading ---
data_cache = {}

# --- Use a more Pythonic loop ---
for i, (idx, row) in enumerate(
    tqdm(top_corrs.iterrows(), total=len(top_corrs), desc="Plotting Top")
):

    # Calculate grid position
    r = i // N_COLS
    c = i % N_COLS

    # Get data from the row
    individual_name = row["individual_name"]
    cell_line = row["cell_line"]
    modification = row["modification"]

    filepath = f"./epigenomic_files/{cell_line}.{modification}.chm13v2.0.mapped_on_repeat_masker.bedGraph"

    # --- Check cache before loading ---
    if filepath in data_cache:
        tqdm.write(f"  -> Found in cache: {filepath}")
        sample_enrichment = data_cache[filepath]
    else:
        # --- Data Loading ---
        tqdm.write(f"  -> Loading data from: {filepath}")
        try:
            sample_enrichment = pd.read_csv(
                filepath,
                sep="\t",
                header=None,
            )
            data_cache[filepath] = sample_enrichment
            tqdm.write("File loaded and cached.")

        except FileNotFoundError:
            tqdm.write(f"File not found, skipping: {filepath}")
            ax = figure.add_subplot(outer_grid[r, c])
            ax.axis("off")
            continue
        except Exception as e:
            tqdm.write(
                    f"An unexpected error occurred while reading {filepath}: {e}"
            )
            ax = figure.add_subplot(outer_grid[r, c])
            ax.axis("off")
            continue

        # --- Process data (only needs to be done once, on load) ---
        sample_enrichment.columns = [
            "Chromosome",
            "Start",
            "End",
            "Divergence",
            "Name",
            "Family",
            "Class",
            "Signal",
        ]
        sample_enrichment["Signal"] = pd.to_numeric(
            sample_enrichment["Signal"], errors="coerce"
        )
        sample_enrichment["Divergence"] = pd.to_numeric(
            sample_enrichment["Divergence"], errors="coerce"
        )
        data_cache[filepath] = sample_enrichment

    # --- Query must be done every time ---
    query_local = f'Name == "{individual_name}"'
    sample_enrichment_local = sample_enrichment.query(query_local)

    # --- Create *single* axis for this plot ---
    ax_main = figure.add_subplot(outer_grid[r, c])

    # --- Plot the data ---
    sns.histplot(
        data=sample_enrichment_local,
        x="Divergence",
        y="Signal",
        bins=48,
        pmax=0.9,
        cmap="viridis",
        ax=ax_main,
    )

    # --- Tidy up: Main plot ticks/labels ---
    ax_main.set_facecolor("black")
    ax_main.set_xlabel("Divergence", fontsize=10)
    ax_main.set_ylabel(f"{cell_line}\n{modification}", fontsize=10)

    # Set spine colors to white

    corr_coef, p_val, ci_low, ci_high = row["r"], row["p_value"], row["ci_low"], row["ci_high"]


    corr_str = f"r = {corr_coef:.3f} [{ci_low:.3f}, {ci_high:.3f}]"

    ax_main.set_title(f"{individual_name}\n{corr_str}", fontsize=10, pad=10)


figure.tight_layout()  # Add tight_layout to prevent label overlap
figure.savefig(save_path_svg, format="svg", facecolor="white", transparent=False)
plt.show()

print(f"Plot saved to {save_path_svg}")

# %% [code]
sample_enrichment

# %% [code]


# %% [markdown]
### Clustermap of correlation coefficients

# %% [code]
correlation_df_individuals = pd.read_csv(
    "Signal_vs_divergence_correlation_by_lines_mods_individuals_detailed.csv", index_col=0
)
correlation_df_individuals = pd.pivot_table(correlation_df_individuals, values='r', index=['cell_line', 'modification'],
                       columns=['individual_name'], aggfunc="sum", fill_value=0).reset_index()
correlation_df_individuals.index = (
    correlation_df_individuals.cell_line + " " + correlation_df_individuals.modification
)
correlation_df_individuals

# %% [code]
correlation_df_individuals[correlation_df_individuals.columns[correlation_df_individuals.columns.str.startswith('AluY')]]


# %% [code]
individuals_counts = pd.read_csv("repeat_masker_individual_name_counts.csv")
individuals_counts.columns = ["individual_name", "count"]

individual_names = individuals_counts['individual_name'][:500]
individual_names=correlation_df_individuals.columns[correlation_df_individuals.columns.str.startswith('AluY')]
individual_names = ['LTR12F', 'LTR5_Hs', 'HERVIP10FH-int', 'LTR13_v']
individuals_by_classes = pd.read_csv("individuals_by_classes_TE.csv").dropna()

individuals_by_classes = individuals_by_classes.set_index("individual_name").reindex(
    individual_names
)


individuals_by_classes

# %% [code]
len(individual_names)

# %% [code]
plt.rcParams.update({'font.size': 16})

# --- 1. Prepare Row Colors ---
# Map the 'cell_line' and 'modification' columns to their respective colors
mapped_cell_colors = correlation_df_individuals["cell_line"].map(cell_line_p)
mapped_mod_colors = correlation_df_individuals["modification"].map(
    modification_p_richer
)

individuals_by_classes = pd.read_csv("individuals_by_classes_TE.csv").dropna()

individuals_by_classes = individuals_by_classes.set_index("individual_name").reindex(
    individual_names
)
mapped_individuals_colors = individuals_by_classes["class_name"].map(class_names_p)

# Create a DataFrame for the row_colors argument
# The column names (e.g., 'Cell Line') will appear as labels.
row_colors_df = pd.DataFrame(
    {"Cell Line": mapped_cell_colors, "Modification": mapped_mod_colors}
)
# Important: The index must align with the data being plotted
row_colors_df.index = correlation_df_individuals.index


# --- 2. Plot the Clustermap ---
data_to_plot = correlation_df_individuals[individual_names].T


g = sns.clustermap(
    data_to_plot.T,
    cmap="coolwarm",
    # Goal 1: Customize figsize
    figsize=(15, 15),
    # Goal 2 & 3: Add row colors using the mapped palettes
    # We use row_colors, as 'cell_line' and 'modification' describe the rows
    row_colors=row_colors_df[["Modification", "Cell Line"]],
    col_colors=mapped_individuals_colors,
    # Other aesthetic improvements
    # vmin=-1, vmax=1,          # Set colorbar limits for correlation
    linewidths=0.5,
    cbar_kws={"label": "Pearson Correlation"},
    #xticklabels=False, yticklabels=False
    # metric='correlation'      # Use correlation for clustering distance
)
#g.ax_col_dendrogram.set_visible(False)

# --- 3. Save the Figure ---
save_path = "./plots/signal_to_divergence_corr_clustermap_individuals_AluY.svg"
save_path_png = "./plots/signal_to_divergence_corr_clustermap_individuals_AluY.png"

# Goal 4: Save as SVG
g.savefig(save_path, format="svg")
g.savefig(save_path_png, format="png")

plt.show()

# %% [markdown]
#### Permutation test

# %% [code]


# Calculate pairwise Euclidean distances
# Ensure you select only the numerical columns for distance calculation
distances = pdist(data_matrix, metric='euclidean')
# Convert the condensed distance matrix to a squareform matrix
distance_matrix_raw = squareform(distances)
pd.DataFrame(distance_matrix)

# %% [code]
distance_matrix_raw

# %% [code]
DistanceMatrix(PairwiseMatrix(distance_matrix_raw))

# %% [code]
skbio.stats.distance.SymmetricMatrix

# %% [code]
import os
import numpy as np
import pandas as pd
from tqdm import tqdm
from scipy import stats
from scipy.spatial.distance import pdist, squareform

# --- Imports for PERMANOVA Test ---
# You must install scikit-bio: pip install scikit-bio
try:
    from skbio.stats.distance import DistanceMatrix, permanova, SymmetricMatrix, PairwiseMatrix
    from sklearn.metrics import pairwise_distances
except ImportError:
    print("Error: scikit-bio or sklearn not found.")
    print("Please run: pip install scikit-bio scikit-learn")
    exit()

# --- Helper Class for Colored Logging ---
class Logger:
    """A simple logger for pretty, colored console output."""
    BLUE = "\033[94m"
    GREEN = "\033[92m"
    YELLOW = "\033[93m"
    ENDC = "\033[0m"
    @staticmethod
    def info(message): print(f"{Logger.BLUE}INFO: {message}{Logger.ENDC}")
    @staticmethod
    def success(message): print(f"{Logger.GREEN}SUCCESS: {message}{Logger.ENDC}")
    @staticmethod
    def warning(message): print(f"{Logger.YELLOW}WARNING: {message}{Logger.ENDC}")

# --- 1. Load the Data ---
# (This logic is copied from your plotting script)
Logger.info("Loading correlation data...")
try:
    # We need the 'cell_line' and 'modification' columns, so we don't use .T
    # We also need the original index, so don't set index_col=0
    correlation_df_individuals = pd.read_csv(
        "Signal_vs_divergence_correlation_by_lines_mods_individuals_detailed.csv", index_col=0
    )
    correlation_df_individuals = pd.pivot_table(correlation_df_individuals, values='r', index=['cell_line', 'modification'],
                           columns=['individual_name'], aggfunc="sum", fill_value=0).reset_index()
    correlation_df_individuals.index = (
        correlation_df_individuals.cell_line + " " + correlation_df_individuals.modification
    )
    # Get the list of TE names (all columns except the first two)
    individual_names = list(correlation_df_individuals.columns[2:])
    
except FileNotFoundError:
    Logger.warning("Error: Data file not found. This script must run after the correlation script.")
    exit()

# --- 2. Prepare Data for PERMANOVA ---
Logger.info("Preparing data for PERMANOVA test...")

data_matrix = correlation_df_individuals[individual_names]
grouping_df = correlation_df_individuals[["cell_line", "modification"]]

# --- 3. Calculate the Distance Matrix ---
# We use 'euclidean' as it's the default metric for sns.clustermap.
# This ensures we are testing the same clustering the user is seeing.
Logger.info("Calculating Euclidean distance matrix...")
distances = pdist(data_matrix, metric='euclidean')
# Convert the condensed distance matrix to a squareform matrix
distance_matrix_raw = squareform(distances)

# Convert to skbio's DistanceMatrix object, using the index
# (This is good practice, though not strictly required)
#dist_matrix = DistanceMatrix(dist_matrix_raw, ids=grouping_df.index)
dist_matrix=DistanceMatrix(PairwiseMatrix(distance_matrix_raw), ids=grouping_df.index)

# We use 9999 permutations for a robust p-value.
Logger.info("Running PERMANOVA (9999 permutations)...")

# Test 'Modification' alone
results_mod = permanova(
    dist_matrix, 
    grouping_df['modification'], 
    permutations=1000000
)

# Test 'Cell Line' alone
results_cell = permanova(
    dist_matrix, 
    grouping_df['cell_line'], 
    permutations=1000000
)


# --- 5. Print the Results ---
Logger.success("\n--- PERMANOVA Test Results ---")

print("\n--- Test 1: Clustering by 'Modification' ---")
print(f"  R-squared: {results_mod['R^2']:.4f}")
print(f"  p-value:   {results_mod['p-value']:.4f}")
if results_mod['p-value'] < 0.05:
    print("  RESULT: Clustering by 'Modification' is STATISTICALLY SIGNIFICANT.")
else:
    print("  RESULT: Clustering by 'Modification' is NOT significant.")

print("\n--- Test 2: Clustering by 'Cell Line' ---")
print(f"  R-squared: {results_cell['R^2']:.4f}")
print(f"  p-value:   {results_cell['p-value']:.4f}")
if results_cell['p-value'] < 0.05:
    print("  RESULT: Clustering by 'Cell Line' is STATISTICALLY SIGNIFICANT.")
else:
    print("  RESULT: Clustering by 'Cell Line' is NOT significant.")


print("\n--- Interpretation ---")
print("R-squared (R^2) is the effect size. It shows how much of the variation")
print("in the distance matrix is 'explained' by the grouping variable(s).")
print("A high R^2 for 'Cell Line' and a low R^2 for 'Modification'")
print("would suggest that the primary clustering is driven by cell line, not modification.")

# %% [code]
results_mod

# %% [code]
results_cell

# %% [code]
r2 = 1 - 1 / (1 + results_mod[4] * results_mod[3] / (results_mod[2] - results_mod[3] - 1))
r2

# %% [code]
r2 = 1 - 1 / (1 + results_cell[4] * results_cell[3] / (results_cell[2] - results_cell[3] - 1))
r2

# %% [code]
np.allclose(distance_matrix_raw, distance_matrix_raw.T, rtol=1e-05, atol=1e-08)

# %% [code]
pd.DataFrame(distance_matrix_raw).dropna()

# %% [code]


# %% [markdown]
### Clustermaps by TE class

# %% [code]


individuals_by_classes

# %% [code]
class_name = 'DNA'

individuals_by_classes = pd.read_csv("individuals_by_classes_TE.csv").dropna()

individuals_by_classes = individuals_by_classes.set_index("individual_name")
individual_names = individuals_by_classes.index[individuals_by_classes['class_name'] == class_name]


plt.rcParams.update({'font.size': 16})

# --- 1. Prepare Row Colors ---
# Map the 'cell_line' and 'modification' columns to their respective colors
mapped_cell_colors = correlation_df_individuals["cell_line"].map(cell_line_p)
mapped_mod_colors = correlation_df_individuals["modification"].map(
    modification_p_richer
)

individuals_by_classes = pd.read_csv("individuals_by_classes_TE.csv").dropna()

individuals_by_classes = individuals_by_classes.set_index("individual_name").reindex(
    individual_names
)
mapped_individuals_colors = individuals_by_classes["class_name"].map(class_names_p)

# Create a DataFrame for the row_colors argument
# The column names (e.g., 'Cell Line') will appear as labels.
row_colors_df = pd.DataFrame(
    {"Cell Line": mapped_cell_colors, "Modification": mapped_mod_colors}
)
# Important: The index must align with the data being plotted
row_colors_df.index = correlation_df_individuals.index


# --- 2. Plot the Clustermap ---
data_to_plot = correlation_df_individuals[individual_names].T


g = sns.clustermap(
    data_to_plot.T,
    cmap="coolwarm",
    # Goal 1: Customize figsize
    figsize=(80, 20),
    # Goal 2 & 3: Add row colors using the mapped palettes
    # We use row_colors, as 'cell_line' and 'modification' describe the rows
    row_colors=row_colors_df[["Modification", "Cell Line"]],
    col_colors=mapped_individuals_colors,
    # Other aesthetic improvements
    # vmin=-1, vmax=1,          # Set colorbar limits for correlation
    linewidths=0.5,
    cbar_kws={"label": "Pearson Correlation"},
    xticklabels=False, yticklabels=False
    # metric='correlation'      # Use correlation for clustering distance
)
g.ax_col_dendrogram.set_visible(False)

# --- 3. Save the Figure ---
save_path = f"./plots/signal_to_divergence_corr_clustermap_individuals_{class_name}.svg"
save_path_png = f"./plots/signal_to_divergence_corr_clustermap_individuals_{class_name}.png"

# Goal 4: Save as SVG
g.savefig(save_path, format="svg")
g.savefig(save_path_png, format="png")

plt.show()

# %% [code]


# %% [markdown]
### Curtosis etc

# %% [code]
correlation_df_individuals = pd.read_csv(
    "Signal_vs_divergence_correlation_by_lines_mods_individuals_detailed.csv", index_col=0
)
correlation_df_individuals["transformed_corr"] = np.arctanh(correlation_df_individuals.r)

correlation_df_individuals = correlation_df_individuals.dropna()
correlation_df_individuals

# %% [code]
correlation_df_individuals[correlation_df_individuals['r'].abs()>0.2]

# %% [code]
correlation_df_individuals[correlation_df_individuals['r'].abs()>0.4]

# %% [code]
sns.histplot(correlation_df_individuals.transformed_corr)

# %% [code]
stats.shapiro(correlation_df_individuals.transformed_corr)

# %% [code]
stats.kurtosistest(correlation_df_individuals.transformed_corr)

# %% [code]
stats.kurtosis(correlation_df_individuals.transformed_corr)

# %% [markdown]
### Levene test for variances comparison between multiple groups

# %% [code]
correlation_df_individuals = pd.read_csv(
    "Signal_vs_divergence_correlation_by_lines_mods_individuals_detailed.csv", index_col=0
)
correlation_df_individuals["transformed_corr"] = np.arctanh(correlation_df_individuals.r)

correlation_df_individuals = correlation_df_individuals.dropna()


individuals_by_classes = pd.read_csv("individuals_by_classes_TE.csv").dropna()
correlation_df_individuals = correlation_df_individuals.merge(individuals_by_classes)
correlation_df_individuals

# %% [code]


# %% [code]
from scipy.stats import levene

compare_list = [correlation_df_individuals["transformed_corr"][correlation_df_individuals["class_name"] == class_name] for class_name in correlation_df_individuals["class_name"].unique()] 

levene(*compare_list)

# %% [code]
from scipy.stats import levene

compare_list = [correlation_df_individuals["transformed_corr"][correlation_df_individuals["modification"] == modification] for modification in correlation_df_individuals["modification"].unique()] 

levene(*compare_list)

# %% [code]
from scipy.stats import levene

compare_list = [correlation_df_individuals["transformed_corr"][correlation_df_individuals["cell_line"] == cell_line] for cell_line in correlation_df_individuals["cell_line"].unique()] 

levene(*compare_list)

# %% [markdown]
# Epigenetic signals, GRE/NGRE - separate notebook and article


# %% [code]
```
